# Cross-species consensus atlas of the primate basal ganglia

**DOI:** 10.64898/2025.12.15.694496

**Authors:** Nelson J. Johansen, Yuanyuan Fu, Matthew Schmitz, Alma Dubuc, Niklas Kempynck, Morgan Wirthlin, Aaron D. Garcia, Madeleine Hewitt, Meghan A. Turner, Stephanie C. Seeman, Brian Long, Xiao-Ping Liu, Shu Dan, Michael DeBerardine, Inkar Kapen, Anna Marie Yanny, Alida Avola, Samuel T. Barlow, Darren Bertagnolli, Ashwin Bhandiwad, Agata Budzillo, Victor Eduardo Nieto Caballero, Lakme Caceres, Tamara Casper, Anish Bhaswanth Chakka, Rushil Chakrabarty, Michael Clark, Scott Daniel, Jeroen Eggermont, Rebecca Ferrer, Leon French, Jessica Gloe, Jeff Goldy, Nathan Guilford, Junitta Guzman, Daniel Hirschstein, Windy Ho, Katelyn James, Danielle Lynne Jones, Matthew Jungert, Madhav Kannan, Kasia Z. Kedzierska, Thomas Kroes, Mckaila Leytze, Arena Manning, Rachel McCue, Chris Morrison, Beagan Nguy, Sven Otto, Nick Pena, Trangthanh Pham, Elliot Phillips, Abhejit Rajagopal, Christine Rimorin, Andrea D. Rivera, Dana Rocha, Raymond Sanchez, Geoffrey Schau, Jessica Schembri, Nadiya V. Shapovalova, Jamie Sherman, Julian Thijssen, Michael Tieu, Amy Torkelson, Alex Tran, Alexander Vieth, Yongqi Wang, Dan Yuan, Jennie Close, Tanya L. Daigle, Rachel Dalley, Nick Dee, Song-Lin Ding, Elysha Fiabane, Nathan Gouwens, Grace Huynh, Brian R. Lee, Boaz P. Levi, Delissa McMillen, Jeremy A. Miller, Tyler Mollenkopf, Lydia Ng, Patrick L. Ray, Cassandra Sobieski, Staci A. Sorensen, Zizhen Yao, Faraz Yazdani, Winrich Freiwald, Boudewijn Lelieveldt, Brian Kalmbach, C. Dirk Keene, Jesse Gillis, David Osumi-Sutherland, Jonathan T. Ting, Hongkui Zeng, Kimberly Smith, Fenna M. Krienen, Rebecca D. Hodge, Ed S. Lein, Trygve E. Bakken

## Abstract

The basal ganglia (BG) are conserved brain regions essential for motor control, learning, emotion, and cognition, and are implicated in neurological and psychiatric disease. Yet a unified cross-species taxonomy of BG cell types is lacking, limiting translation of BG circuit mechanisms, interpretation of human genetic risk, and development of cell type-targeted tools. We present a multiomic consensus atlas of 1.8 million nuclei from human, macaque, and marmoset spanning eight BG structures. Integrating cross-species gene expression, open chromatin, and spatial profiling enables definition of conserved and divergent cell types. Alignment to existing mouse and human atlases identifies 61 homologous cell types conserved over 80 million years. We identify a STRd D2 StrioMat Hybrid medium spiny neuron (MSN) type with molecular, electrophysiological, and morphological features that clarify hybrid MSN identities. Comparative cis-regulatory analysis reveals conserved sequence grammars that encode cell identity and inform viral targeting strategies, providing a foundational resource for BG evolution, function, and disease.

## INTRODUCTION

Basal ganglia circuits are central to action selection, reinforcement learning, and habit formation, and their dysfunction underlies a broad spectrum of movement and psychiatric disorders ^1–3^. However, despite rapid growth of single-cell multiomic profiling, BG cell type definitions remain fragmented across regions, species, and modalities, and a unified cross-species molecular taxonomy that aligns homologous populations across structures and along the rostrocaudal axis is still lacking. In primates, GABAergic MSNs constitute the vast majority of projection neurons in caudate (Ca), putamen (Pu), and nucleus accumbens (NAC), and are classically organized into dopamine-modulated direct and indirect pathways, striosome and matrix compartments, and dorsal-ventral and limbic-sensorimotor territories ^4–8^. These striatal populations participate in parallel cortico-BG-thalamo-cortical loops via pallidal and nigral output nuclei. The projection-defined D1 and D2 MSNs have been functionally characterized using genetic and optogenetic tools ^4,9,10^. Decades of neuroanatomical and physiological work established D1- and D2-expressing MSNs as the cellular substrates of the direct and indirect pathways while linking striosome and matrix regional specializations to distinct behavioral functions and disease vulnerability ^1,2,11,12^. Beyond these dorsal striatal populations, ventral MSNs of the nucleus accumbens interface limbic cortico-BG circuits to regulate reward and motivation and are central to maladaptive plasticity in drug and alcohol use disorders ^13–16^.

This rich cellular heterogeneity has motivated modern cell type-level analyses of BG circuits. Large-scale brain atlases ^17–19^ and region- or cell type-focused comparative studies ^20–22^ show that cell type repertoires are broadly conserved across species but differ in abundance, spatial distribution, and molecular specializations, providing a framework for comparative analysis of BG circuits. Building on this foundation, single-cell and single-nucleus transcriptomic surveys in mouse, rat, and primate striatum (STR) have formalized and extended the diversity of both projection neurons and interneurons ^23–31^. These studies delineate discrete and continuous MSN cell types along with structured interneuron repertoires that align with and elaborate canonical pathway, compartment, and spatial dimensions. However, most prior genomics studies of BG have focused on restricted subsets of nuclei, a single primate species, or a single modality, limiting the development of unified cross-species taxonomies that jointly position primate MSNs and other BG cell types across nuclei and along the rostrocaudal axis ^27,31^. Within the BRAIN Initiative Cell Atlas Network (BICAN), the Human and Mammalian Brain Atlas (HMBA) project is charged with building cross-species reference atlases for mammalian brain cell types. Here, we generate a BG-focused single-nucleus multiomic atlas from >2 million nuclei collected in a coordinated manner from human, macaque, and marmoset Ca, Pu, NAC, ventral pallidum (VeP), the external and internal segments of globus pallidus (GPe, GPi), subthalamic nucleus (STH), and substantia nigra (SN) sampled densely along the rostrocaudal axis. We integrate these data with spatial transcriptomics (Hewitt, M.N. et al. co-submtted) ^32^, Patch-Seq (Liu et al., co-submitted)^33^, and whole-brain mouse transcriptomic resources ^17,18^. From these data, we construct a consensus cross-species BG taxonomy that emphasizes conserved marker genes and transcription factor (TF) grammars of cis-regulatory elements (cCREs) while defining a cell type nomenclature that retains established community names. This taxonomy encodes evolutionary relationships and functional distinctions among conserved MSNs, interneuron classes and projection neurons in GPe, GPi, STH, and SN. Within MSNs, our taxonomy organizes diversity along canonical pathway, compartment, and spatial axes and reveals a distinct STRd D2 StrioMat Hybrid MSN type with combined striosome-like, matrix-like and eccentric-like transcriptomic features, expanding on previously characterized hybrid populations observed in primate STR ^19,27^.

By defining human BG cell types within a consensus cross-species taxonomy, we link them to counterparts in model species, enabling inference of otherwise inaccessible human properties from experimental studies in animals. Using our human-centered taxonomy, we map common genetic risk for psychiatric and neurodegenerative disorders onto BG Groups, building on cell type enrichment analyses for interpreting brain genome-wide association studies (GWAS) ^34–37^. We find that polygenic risk for several psychiatric disorders is preferentially concentrated in ventral striatal MSNs, with additional enrichment in the STRd D2 StrioMat Hybrid MSN type for schizophrenia (SCZ). By working with high resolution, literature-informed cell types rather than broad transcriptomic classes, these analyses link genetic associations to established circuit and disease literature.

## RESULTS

### Consensus taxonomy of BG cell types for human, macaque, and marmoset

We built a cross-species consensus atlas of the BG through systematic dissections and high-throughput single-nucleus profiling in human, macaque, and marmoset. Regional dissections from 4-9 donors per species generated high-quality snRNA-seq and snATAC-seq datasets using the 10X Genomics multiome assay (Figure 1A). In marmoset, tiled dissections covered the entire BG and surrounding subcortical structures (described in Dan, Turner, DeBerardine et al., co-submitted)^38^, facilitating alignment of homologous populations across species. Single nuclei were enriched for neurons using NeuN-based sorting, yielding approximately the targeted 70% neuronal and 30% non-neuronal cells. After stringent quality control (Figure S1.1) and donor-effect correction, we applied iterative clustering workflows developed for recent whole-brain taxonomies^17^, producing well-resolved transcriptomic clusters for each species (Tables S1.1 and S1.2). These per-species clusters, derived from 1.8 million high quality nuclei across human, macaque, and marmoset, provide the foundation for a consensus, cross-species taxonomy of BG cell types.

**Figure 1.**
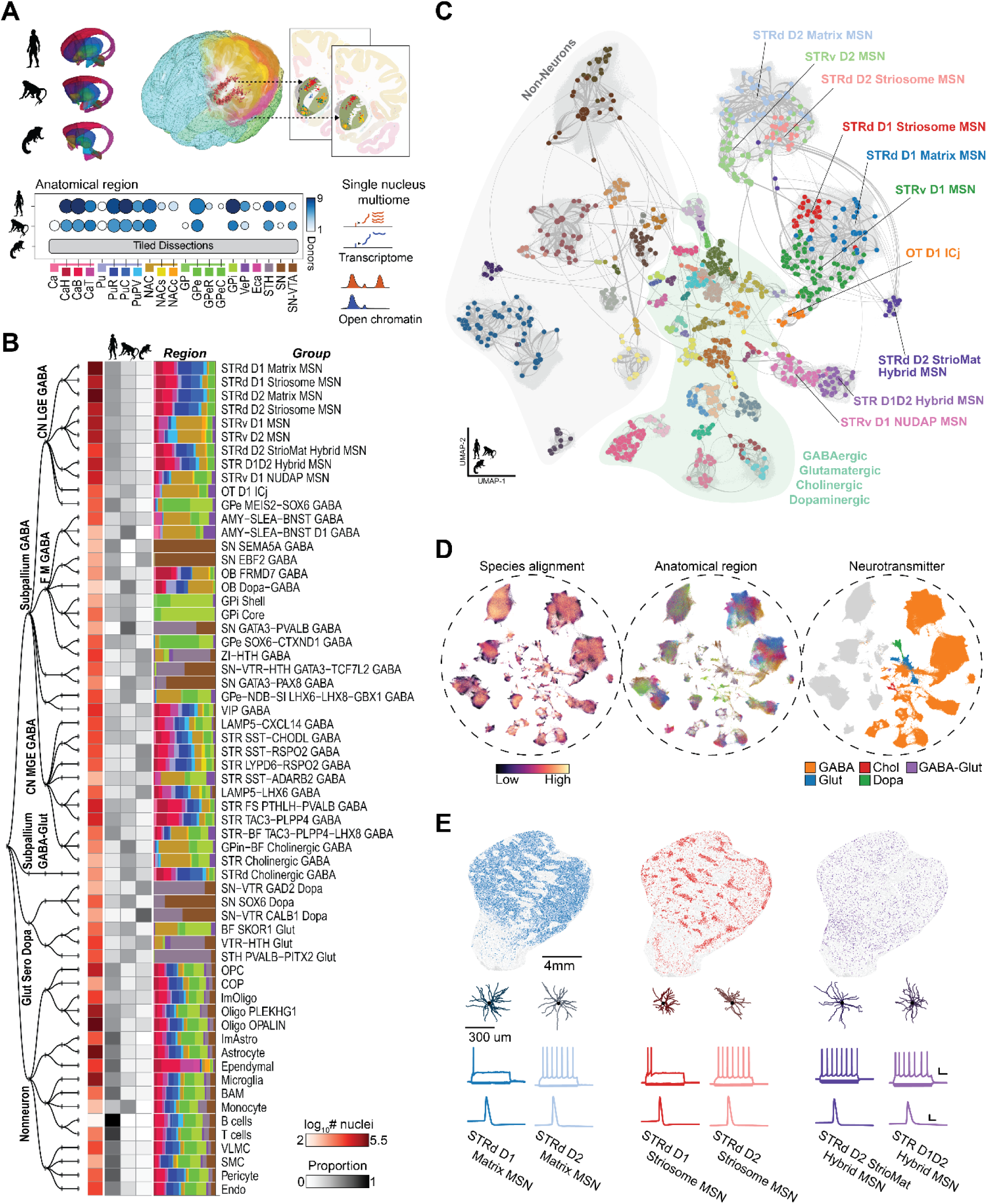
BG consensus cell type atlas across human, macaque, and marmoset. (**A**) Comprehensive single-nucleus multiomic profiling of BG regions across human, macaque, and marmoset. (**B**) Taxonomic hierarchy to the Group-level with heatmap annotation showing number of nuclei sampled, dissected region distribution. (**C**) UMAP visualization of the integrated scVI embedding of nuclei from all species overlaid with a constellation diagram of human clusters colored by cell groups. (**D**) UMAPs visualizing the species alignment, dissected region (legend in panel A), and inferred neurotransmitter. (**E**) Spatial transcriptomics, morphology, and electrophysiological properties of distinct MSN sub-populations in macaque.

We organized the transcriptomic clusters into a hierarchical cell type taxonomy based on conserved molecular, computational, and developmental evidence placing projection neurons, interneurons, and non-neuronal populations into a unified hierarchy starting from Neighborhoods (n=4), Classes (n=12), Subclasses (n=36), and Groups (n=61) informed by literature (Figure 1B, Figure S1.2). Group abundances were approximately conserved despite variation in sampling across species and exhibited both highly regional and distributed localization (Figure 1B, Figure S1.3). Despite denser anatomical sampling in humans, non-human primates contributed disproportionate recovery of rare cell types, enhancing transcriptomic characterization of low-abundance populations (Figure 1B). Annotation transfer from published human, non-human primate, and mouse references established an initial scaffold for major neuronal and non-neuronal types ^17,19,26–28,39–42^. In parallel, broad neurotransmitter-defined classes were validated using canonical marker genes and conserved TFs, ensuring consistent high-level divisions. At the coarsest level, Neighborhoods delineate Subpallium GABA, Subpallium GABA-Gluta, Gluta Sero Dopa, and Non-neuronal terms, consistent with whole-brain nomenclature (Yao et al. 2023; Fig. S1.3).

Progressive refinement at the Class level separates neurons by developmental origin (CN LGE GABA, CN MGE GABA, and CN CGE GABA; lateral, medial, and caudal ganglionic eminences, respectively), neurotransmitter identity (M Dopa), and conserved neurotransmitter and anatomical localization as confirmed by spatial transcriptomics (Hewitt et al. 2025) (F M GABA, Cx GABA, CN GABA-Glut, F M Glut) (Table S1.2). Subclass terms further integrate domain-specific nomenclature to resolve biologically coherent types within each Class; for example, CN LGE GABA subdivides into STR D1 MSN, STR D2 MSN, STR Hybrid, OT Granular GABA, and CN MEIS2 GABA. Furthermore, developmental lineage analyses demonstrated that embryonic origin closely mirrors adult structure, strengthening the cell type hierarchy and distinguishing classical from eccentric ^43^ MSN subclasses (Table S1.3).

Integrated cross-species analysis resolved a high-resolution set of 61 Group-level cell types underlying Subclasses and that were conserved across primates while revealing previously unappreciated diversity within major neuronal populations. A unified embedding of human, macaque, and marmoset transcriptomes demonstrated strong alignment of homologous cell types and clear separation of major neurotransmitter-defined populations, including dopaminergic, cholinergic, glutamatergic, and diverse GABAergic lineages (Figures 1C and,1D). Within MSNs, the predominant neuronal class of the BG, we consistently identified, across species, distinct Groups comprising matrix (STRd D1 Matrix MSN, STRd D2 Matrix MSN) ^7^, striosome (STRd D1 Striosome MSN, STRd D2 Striosome MSN) ^8^ and dorsal (STRd) and ventral (STRv D1 MSN, STRv D2 MSN) ^29^ (STRv) subregional distinctions, together with hybrid or transitional phenotypes (STR D1D2 Hybrid MSN, STRv D1 NUDAP MSN, STRd D2 StrioMat Hybrid MSN). We use ‘Group’ to denote transcriptomically defined MSN populations that are striosome-enriched or matrix-enriched by molecular signature and spatial mapping, rather than implying a strictly bounded histological compartment.

Cross-species MetaNeighbor ^44^ analyses established the robustness of these Group-level identities, guiding refinement of cluster boundaries and ensuring maximal correspondence of assignments across primates (Figure S1.4, Table S1.4). To maintain continuity with established BG terminology, Group-level nomenclature was grounded in conserved marker genes (Table S1.5) and shared molecular signatures while incorporating naming conventions from primate and rodent literature. These Group identities anchor analyses to the BICAN consensus taxonomy and unify BG nomenclature across species.

Spatial transcriptomics using MERSCOPE (human, macaque) and Xenium (marmoset) mapped Group identities onto anatomical regions^32^, while Patch-seq (macaque) linked them to morpho-electric phenotypes (Ping et al., co-submitted)^33^. Combined with iterative integration into transcriptomic clustering, these modalities resolved populations with subtle transcriptomic differences and sharpened Group boundaries. Spatial profiling across all primate species precisely localized MSN Groups, particularly matrix and striosome populations of the dorsal STR, and reproduced the expected compartmental patterns at the population level (Figure 1E). Patch-seq recordings from macaque STR revealed characteristic firing properties and morphological reconstructions of defining dendritic architectures corresponding to major transcriptomic Group terms (Figure 1E). Integrative multiomic analysis was critical for resolving ambiguities that could not be addressed by single-nucleus transcriptomics alone, resulting in a consensus taxonomy of the BG that provides a unifying, high-resolution cell type reference.

### Expression variation across primates and regions

Most BG Groups exhibited strong cross-primate correlations in RNA expression (r > 0.8), with marmoset showing greater divergence than human and macaque (Figure S2.1), consistent with known evolutionary relationships. Non-neuronal populations, especially microglia, were more divergent than neuronal cell types, in agreement with findings from primate neocortex ^45,46^. Despite this overall conservation, each Group exhibited thousands of 1:1 orthologous genes with at least two-fold expression differences in at least one pairwise cross-species comparison, and up to 30 1:1 orthologs with at least two-fold sex-biased expression (male versus female) within a species (Figure S2.1). Expressolog ^47^ analyses further identified genes with deeply conserved expression magnitude and pattern, such as *ADORA2A*, as well as species-biased shifts exemplified by *MCHR2* (Figure 2A). We also observed lineage-dependent rewiring of compartment-defining markers, including the primate-conserved *KCNIP1* and canonical rodent-biased *OPRM1* ^48^, the latter showing a lack of striosome specificity in primates (Figure 2A), underscoring both conserved and lineage-specific regulatory pressures shaping BG cell type identity.

**Figure 2.**
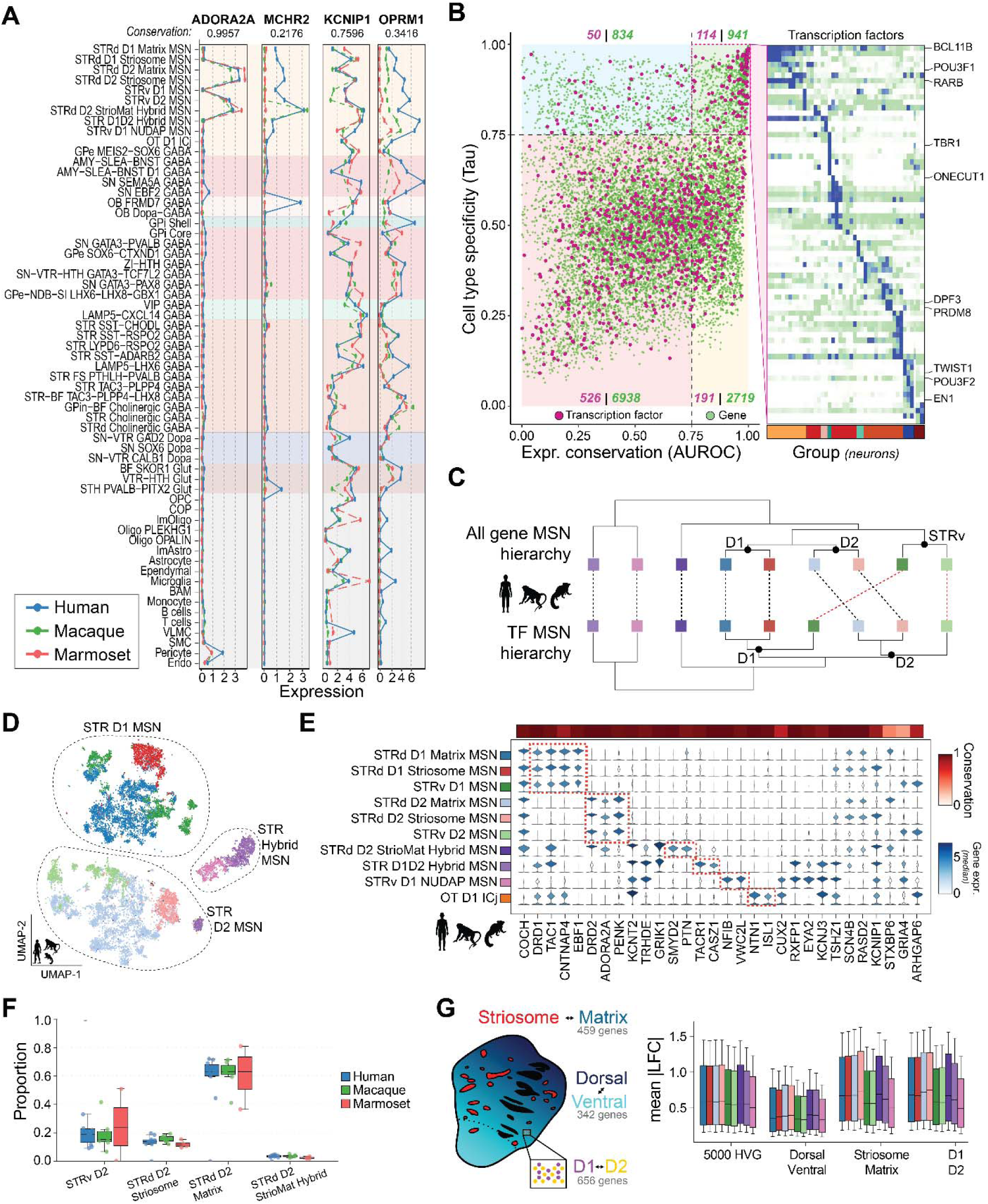
Comparative primate transcriptomics. (**A**) Group expression levels in each species for four genes with conserved or divergent conservation (mean expressolog score listed at top) and broad or cell type-enriched expression. Note that the expression range varies for each gene. Color shading indicates major divisions of cell Groups. (**B**) Left: Gene expression conservation (x-axis) versus cell type specificity (y-axis). The plot is divided into four quadrants and labeled with the total number of genes and TFs in each quadrant. Color bar corresponds to Group shading shown in (A). Right: Heatmap of conserved TF markers showing the column-scaled mean expression across species. (**C**) Dendrograms indicating the transcriptional relatedness of MSN Groups based on all genes or only TFs and colored as in (E). Major branches are labeled based on Group identities of leaves. (**D**) UMAP plots showing cross-species integrated MSNs, colored by Group as in (E). (**E**) Stacked violin plot showing expression of MSN Group markers across three primates. Conservation (AUROC) values were computed using data from all three species. (**F**) Boxplot showing the relative proportions of D2 Groups, highlighting the relative rarity of STRd D2 StrioMat MSN. (**G**) Left: Schematic showing canonical axes of variation of MSNs in the STR and the number of DEGs associated with each axis (P-adjusted < 0.01, LFC > 1.5, expressed in 20% of cells in at least one species Group). Right: Boxplots showing the mean absolute value of log-fold change for each species relative to the 4 species mean for 5000 highest variance genes across MSN Groups, and genes from left. Whiskers represent 15th and 85th percentiles and colored by Group as in (E).

Based on expression conservation and cell type-specificity, we grouped genes into four evolutionary regimes: 1) 941 conserved Group markers (expressolog AUROC > 0.75, tau > 0.75); 2) 2719 conserved broadly expressed genes (AUROC > 0.75, tau < 0.75); 3) 834 divergent Group markers (AUROC < 0.75, tau > 0.75); and 4) 6938 divergent broadly expressed genes (AUROC < 0.75, tau < 0.75) (Figure 2B). TFs were significantly enriched among Group markers (adjusted Fisher’s exact P = 1.8×10^-6^; Figures 2B and S2.2), reinforcing their central role in establishing BG cell type identity. For example, *TBR1* and *ONECUT1* separated GPi core and shell neurons, each defining transcriptomic and spatially coherent populations^32^. Enrichment of *DPF3*, which shapes Group-specific enhancer landscapes, together with *PRDM8*, a determinant of GABAergic Group identity, suggests that TAC-expressing interneurons are defined by a distinct regulatory architecture not shared with other BG interneurons. Dopaminergic neurons were partitioned into SN SOX6, SN-VTR GAD2, and SN-VTR CALB1 Groups, characterized by divergent TF signatures including *TWIST1*, *POU3F2*, and *EN1*. Canonical regulators such as *BCL11B* marked the LGE-derived MSNs while *RARB* and *POU3F1* distinguished ventral and eccentric MSNs (Figure 2B).

MSNs are the predominant neuronal type in the BG, accounting for about 75 percent of BG neurons in the profiled primate species. When we hierarchically clustered Groups using expression of all 1:1 orthologous genes, ventral (canonical D1 and D2) Groups segregated clearly from dorsal Groups. When clustering was restricted to TF expression, the dominant axis instead separated all D1 from D2 Groups. Across both gene sets, eccentric Groups consistently branched as an outgroup to D1 and D2 Groups, consistent with their distinct developmental lineage (Figure 2C). A higher-resolution Cytosplore 2D projection of cross-species MSNs using highly variable genes (Figure S2.3) showed clear separation of classical (STR D1 MSN and STR D2 MSN) and eccentric (STR Hybrid MSN) subclasses, and D1 and D2 MSNs were separated into matrix, striosome and ventral Groups (Figure 2D, Supp Figure S2.3). Canonical MSN markers were conserved across species, with *DRD1* and *TAC1* enriched in D1 MSNs, *DRD2* and *PENK* enriched in D2 MSNs, and *RXFP1* marking eccentric MSNs ^23,27,43^ (Figure 2E).

We identified a novel STRd D2 StrioMat Hybrid MSN Group within the STR D2 MSN subclass, characterized by a distinctive admixture molecular profile and ambiguous classification as either classical or eccentric (Figure 2C). This hybrid D2 population is rare, comprising only 3.4% of classical STR D2 MSNs (Figure 2F) and 1% of all MSNs (Figure S2.4; Table S2.3) across species (Figure 2F), likely contributing to its absence from prior single-cell studies despite being transcriptomically distinct from previously reported MSN cell types. Canonical primate markers of both striosome and matrix MSNs, including KCNIP1 and STXBP6, were co-expressed, alongside shared expression of COCH with classical D1 and D2 MSNs and selective sharing of KCNT2 with eccentric MSNs, but notably lacking expression of the eccentric marker RXFP1 (Figure 2E). Expression of SMYD2 was selectively enriched in this population (Figure 2E), emerging as a strong primate-specific candidate marker and consistent with its identification as the top differentially methylated marker of STRd D2 StrioMat Hybrid MSNs in a companion epigenomic study of the human BG (Ding et al., co-submitted)^49^.

To evaluate the evolutionary conservation of MSN transcriptomic architecture, we focused on three approximately orthogonal axes: D1 and D2 receptor expression, striosome and matrix compartmentalization, and dorsal and ventral spatial gradients (Figure 2G). First, we defined gene sets associated with D1-D2 and striosome-matrix axes in each species by running Wilcoxon differential expression tests between the contrasts. We identified 656 differentially expressed genes (DEGs) between D1 and D2, 459 DEGs between striosome and matrix, as well as 342 genes that had similar expression patterns to CNR1 (absolute Spearman correlation > 0.3 in any species), a gene which has robust dorsal-ventral gradient expression in spatial transcriptomics profiling of the primate STR ^32^. Comparing these genes sets to the 5,000 most highly variable genes across MSN Groups, we quantified cross-species divergence as the mean absolute deviance of log-expression across aligned MSN types between all four species. D1-D2 and striosome-matrix axes were significantly more diverged than highly variable genes, whereas dorsal-ventral gradient genes were significantly less diverged. This suggests that dorsal-ventral patterning of MSNs is under stronger evolutionary constraint than other aspects of their molecular identity within a striatal subregion (Figure 2G).

### Primate and mouse cell type homologies

To place the HMBA BG taxonomy in the context of BG profiling within recent whole-brain atlases, we integrated it with mouse ^17,18^ and human ^19^ snRNA-seq data derived from BG regions. The cross-atlas alignments showed clear cell type homologies and greater sampling of all Groups (Figures 3A, S3.1 and S3.2; Tables S3.1, S3.2, and S3.3) that increased detection of cellular diversity in this study. For example, the HMBA taxonomy defines three transcriptionally and spatially distinct cholinergic Groups ^32^, whereas the mouse and human atlases grouped these together (Tables S3.1 and S3.2). Several Groups lacked homologies with mouse whole brain due to low sampling, species differences in anatomical distributions, or inclusion of cell types in the HMBA taxonomy from regions adjacent to BG. GPi Core and Shell groups were missing from the mouse snRNA-seq data ^18^ and were subsequently identified using whole-brain single-cell RNA-seq data (Tables S3.1) ^17^. OB-Dopa-GABA and OB FRMD7 GABA Groups were not initially detected in mouse BG, and further analysis showed that this was a species difference as discussed below. Finally, the SN EBF2 GABA and SN SEMA5A GABA Groups lacked homologous types in the mouse and human atlases, and nuclei may have been dissected from midbrain nuclei adjacent to substantia nigra.

**Figure 3.**
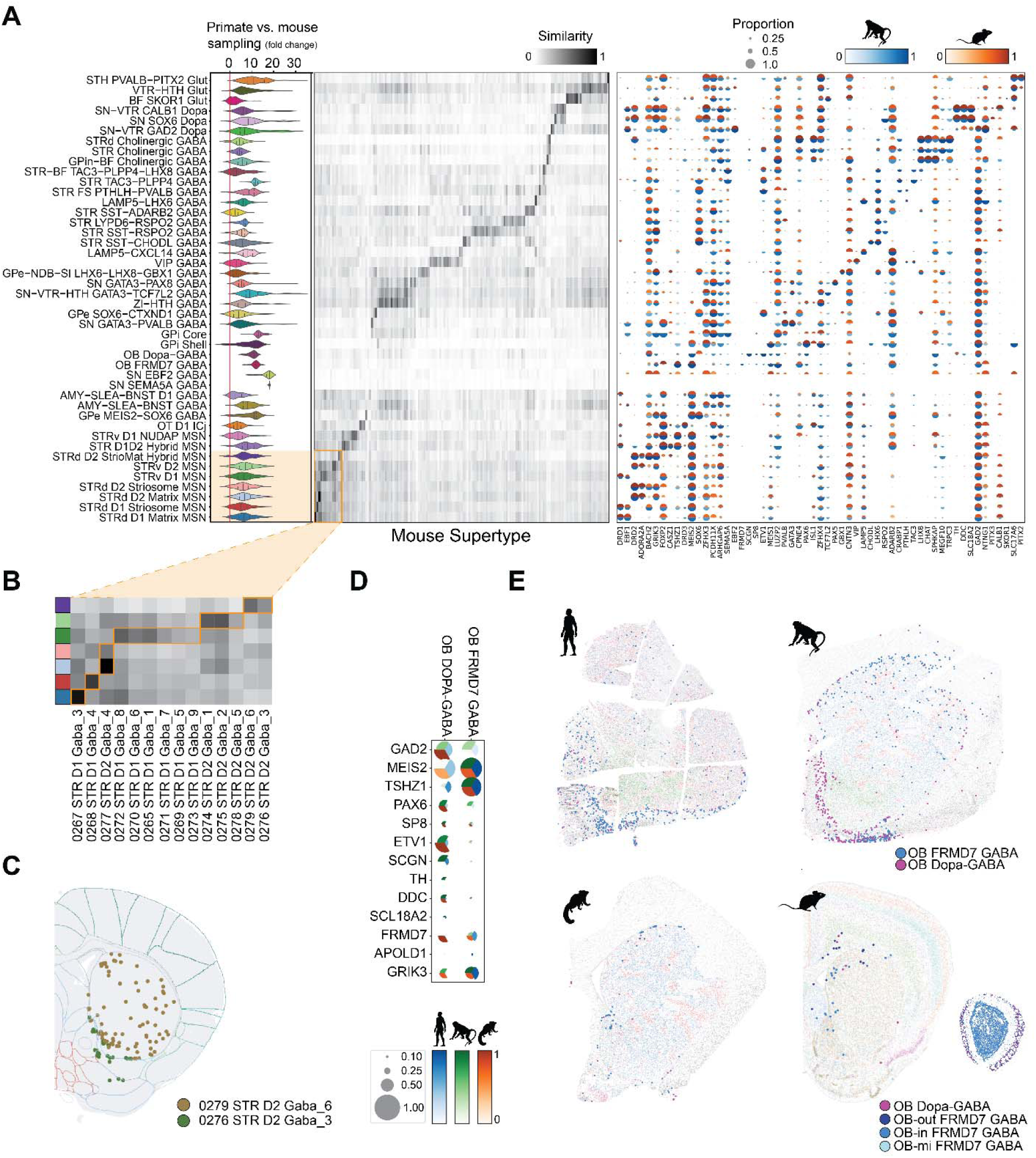
Primate and mouse cell type homologies. (**A**) Comparison of primate BG cell sampling and groups in this study versus mouse BG cell sampling and supertypes. Fold change indicates the increase in sampling depth based on a Milo analysis of the integrated data sets. Cell type similarities and conserved markers between macaque and mouse. (**B**) Mouse supertypes that correspond to a subset of MSN groups in this study. (**C**) STRd D2 StrioMat Hybrid MSN homologous supertypes in mouse caudoputamen and nucleus accumbens. (**D**) Pie dot plot showing the expression level (color) and proportion of nuclei with expression > 0 (size) of primate conserved markers for two Groups of OB-related cell groups. (**E**) Peri-striatal distribution of OB Groups is conserved across primates and mouse. The homologous mouse supertypes are enriched in OB (inset at bottom right) and were not included in the cell type homology analysis shown in (A).

The HMBA literature-grounded nomenclature resolves previously ambiguous clusters in whole-brain atlases. We observed greater MSN diversity in primates, with 130, 85, and 132 MSN clusters identified in human, macaque, and marmoset, respectively, compared with the 73 MSN clusters reported in mouse (Yao et al. 2023). Matrix, striosome, and ventral MSN Groups are represented as unlabeled supertypes of STR D1 MSN and STR D2 MSN subclasses in the mouse atlas (Figure 3B) or as unlabeled clusters of MSNs in the human atlas (Figure S3.2). The newly characterized STRd D2 StrioMat Hybrid MSN Group maps to human MSN_221 and two mouse supertypes that are broadly distributed in STR (0279 STR D2 GABA_6) or ventrally enriched (0276 STR D2 GABA_3; Figure 3C). Primate STRd D2 StrioMat Hybrid MSNs show a conserved dorsally enriched spatial pattern, with a subset of ventrally localized clusters marked by *HTR7* (Figure S3.3). The 52 conserved markers that are differentially expressed in STRd D2 StrioMat Hybrid MSNs compared to canonical STR D2 MSNs (Figure S3.4A and Table S3.3) include genes expressed by STR D2 and eccentric MSNs (Figure S3.4B), suggesting this newly-characterized hybrid may share features of both these MSN Groups. Conserved markers also include synaptic adhesion molecules ^50^ (e.g., *FAT1*, *SLITRK1/4*, *NTNG1*, and *PCDH10/15*) that may contribute to the distinct connectivity of this MSN population.

Surprisingly, two related Groups share expression patterns with outer layer olfactory bulb (OB) neurons and are sparsely distributed at the white-matter margins of the STR and near the rostral migratory stream in human, macaque, and marmoset (Figure 3E). OB Dopa-GABA shares *PAX6*, *TH*, *SCGN*, and *SP8* markers with previously identified STR laureatum neurons ^51^, and *DDC* and *SLC18A2* are newly described markers (Figure 3D). In contrast, OB FRMD7 GABA represents a previously unrecognized striatal Group. Homologous Groups were extremely rare in mouse peri-striatal regions (Figure 3E).

### Conserved and human-specialized cCREs

To characterize gene regulatory evolution in primate BG, we generated cell type-resolved cCRE maps from snATAC-seq profiles in human, macaque, and marmoset, identifying 1.36M, 1.37M, and 1.22M accessible regions, respectively. We quantified sequence conservation for each species-specific cCRE using an updated 447 mammal Cactus genomic alignment and reconstructed ancestral genomes from the Zoonomia Consortium (see Methods)^52^, enabling comparison across 239 primate and 208 non-primate species (Figure 4A). For comparative analyses, we classified cCREs by sequence alignment status and chromatin accessibility patterns (Table S4.1, Figure S4.1). Species-specific cCREs were defined as regions lacking ≥50 percent sequence alignment in any other primate, and sequence-conserved cCREs as regions that were aligned across human, macaque, and marmoset. Comparative analysis of chromatin accessibility across primates was performed using an accessolog metric that quantifies similarity in Group accessibility patterns across sequence-conserved regions (see Methods) similar to expressologs ^47^. We designated epi-conserved cCREs with high accessolog scores and species-biased cCREs as those with elevated cell type specificity (tau) in only one species (Figure 4A).

**Figure 4.**
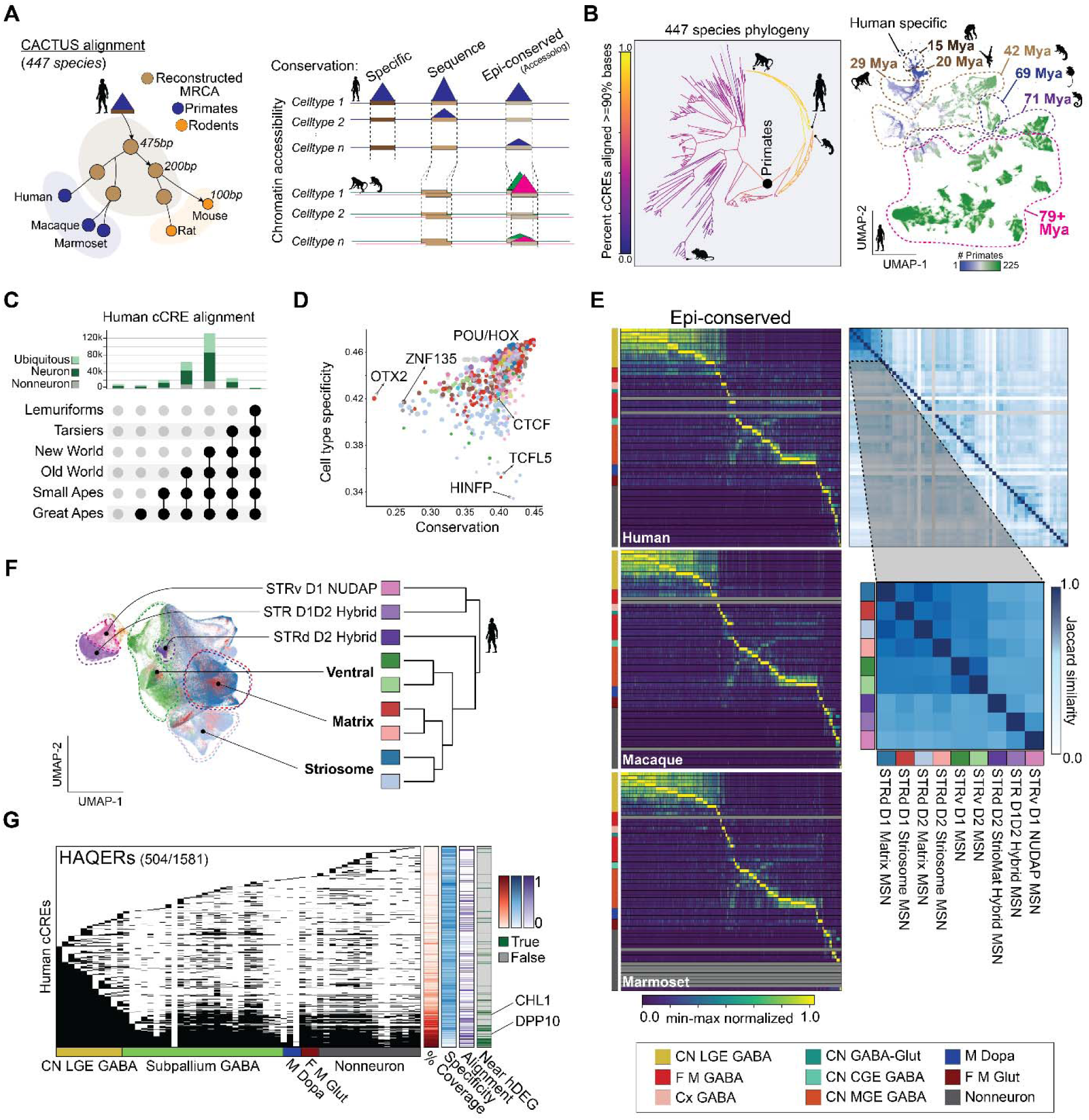
Human cCRE evolution across mammals. (**A**) Schematic of conservation patterns for cCRE sequence alignment. (**B**) Human cCRE conservation across mammals and primates. Percent of human cCREs aligning to each node in the 447 species phylogeny. UMAP projection of human cCREs (1.3M) by the count of positions aligning to the 447 mammalian genomes; color represent the count of primates to which each cCREs aligns. (**C**) Number of human open chromatin regions conserved across specific primate clades with bars colored by Neuron and Nonneuron. (**D**) TF motifs enrichment in human cCREs across cell types and primate species. Each point is a unique TF-motif colored by TF Class annotation. (**E**) Heatmaps of conserved Group-specific cCRE accessibility where gray bars indicate cell types with either low sampling or missing in a particular species. Jaccard similarity of cCRE accessibility across Groups in human. (**F**) Left: UMAP of snATAC-seq colored by MSN Groups. Right: Dendrogram showing similarity of MSN Groups measured by Jaccard distance on binarized peak calls for human. (**G**) Human cCRE intersection with rapidly evolving genomic regions annotated with cCRE cell type, alignment and hDEG proximity metrics.

Genome-wide analysis of human cCREs revealed extensive regulatory turnover within primates over 76 million years alongside a deeply conserved regulatory core maintained across mammals (Figure 4B). UMAP embedding of cCRE base-pair alignment fractions across 446 non-human genomes grouped cCREs by shared evolutionary history. The resulting groups included cCREs that arose at different ancestral nodes along the primate lineage but at similar times in neuronal and non-neuronal Groups (Figure 4C, Figure S4.2), which is striking given that non-neuronal expression has diverged more rapidly than neuronal expression. Human-specific cCREs (0.0009%) were enriched for transposable regions previously associated with rapid evolutionary change ^53^, whereas sequence and epi-conserved cCREs (46%) were enriched near promoters (Figure S4.1).

To define the regulatory features associated with conservation levels, we examined TF motif enrichment across conserved and divergent human cCREs (Figure 4D; Table S4.2). Conserved cell type-selective cCREs were enriched for POU, MEIS, and HOX family motifs, consistent with the role of these TFs in defining neuronal identity during BG development ^54,55^. More ubiquitous cCREs were enriched near promoters and for HINFP and TCFL5 motifs, reflecting the evolutionary stability and functional importance of regions linked to housekeeping and early developmental genes. By contrast, cCREs that emerged more recently in the human lineage were enriched for OTX2 and ZNF family motifs. Notably, OTX2 is associated with diencephalic origin while regulating dopaminergic neuron function ^56^, neural plasticity ^57^, and has been associated with depression risk ^58^.

Combining accessolog scores with specificity (tau) revealed epi-conserved marker cCREs for all Groups, with expanded sets of markers of some neuronal Groups, including cholinergic neurons. In contrast, non-neuronal Groups had comparatively few conserved elements, reflecting more rapid regulatory turnover. Among MSNs, conserved and species-enriched peaks robustly separated matrix, striosome, ventral, and eccentric Groups (Figures 4E, 4F and S4.3). Surprisingly, D1 and D2 Groups share most cCREs that are bound by differentially expressed TFs (Figures 2C and 2E) to regulate distinct D1 and D2 transcriptional profiles. The STRd D2 StrioMat Hybrid MSN Group exemplified this complexity, displaying an intermediate accessibility profile bridging STR D1D2 Hybrid and striosome identities, suggestive of dual regulatory program integration within a single lineage.

To assess how recent human evolution has reshaped BG regulatory programs, we intersected cCREs with Human Accelerated Regions (HARs), Human Ancestor-Qualified Evolutionary Regions (HAQERs) that are divergent from chimpanzee, and human-specific deletions (hCONDELs) (Figure 4G). These rapidly evolving sequences overlapped broadly accessible and cell type-restricted cCREs, suggesting that evolution has acted on regulatory sequences governing core BG functions and cell type-specific identities (Figure S4.4). Although many HAQERs show limited conservation beyond apes, Cactus alignments revealed a subset (n=113, mean evo-distance=0.64) whose underlying cCREs are retained across distant mammalian lineages (Figure 4G; Table S4.3), suggesting that human-specific substitutions accumulated atop deeply conserved regulatory scaffolds. The activity-by-contact model ^59^ revealed HAQER-associated cCREs located near human DEGs (hDEGs), including *DPP10*, a gene with divergent expression in human compared to macaque and marmoset (Figure 1A). These findings show that recent human evolution has modified both conserved and lineage-specific cCREs in the BG, providing regulatory mechanisms for primate- and human-specific neuronal features.

### Cell type-specific enhancer codes across evolution

To define conserved DNA sequence grammars underlying Group-specific chromatin accessibility, we trained species-matched dilated convolutional neural network models ^60^ on human and macaque snATAC-seq datasets linked to the multiomic BG taxonomy (Figure 5A). The DeepHumanBG and DeepMacaqueBG models predict chromatin accessibility directly from cCRE DNA sequence across all annotated Groups. Models for both species achieved strong performance on held-out chromosomes (Spearman r = 0.71, human; r = 0.75, macaque) and accurately reproduced Group accessibility profiles (Figure S5.1). This indicated that the DNA sequence models learned to encode TF motif grammars that specify BG cellular identity.

**Figure 5.**
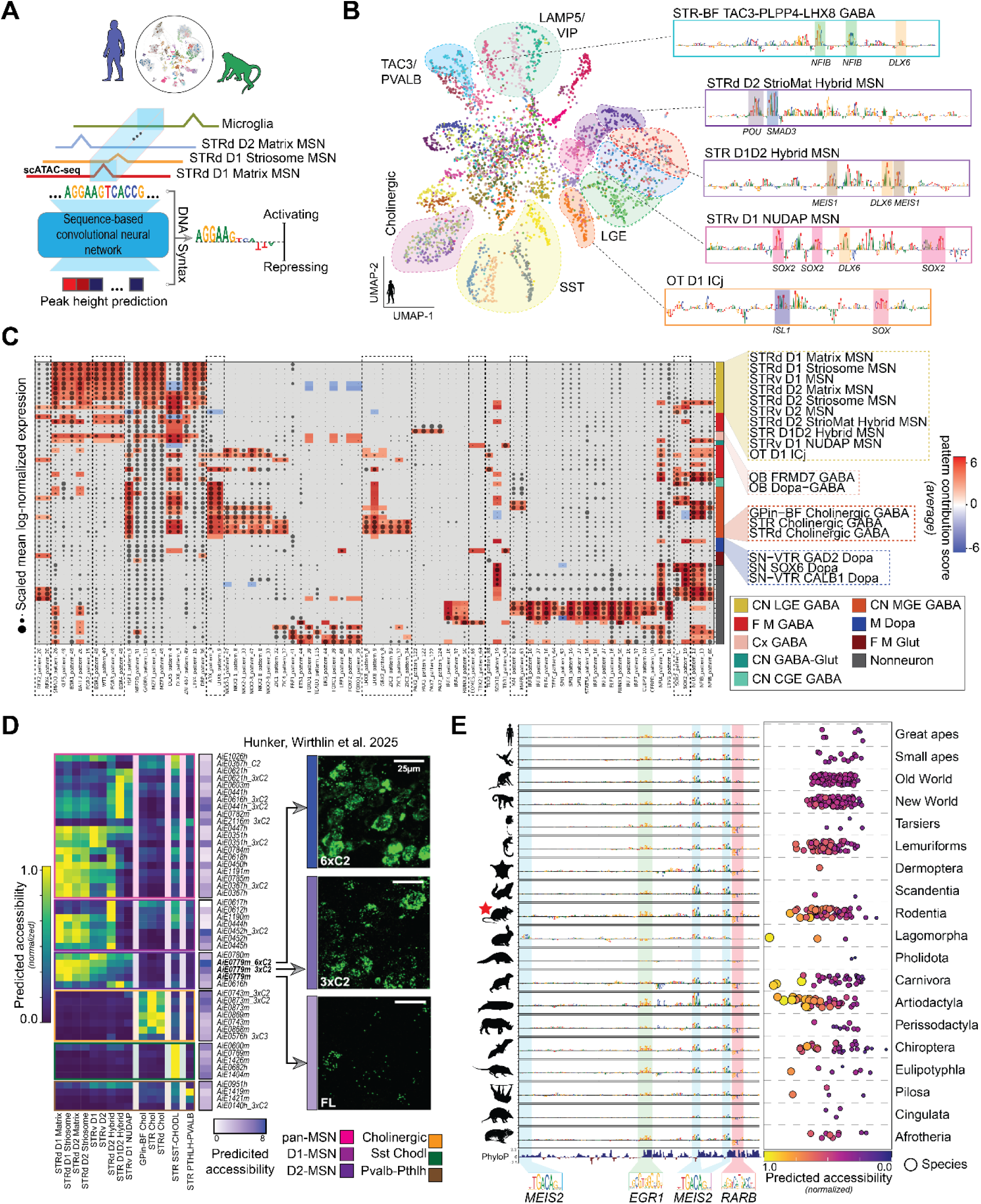
Conserved cell type-specific TF grammar. (**A**) Schematic of the training procedure for each species CRESted model and description of the per-base importance scores. (**B**) UMAP of DeepHumanBG latent embeddings for the top 100 Group-specific cCREs. Contribution score plots of highly specific peaks for select cell types are displayed on the right with TF-motifs highlighted. (**C**) Clustermap of scaled mean log-normalized TF expression over Groups, indicated by dot size. Sequence patterns from DeepHumanBG matched to known TF motifs (columns) with mean importance per Group indicated by color. (**D**) DeepHumanBG Group normalized predicted accessibility heatmap for viral tools from Hunker, Wirthlin et al. 2025^61^. Heatmap annotation for the magnitude of predicted accessibility aggregated across Groups targeted by each enhancer alongside in-vivo quantification of enhancer activity. (**E**) Comparative analysis of AiE0779m; sequence importances for species with the highest predicted activity of AiE0779m per-clade across the Cactus alignment. Scatterplot of each species (dot) organized by clade and colored by normalized predicted strength.

To examine the sequence features underlying these predictions, we passed the top 100 neuronal Group-specific human cCREs through the DeepHumanBG model and extracted their internal sequence embeddings. After dimensionality reduction, the embeddings formed discrete clusters corresponding to neuronal Groups, as well as a central set of multi-Group cCREs (Figure 5B). Embedding-based classification saturated early during training, reflecting rapid convergence on stable and biologically meaningful sequence features that delineate nearly all Groups in both human and macaque (Figure S5.1). Extending this analysis to the top 100 Group-specific cCREs from both species revealed a highly congruent embedding structure, indicating that the human and macaque models convergently learned conserved sequence grammar. Likewise, predicted accessibility for Group-specific cCREs was strongly correlated across species, demonstrating that the relationship between DNA sequence and chromatin accessibility is evolutionarily preserved (Figure S5.2). Consistent with the snATAC-seq data, both models robustly resolved matrix, striosome, ventral, and eccentric MSN Groups, and neither model distinguished canonical D1 and D2 MSNs. Thus, MSN Group identity is encoded by distinct TF motif grammars, while D1 and D2 counterparts of matrix, striosome and ventral MSN Groups share an overlapping grammar.

Nucleotide-level contribution scores for representative cCREs revealed TF motifs predictive of Group-specific accessibility, providing direct sequence-level evidence for the enhancer codes underlying each cell type (Figure 5B). Within the eccentric MSN lineage, POU and SMAD3 motifs were uniquely enriched in the STRd D2 StrioMat Hybrid, whereas DLX6 motifs were shared across the STR D1D2 Hybrid and STRv D1 NUDAP, mirroring their transcriptional and chromatin-level similarity. To systematically map DNA grammar across Groups, we applied a global motif-importance analysis (Figure 5C), integrating model-derived motif enrichments with snRNA-seq expression to assign candidate TFs to each Group (Methods). Cholinergic neurons exhibited a coherent motif signature composed of LHX8, GBX2, and ZIC3 elements, with subtle ZIC3 variants distinguishing neighboring Groups; notably, GBX2 was selectively enriched in striatal cholinergic neurons but absent from the interstitial globus pallidus cholinergic population (GPin-BF). Across GABAergic neurons, DLX5/6 motifs were broadly important yet displayed Group-specific polarity strongly enriched in eccentric MSNs, attenuated in dorsal matrix and striosome MSNs, and negatively weighted in ventral MSNs indicating that TF families such as DLX, LHX, ZIC, and POU are reused across MSNs but deployed with quantitatively distinct motif grammars to encode Group identity. In addition to motifs with assignable TF identities, numerous additional sequence patterns exhibited strong cell type specificity, suggesting the existence of enhancer grammars not attributable to known TFs (Figure S5.4). Performing the same analysis in macaque yielded highly concordant TF-Group associations, demonstrating that core enhancer grammars are conserved across primates (Figure S5.3-5).

To identify candidate TFs driving enhancer functional activity, we applied DeepHumanBG to a published collection of enhancer-driven AAVs ^61^ that target broad populations of MSNs and striatal interneurons (Figure 5D). Model-predicted activities closely matched published targets and resolved higher-resolution Group targets. For example, AiE0441h and its optimized variants described as D1-selective were predicted to preferentially target D1D2 Hybrid MSNs, separating them from classical D1 matrix and striosome populations. The model identified AiE1419m as the most selective enhancer for fast-spiking PVALB interneurons and AiE1404m as the strongest enhancer for SST Chodl interneurons. Strikingly, we correctly predicted increased activity of optimized variants of AiE0779m (Figure 5D), consistent with sequence-based models outperforming epigenomic measurements alone in predicting strength of enhancer activity in the neocortex ^62^.

To assess how enhancer codes operate across evolution, we aligned each of the published enhancer sequences from the reference species (human, hg38 or mouse, mm10) to the remaining mammalian genomes in the 447-way Cactus alignment. For each enhancer ortholog, we used DeepHumanBG to predict activity in the Groups that corresponded to the target population of the enhancer. Many enhancers showed conserved predicted activity across mammals, whereas others retained alignable sequence only within restricted clades, suggesting lineage-dependent divergence in enhancer logic and function (Figure S5.6). For example, the D1-MSN enhancer AiE0779m sequence was predicted to be active across diverse mammals with MEIS2, EGR1, and RARB motifs as the principal drivers of specificity (Figure 5E). Yet several order-specific deviations emerged: Pholidota (pangolins) and Cingulata (armadillos) orthologs showed disrupted MEIS2/EGR1/RARB motifs with corresponding predicted loss of D1-MSN activity, and Lagomorpha (specifically rabbits and hares) orthologs acquired a novel MEIS2-like motif absent from all other lineages. These findings demonstrate that the TF-motif grammars conferring cell type selectivity are deeply conserved, and also evolve through clade-specific rewiring.

### Disease associations with BG cell types

To identify the BG cell populations from human that most strongly linked to psychiatric disease risk, we combined MAGMA-based GWAS enrichment analysis with our high-resolution consensus taxonomy (Figure 6A). We evaluated four psychiatric GWAS ^63–66^, an alcohol-use phenotype^67^, and two benchmarks expected to enrich in non-neuronal immune populations: Alzheimer’s disease (AD) GWAS ^68^ and ancient DNA time-series estimates of recent directional selection in West Eurasians over the past ∼14,000 years ^69^. This analysis first revealed the overall structure of phenotype and cell type relationships across the full BG taxonomy (Figure S6.1).

**Figure 6.**
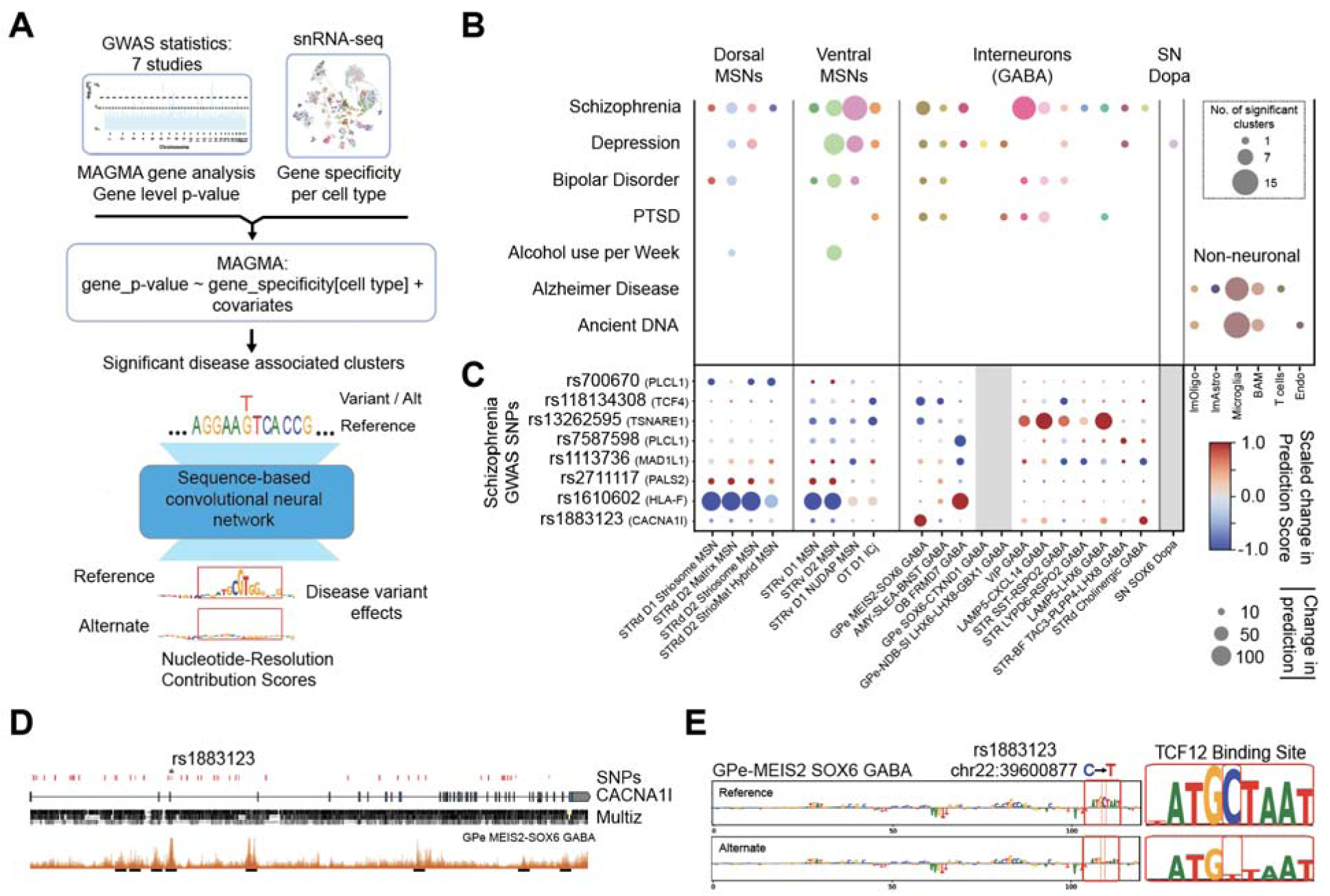
Cell type-specific associations with human brain diseases. (**A**) Schematic of cell type enrichment analysis using MAGMA with SNPs from 7 large GWAS and disease variant prioritization using the CREsted sequence model. (**B**) Significant (P < 0.0001) phenotype associations with BG cell types using MAGMA. Dot size indicates the number of significant clusters per group. (**C**) Dot plot illustrating the predicted effects of significant SCZ-associated SNPs on cell type-specific chromatin accessibility. The size of the dots corresponds to the magnitude of the effect, while the color indicates the direction (signed effect) of the effect. (**D**) SNPs significantly associated with SCZ overlapping the *CACNA1I* gene. Schematic representation of *CACNA1I* exons and introns, Multiz alignment of human, macaque, and marmoset, and human chromatin accessibility from GPe Meis2 Sox6 GABA neurons with cCREs indicated by lines. rs1883123 (also shown in (C)) is located in an intronic cCRE. (**E**) The SCZ-associated allele of rs1883123 is predicted to reduce the activity of a TCF12 motif binding site within this cCRE, specifically in the GPe Meis2 Sox6 GABA Group.

Across disorders, MSNs showed the strongest enrichment (Figure 6B). Ventral MSNs accounted for most significant clusters, including 14 STRv D1 NUDAP MSN clusters linked to SCZ. Our atlas also detected SCZ associations in MSN populations not previously implicated, including the STRd D2 Striomat Hybrid MSN Group (Figure S6.2), and established clear correspondence between our BG-associated SCZ clusters and those identified in whole-brain analyses^19^. Many of these associations overlapped with signals previously reported in genome-wide studies of SCZ risk^35–37^, although our higher-resolution atlas also revealed MSN-specific enrichments that were previously unresolved. Analyses incorporating larger SCZ cohorts expanded the range of MSN and interneuron associations (Figure S6.3), underscoring the need to profile more non-European individuals to detect additional disease-associated variants relevant to those populations.

Cross-disorder comparisons demonstrated both shared and distinct patterns (Figure S6.4). SCZ, bipolar disorder (BD), and major depressive disorder (MDD) exhibited highly correlated association profiles, including VIP (13 clusters) and MGE-derived interneurons. In contrast, post-traumatic stress disorder (PTSD) and MDD selectively engaged the GPe-NDB-SI LHX6-LHX8-GBX1 interneurons, and MDD uniquely mapped to two dopaminergic clusters (Figure 6B). As expected from previous work, AD GWAS signals localized exclusively to non-neuronal cell types, with significant enrichment in 13 microglial clusters and one T-cell cluster (Figure 6B). This pattern aligns with the immune-dominant architecture of AD ^70^ and recent findings implicating adaptive immune contributions to AD risk ^71^ linking caudate non-neuronal cell populations with disease progression in AD (Kana et al., co-submitted)^72^.

To connect enriched cell types to potential causal variants, we intersected genome-wide significant SNPs with cell type-specific open chromatin and applied the DeepHumanBG model to predict allele-specific regulatory effects (Figure 6C). Several variants showed strong, cell type-restricted differences in predicted accessibility. For example, rs1883123 in CACNA1I, a SCZ-associated locus^73^, decreased predicted accessibility selectively in GPe-MEIS2-SOX6 GABA neurons (Figures 6C and 6D), consistent with a predicted disruption of a TCF12 motif (Figure 6E). Together, the MAGMA and DeepHumanBG analyses map disease-associated loci to specific BG cell types and nominate candidate regulatory variants likely to exert cell type-specific effects.

## DISCUSSION

This study generates a cross-species, single-nucleus multiomic reference for the primate BG that facilitates mechanistic and genetic dissection of BG circuits. By jointly constructing a consensus taxonomy, a cis-regulatory atlas with sequence-based models, and a cell type-resolved map of genetic risk, we identify three overarching principles for primate BG organization: (1) canonical BG axes of organization are not evolutionarily equivalent, with dorsoventral spatial patterning more conserved than direct-indirect pathway or striosome-matrix compartment molecular identity; (2) BG cell types are specified by modular TF grammars that predict enhancer activity and can be used to generate hypotheses about noncoding variant effects across species; and (3) common variant risk for psychiatric disorders is non-uniform and is preferentially enriched in specific MSN and interneuron Groups, including a novel hybrid MSN population.

Classical models of BG function emphasize direct and indirect pathway MSNs, striosome-matrix compartments, and parallel cortico-BG-thalamo-cortical loops as substrates for action selection and reinforcement learning ^1,11,74^. Our consensus taxonomy recovers this architecture while placing projection neurons, interneurons, and non-neuronal populations into a unified hierarchy of Neighborhoods, Classes, Subclasses, and 61 Groups shared across three primates and aligned to mouse and human whole-brain atlases. Because taxonomic boundaries are informed by paired RNA-chromatin profiles from the same nuclei and anchored by spatial mapping, Groups are both molecularly coherent and anatomically interpretable, including BG-specialized populations that are under-sampled in whole-brain references.

Prior single-cell studies in STR described multiple MSN subclasses and continuous gradients but could not systematically assess how these axes differ in evolutionary stability. By quantifying divergence of gene sets associated with each axis across three primate species, we find that dorsal-ventral expression is more conserved than D1-D2 or striosome-matrix programs, indicating that spatial patterning is under tighter evolutionary constraint and that pathway and compartment identities are more flexible features layered on this spatial scaffold. More generally, representing each axis as a gene program and quantifying its evolutionary divergence can be applied to additional BG axes and to developmental or disease-related state transitions.

The STRd D2 StrioMat Hybrid Group exemplifies this principle, where mapping across primates and mouse unifies previously unlabeled MSN clusters in whole-brain atlases. This population exhibits a reproducible dorsal-ventral subdivision marked by *HTR7* and a characteristic set of synaptic adhesion genes shared with eccentric MSNs. In contrast to DRD2- and HTR7-positive populations previously reported to coexpress TH in rodent STR ^23,26^, primate STRd D2 StrioMat Hybrid neurons lack dopaminergic markers, clarifying their identity as a distinct MSN lineage rather than a dopaminergic hybrid. Combined transcriptomic, spatial ^32^, and Patch-seq data (Liu et al., co-submitted)^33^ indicate that these neurons occupy stereotyped positions in dorsal and ventral STR and have distinct morpho-electric properties, suggesting that they implement circuit motifs that are not easily captured by a simple direct-indirect or striosome-matrix dichotomy.

A notable primate feature revealed by the atlas is a peri-striatal shell of OB-like GABAergic neurons, resolved into two Groups (OB Dopa-GABA and OB FRMD7 GABA) that are scattered along striatal white-matter margins near the rostral migratory stream and are strongly depleted in mouse ^17^. Multiomic profiles show that these neurons express a PAX6/MEIS2/SP8 transcriptional program characteristic of LGE-derived OB periglomerular interneurons^75–77^. OB Dopa-GABA matches the STR laureatum population described in macaque, and here we show that like its OB sister cells, it expresses the full complement of dopaminergic genes^51^. The novel OB FRMD GABA Group is more dorsally and medially distributed, but additional data on its morphology, connectivity and electrophysiology will be required to determine its function. Work on the olfactory tubercle and ventral STR has emphasized these regions as hubs where olfactory inputs converge with limbic circuits to encode odor valence and reward ^78,79^. It remains possible that these OB-like Groups participate in non-olfactory functions; for example, OB Dopa-GABA may provide supplemental dopaminergic input to striatal MSNs.

Matched snATAC-seq across three primates identifies over one million cCREs per species, extending prior cortex- and STR-focused studies that lacked BG-wide regulatory coverage in primates ^80–83^. By quantifying sequence conservation and cell type-specific accessibility, we find that promoter-proximal elements are disproportionately conserved at both sequence and activity levels, in line with these atlases. Distal enhancers, in contrast, partition into conserved, primate-biased, and human-biased regimes, with the human-biased subset showing cell type-specific overlap with human-accelerated genomic regions. Chromatin accessibility distinguishes matrix, striosome, and ventral MSN Groups but not underlying D1 and D2 populations, whereas gene expression shows clear separation of these lineages.

Sequence-based CREsted models trained on these cCREs (DeepHumanBG and DeepMacaqueBG) recover modular TF grammars that specify Group-level enhancer activity. Inhibitory lineages rely on DLX-and MEIS-related motifs, while particular MSN and pallidal projection neuron Groups additionally use LHX- and POU-related motifs. Importantly, within a given MSN Group, D1- and D2-expressing neurons share highly similar enhancer grammars even when their transcriptomes diverge. Together with the accessibility analysis, these results suggest that D1 and D2 pathway identities arise from additional regulatory layers acting on broadly shared cis-regulatory elements, including differential TF binding, histone modifications (Xie et al., co-submitted)^84^, DNA methylation, higher-order chromatin architecture (Ding et al., co-submitted)^49^, or circuit-level mechanisms. Cross-species prediction further suggests that similar motif grammars can specify homologous BG types in different primates even when individual enhancers and surrounding sequence contexts diverge, consistent with a partial decoupling of regulatory logic from strict base-pair conservation.

Linking enhancer sequence to Group-level activity enables a forward model from regulatory grammar to experimental tools. Application of DeepHumanBG to enhancer-AAVs developed for striatal MSN and interneuron types ^61^ shows that predicted activity profiles broadly recapitulate observed *in vivo* activity across MSNs and interneurons, with reduced performance for distinguishing D1 versus D2 MSN activity, consistent with shared enhancer grammars between these pathways and limitations of the current training data. Using DeepHumanBG, we connect evolutionary changes in orthologous enhancer sequences to predicted changes in cell type-specific activity across 447 mammalian species. For the D1-MSN enhancer AiE0779m, the model predicts largely conserved D1-MSN activity across most mammals but also reveals clade-specific rewiring, with disrupted MEIS2/EGR1/RARB motifs and loss of predicted D1-MSN activity in pangolins and armadillos and a rabbit- and hare-specific gain of a novel MEIS2-like motif. Together with ongoing large-scale enhancer-AAV efforts ^85,86^ (also, see Wirthlin, Hunker et al., co-submitted)^87^, these cell type-specific regulatory grammars can guide the optimization and cross-species evaluation of viral vectors by prioritizing candidate enhancers and predicting their activity in defined BG cell types, as well as informing the rational design of synthetic enhancer-AAVs that maximize the strength and specificity of cell type targeting.

By integrating our BG taxonomy with MAGMA-based enrichment, we find that risk for SCZ, BD, and MDD is non-uniform and is preferentially enriched in specific BG MSN and interneuron modules, particularly ventral STRv D1 NUDAP and related ventral MSN Groups, as well as the STRd D2 StrioMat Hybrid Group. These findings extend whole-brain analyses by resolving risk from generic MSN categories to defined MSN Groups with known spatial organization and regulatory grammar and complement prior cortical enrichments that emphasize excitatory projection neurons and interneuron classes ^35–37^. Across disorders, VIP and MGE-derived interneurons show shared disease enrichment with the cortical classes highlighted in these studies, whereas PTSD and MDD exhibit more selective associations with GPe-NDB-SI LHX6-LHX8-GBX1 GABA interneurons and subsets of dopaminergic neurons, pointing to candidate BG microcircuits for distinct components of affective and stress-related risk. Overlaying genome-wide significant loci on BG cCREs and scoring alleles with DeepHumanBG further localizes these signals to specific regulatory elements and motifs in disease-relevant cell types. For example, within a CACNA1I intron, a SCZ variant falls in a GPe MEIS2 SOX6 GABA neuron cCRE where the risk allele is predicted to disrupt a TCF12 motif, providing a testable, model-based hypothesis for a route from noncoding variation to altered pallidal physiology.

This work provides a cross-species molecular atlas for the primate BG that links cell types, regulatory grammars, experimental tools, and genetic risk across human and model species. Future multimodal datasets can be mapped to this atlas, refining BG cell type definitions, and relating molecular programs to circuit function and behavior. Within this framework, cell type-resolved regulatory maps and sequence models support a systematic, hypothesis-generating “variant-to-cell-type-to-mechanism” workflow that can be tested using enhancer-based tools in defined BG circuits.

### Limitations of the study

We profiled adult postmortem BG from a limited number of donors per species, so the resource is not powered to assess sex, ancestry, disease, or acute state-dependent variation. Although dissections tiled dorsal-ventral and rostrocaudal axes and we used spatial transcriptomics and Patch-seq to anchor clusters, we likely under-sampled rare, localized populations and did not fully resolve all BG-adjacent populations. Our consensus taxonomy emphasizes 61 Group-level types aligned across the three primates using 1:1 orthologs and conserved gene programs, and we provide species-specific clusters and genes for investigation of specializations, some of which are explored in companion manuscripts.

At the regulatory level, sequence model performance and interpretability will benefit from additional epigenomic modalities and perturbation or reporter assays, particularly for closely related MSN Groups. Finally, our genetic enrichment analyses rely on common-variant GWAS, which are dominated by individuals of European ancestry and cis-focused gene-level tests. As a result, some cell types whose liability is mediated by other genetic architectures or contexts may be missed, and enrichments may include genes that modulate disease responses or compensatory remodeling rather than initiating pathology in those cells.

## RESOURCE AVAILABILITY

### Lead Contact

Further information and requests for resources and reagents should be directed to and will be fulfilled by the lead contact, Trygve E. Bakken.

### Materials Availability

This study did not generate new unique reagents.

### Data and Code Availability

- Sequencing data and enhancer validation data used in this project are all from previous studies and are publicly available. Accession numbers and links to the datasets are listed in the key resources table.
- All original code has been deposited at GitHub and is publicly available as of the date of publication. GitHub URLs and DOIs are listed in the key resources table.
- Any additional information required to reanalyze the data reported in this paper is available from the lead contact upon request.

## ACKNOWLEDGMENTS

This publication was supported by and coordinated through the BRAIN Initiative Cell Atlas Network (BICAN) (https://braininitiative.nih.gov/research/tools-and-technologies-brain-cells-and-circuits/brain-initiative-cell-atlas-network). This work was funded by the Allen Institute for Brain Science and by the National Institutes of Health R01CA296792 (ARa), U01MH128339 (JTT), U24MH130918 (PLR), U24MH130919 (PLR), U24MH130968 (JG, LF), U24NS133077 (JAM), and UM1MH130981 (ABC, ABh, ABu, ADG, AM, AMY, ATo, ATr, BK, BLe, BLo, BN, BRL, CDK, CM, CR, CS, DB, DH, DLJ, DM, DO, DR, DY, EF, EP, ESL, FMK, HZ, IK, JAM, JC, JGl, JGo, JGu, JSc, JT, JTT, KJ, KS, LN, MAT, MC, MH, MJ, MK, ML, MS, MT, MW, ND, NG, NGu, NJJ, NP, NVS, RC, RD, RDH, RF, RM, RS, SCS, SDaniel, SDi, SO, STB, TC, TEB, TLD, TM, TP, WF, WH, XL, YF, ZY); Netherlands Organization for Scientific Research NWO:024.004.012 (AV, TK); Research Foundation Flanders (FWO) PhD fellowship 1SH6J24N & V428025N (NK). The authors thank Evan Biederstedt and Mary Futey for updating the Cell Annotation Platform (https://celltype.info/) that facilitated refinement of the taxonomy, William Stauffer for contributing to cell type nomenclature, and the founder of the Allen Institute, Paul G. Allen, for his vision, encouragement and support.

## AUTHOR CONTRIBUTIONS

Tissue acquisition: ATr, CDK, EP, FY, JGl, MC, MJ, MK, MS, ND, RDH, RM, TC, TLD, WF, WH

Sample preparation and multiomic data generation: ADG, AM, AMY, ATo, ATr, BN, CR, CS, DB, DH, DLJ, DR, EP, ESL, JGl, JGu, JSc, KJ, KS, LC, MC, MJ, MK, ML, MT, ND, NGu, NP, NVS, RDH, RF, RM, STB, TC, TEB, TP, VENC, WH

Spatial transcriptomic data generation: BLo, DM, JC, MAT, MH, RDH, SCS

Data archive / Infrastructure: AA, ABC, ABh, ARa, AV, BLe, CM, DO, EF, GH, GS, JAM, JE, JGo, JSh, JT, KZK, LN, PLR, RC, RS, SDaniel, SO, TK, TM

Data analysis: ABC, ABu, AD, ARi, AV, BK, BLe, BLo, BRL, DY, ESL, FMK, IK, JC, JE, JGo, JT, KS, MAT, MD, MH, MS, NG, NJJ, NK, RC, RD, SCS, SDan, TEB, TK, XL, YF, YW, ZY

Data interpretation: ABu, AD, BK, BLo, BPL, BRL, DO, DY, ESL, FMK, JC, JG, JTT, LF, MAT, MD, MH, MS, MW, NG, NJJ, NK, RD, RDH, SAS, SCS, SDan, SDi, TEB, TLD, XL, YF, YW, ZY

Writing manuscript: AD, ESL, JG, LF, MS, NJJ, NK, TEB, YF

Funding acquisition: BK, BLe, CDK, DO, ESL, FMK, HZ, JTT, KS, RDH, TEB, WF, ZY

## DECLARATION OF GENERATIVE AI AND AI-ASSISTED TECHNOLOGIES

During the preparation of this work, the authors used OpenAI ChatGPT-5 to support certain coding tasks, assist with refining the main text to improve conciseness and readability, and identify potentially relevant citations. After using this tool, the authors reviewed and edited the content as needed and take full responsibility for the final content of the publication.

## STAR METHODS

### METHOD DETAILS

#### Human Cohort Selection and Preparation

Tissue collection was performed in accordance with the provisions of the United States Uniform Anatomical Gift Act of 2006 described in the California Health and Safety Code section 7150 (effective 1 January 2008) and other applicable state and federal laws and regulations. The Western Institutional Review Board reviewed the use of these tissues and determined that they did not constitute human subject research requiring institutional review board (IRB) review. Male and female individuals 18 to 70 years of age were considered for inclusion in the study. Donor screening involved review of clinical and psychological history, exposure data, toxicology, serological testing for infectious diseases, diagnostic neuroimaging evaluation, and comprehensive neuropathology evaluation to assess all categories of developmental, reactive, neoplastic, and neurodegenerative processes. Standard tissue quality metrics (RNA integrity evaluation, RIN; pH; tissue slab quality) were assessed for all donors and a RIN value of ≥7.0 was required for inclusion in the study. A structured consensus committee, including a bioethicist, reviewed donor information using well-defined inclusion and exclusion criteria and research prioritization methods. Donor tissues for the current study were obtained from six donors (3 male, 3 female ages 18-61 years) via the University of Washington BioRepository and Integrated Neuropathology (BRaIN) laboratory. A rapid autopsy was performed on each donor where the brain was removed, bisected along the midline, and each hemisphere was embedded in alginate. Thin (4mm) coronal brain slabs were generated, photographed, and frozen using a dry ice and isopentane bath. Sagittal cerebellum and axial brainstem slabs were separately generated for each donor brain. Tissue slabs were transferred to vacuum sealed bags and stored at -80°C. Tissues were transferred to the Allen Institute on dry ice and were stored at - 80°C until the time of dissection.

#### Macaque Cohort Selection and Preparation

Tissue samples were obtained from 3 adult male rhesus macaque monkeys (Macaca mulatta, 6-14 years of age). Prior to euthanasia, the animals underwent a series of cognitive behavioral paradigms in the Laboratory of Neural Systems at the Rockefeller University. Immediately after euthanasia, brains were perfused, extracted, and embedded in alginate to stabilize tissues for thin slabbing. Coronal slabs (4mm) were generated for each hemisphere, and the cerebellum and brainstem were removed and frozen as a separate sample for each animal. Flash freezing of tissue samples was conducted using dry ice and isopentane as described above for human tissues. Tissues were shipped to the Allen Institute on dry ice and were stored under vacuum seal conditions at -80°C until the time of dissection.

#### Determination of RNA Integrity from Frozen Human and Macaque Brain Tissue

To assess RNA quality, singular tissue samples (roughly 50□mg each) were collected from tissue slabs corresponding to the frontal and occipital pole of each brain. Dissected tissues were stored in 1.5-ml microcentrifuge tubes on dry ice or in −80□°C until the time of RNA isolation. Tissue samples were homogenized using a sterile Takara BioMasher (cat. no. 9791A). RNA isolation was performed using a QIAGEN RNeasy Plus Mini Kit (cat. no. 74134) or a Takara NucleoSpin RNA Plus kit (cat. no. 740984) according to the manufacturer’s protocol. RIN values for each sample were determined using the Agilent RNA 6000 Nano chip kit (cat. no. 5067-1511) and an Agilent Bioanalyzer 2100 instrument according to the manufacturer’s protocol.

#### NIMP Integration and Human and Macaque Tissue Dissections

In collaboration with the BICAN consortium, fresh tissue images captured for all brain specimens were uploaded to the Neuroanatomy-anchored Information Management Platform (NIMP) (Zhang et al., co-submitted)^88^. Slab photos were digitally cropped and then assigned to dissection-specific requests in NIMP, which allowed dissectors to draw dissection plans and assign anatomical pins linking drawn regions of interest (ROIs) with neuroanatomical structures defined in our HOMBA ontology (https://alleninstitute.github.io/CCF-MAP/docs/HOMBA_ontology_v1.html). Tissue was dissected using dry-ice-chilled razor blades and forceps on a custom platform maintained at -20□°C, and dissections were documented using a custom blockface imaging system. Blockface tissue sections were captured from all tissue blocks prior to dissection of specific ROIs. Tissue blocks containing several ROIs were removed from the parent tissue slab and mounted onto a cryostat sectioning chuck. A series of 10 sections cut at 10µm were collected for each tissue block. This parent tissue was then returned to the dissection platform, removed from the chuck, and individual ROIs were dissected for downstream nuclei isolation. Final ROI blocks were stored in 1.5-ml microcentrifuge tubes on dry ice or in −80□°C until the time of nuclei isolation.

#### Human and Macaque Sample Processing for 10x Genomics Multiome

Single-nucleus suspensions were generated using a previously described standard procedure (https://dx.doi.org/10.17504/protocols.io.5qpvok1exl4o/v2). Briefly, after tissue homogenization, isolated nuclei were stained with primary antibodies against NeuN (FCMAB317PE, 1:100 dilution, Sigma Aldrich) to label neuronal nuclei and OLIG2 (Abcam ab225099, 1:400 dilution) to distinguish non-neuronal populations. Nucleus samples were analyzed using a BD FACSAria II or FACSAria Fusion flow cytometer (software BD Diva v.8.0, BD Biosciences); nuclei were sorted using a standard gating strategy to exclude multiplets. A defined mixture of neuronal (70% from the NeuN+ gate) and non-neuronal (20% from the OLIG2− and 10% from the OLIG2+ gate) nuclei was sorted for each sample. After sorting, nuclei from all sorted populations were combined and concentrated by centrifugation. Chip loading and postprocessing to generate sequencing libraries were done with the Chromium Next GEM Single Cell Multiome Gene Expression kit according to the manufacturer’s guidelines. Nuclei concentration was calculated either manually using a disposable hemocytometer (DHC-NO1, INCYTO) or using the NC3000 NucleoCounter. All 10X libraries were sequenced according to the manufacturer’s specifications on a NovaSeq 6000 using either a NovaSeq X or S4 flow cell. Reads were demultiplexed to FASTQ files using BCL Convert (v.4.2.7) for libraries run on NovaSeq X flow cells and bcl2fastq (v.2-20-0) for libraries run on S4 flow cells. Reads from snRNA-seq libraries were mapped to the 10X Genomics official human reference (GRCh38-2020-A); UMIs per gene were counted using the Cell Ranger (v.6.1.1) pipeline with the --include--introns parameter included. Reads from the snATAC-seq and snMultiome libraries were mapped to the same reference using Cell Ranger ATAC (v.2.0.0) and Cell Ranger Arc (v.2.0.0) pipelines, respectively, with default parameters.

#### Marmoset Tissue Processing for 10x Genomics Single-Nucleus Multiome

Marmoset single-nucleus data was collected as described in the marmoset subcortex study to be published concurrently (Dan, Turner, DeBerardine et al. 2025). Briefly, for 4 animals (2 males, 2 females, aged 2-6 years), 2 mm-thick coronal slabs were frozen, and subcortical regions were microdissected into tiles of 14 to 60 mg. Within this size range, tiles were cut to maintain continuity of structures where possible. After nucleus isolation and labeling with NeuN and Olig2 antibodies, sorted nuclei were pooled to target a ratio of 70% NeuN positive, 10% Olig2 positive, and 20% double negative. After centrifugation and resuspension in nuclei buffer, nuclei were quantified and diluted for 10x Chromium loading. GEM generation and library preparation followed the 10x Multiome protocol, and libraries were quality-checked and sequenced on an Illumina NovaSeq X Plus 25B flow cell at 120,000 reads per nucleus.

#### Quality Control of snRNA-seq data

We performed multi-stage quality control (QC) across species when constructing the consensus taxonomy. For each species, we applied basic cell-level QC to remove low quality profiles. Cells with fewer than 1,000 detected genes or 2,000 UMIs were excluded, and cells with abnormally high gene or UMI counts were removed as well. After an initial round of clustering and label-transfer annotation, we performed a label-aware QC pass at the cluster-level. Clusters with low median gene or UMI counts, elevated doublet scores, or highly heterogeneous broad annotation profiles were removed. In addition, once major neuronal and non-neuronal identities were established, we applied a stricter neuronal filter by excluding neurons with <2,000 detected genes. Minor variations in thresholds were applied across species to account for differences in data characteristics.

Next, high-quality cells from all species were aligned in a shared embedding space. Clusters showing poor cross-species alignment were manually reviewed based on multiple criteria, including within-species embedding structure, species-mixing patterns in the aligned space, and expression patterns of marker genes. Clusters that failed alignment criteria were excluded. Finally, evidence from spatial transcriptomics and additional multiomic assays was used to further corroborate cluster identity, assess anatomical consistency, and refine cross-species of QC decisions.

#### Clustering of snRNA-seq data

Clustering was performed using a Python implementation of the scrattch.hicat iterative clustering package (https://alleninstitute.github.io/scrattch.hicat/). To minimize donor-specific variation prior to clustering, we generated a single donor-corrected latent representation (via scVI) and used this corrected embedding throughout the entire iterative procedure. No additional donor-level adjustment was performed during downstream rounds of subdivision.

We applied the iter_clust automatic top-down clustering function which recursively partitions cells into increasingly fine transcriptomic groups. This algorithm identifies new clusters only when all resulting subclusters are mutually separable by stringent differential gene expression (DGE) criteria. Following Yao et al. (2023), we used parameters optimized for detecting small but transcriptionally distinct populations:

- q1.th = 0.5 (minimum fraction of expressing cells in the candidate foreground cluster),
- q.diff.th = 0.7 (minimum normalized expression-fraction difference distinguishing foreground and background),
- de.score.th = 150 (minimum cumulative differential-expression score), and
- min.cells = 4

These criteria ensure that any pair of final clusters is separated by at least 8 binary DEGs. Each DEG’s contribution to de.score is capped at 20, so a minimum of eight genes is required to exceed the threshold of 150. Binary DEGs were defined as genes expressed in ≥40% of cells in the foreground cluster, with |log2 fold-change| > 1, adjusted P < 0.01, and a foreground-normalized expression-fraction difference > 0.7.

#### Replicability of Group-terms across species with MetaNeighbor

Group-level replicability across species was evaluated using MetaNeighbor^44^, which quantifies how well cell type labels can be recovered using cross-dataset classification. Expression matrices were restricted to the conserved marker genes, and transcriptomic Groups were used as labels. For one-vs-best replicability, MetaNeighbor identifies, for each target and reference species pair, the two closest matching transcriptomic Groups and computes an AUROC based on the voting results, comparing cells from the closest match to cells from the second closest match. The resulting AUROC values reflect how distinctly each Group can be identified across datasets in a local context. High one-vs-best AUROC scores (>0.7) indicate strong Group replicability, whereas lower values highlight Groups with weaker cross-dataset correspondence.

#### ATAC-seq processing, QC, and Annotation

Single-nucleus ATAC-seq data from 10x Genomics multiome libraries were processed with snapatac2 ^89^ (version 2.8). For each sample, we imported fragment files into an on-disk AnnData object using *snap.pp.import_fragments* with appropriate reference (fasta, GTF, and chromosome sizes). We restricted analysis to nuclei that were also present in the matched RNA-seq dataset, enabling direct transfer of transcriptomic labels (Neighborhood, Class, Subclass, Group, and cluster ID) from the RNA to the ATAC modality on a per-nucleus basis via shared barcodes.

Quality control was performed in snapatac2 using TSS enrichment, nucleosome signal, and fragment counts as key metrics. TSS enrichment was computed with *snap.metrics.tsse*, and nuclei were retained if they had TSS enrichment ≥ 5 and ≥ 1,000 fragments (*snap.pp.filter_cells(min_tsse=5.0, min_counts=1000*).

To generate bigWigs, we concatenated all high-quality nuclei into a single snapatac2 dataset and computed coverage grouped by transcriptomic Group. TSS regions were defined using GENCODE annotations: all “transcript” entries in the GTF were converted into ±100 bp windows around the transcription start site, and overlapping windows were collapsed to a non-redundant set of TSS intervals. These TSS intervals were supplied to *snap.ex.export_coverage* and coverage was exported in 10 bp bins using insertion counts and CPM scaled by TSS reads, such that each bin value equaled *(reads in bin / total reads in TSS intervals) × 10*_. This procedure yields cross-sample and cross-Group bigWigs normalized to TSS-proximal signal.

Peak calling was performed in snapatac2 via MACS3 using default parameters, applied per Group (*snap.tl.macs3*). Group-specific peak sets were then merged across Groups and chromosomes using *snap.tl.merge_peaks* to generate a unified, non-redundant peak catalog. A peak-by-cell count matrix was built from this merged peak set with *snap.pp.make_peak_matrix*, and the top 250,000 most variable peaks were selected as features (*snap.pp.select_features*).

For dimensionality reduction and donor correction, we computed a spectral embedding (*snap.tl.spectral*) and UMAP visualization (*snap.tl.umap*) on the peak matrix. Donor identities were mapped to each nucleus by parsing sample IDs from cell barcodes and joining to external metadata. To control for donor-specific effects, we applied Harmony batch correction on the spectral embedding (*snap.pp.harmony(batch=“donor”)*) and computed UMAP using the Harmony-corrected representation. This donor-corrected latent space was used for all downstream ATAC-based visualizations.

#### Annotation transfer with MapMyCells

Transcriptomic annotations were transferred onto the HMBA BG dataset using MapMyCells (RRID:SCR_024672) implemented in the Python package cell_type_mapper (*AllenInstitute/cell_type_mapper*). Briefly, a curated reference atlas with taxonomic labels was used as the training reference, and HMBA BG cells were treated as the query. MapMyCells then computed gene-level similarity scores and performed both correlation and hierarchical label transfer, assigning each HMBA cell the most likely reference label at multiple taxonomic levels (e.g., Class, Subclass, Group). The method returns both discrete labels and confidence scores, which we used to filter low-confidence mappings and to refine downstream consensus annotations.

#### Alignment of whole-brain RNA-seq studies

Whole-brain reference datasets were aligned to the HMBA BG taxonomy using scVI ^90^. For each whole-brain reference, we first restricted the data to BG-relevant populations by iteratively removing clusters from non-BG regions based on published annotations and spatial transcriptomics. Gene expression matrices were harmonized by converting species-specific gene symbols to 1:1 orthologs and retaining only shared genes. We then selected the top 3,000–4,000 highly variable genes per dataset and trained scVI models using species as the batch variable and donor identity as an additional covariate. The resulting scVI latent space was used to compute neighbor graphs, UMAP embeddings, and clustering, providing a unified cross-species representation for downstream label transfer and homology analysis.

#### Differential abundance analysis with Milo

Differential abundance (DA) between HMBA BG neurons and the whole-brain references was assessed using Milo implemented in pertpy ^91^ from the scverse. Analyses were performed in the aligned scVI latent space, which provided the neighborhood graph for Milo. HMBA consensus Group labels were first propagated to all whole-brain reference cells using a balanced kNN–majority-vote procedure. We constructed a kNN graph (150 neighbors) on the scVI embedding and generated overlapping neighborhoods (prop=0.1). Milo counted cells per neighborhood using sequencing library as the sample identifier and encoded dataset origin. DA was tested using a negative binomial model (design=“∼study”, solver=“pydeseq2”), producing neighborhood-level log fold changes and SpatialFDR values. Neighborhoods were annotated by their dominant Group label; those with <60% agreement were marked “Mixed.” Significant DA was defined as SpatialFDR < 0.1 and visualized on the neighborhood graph and via beeswarm/violin plots summarizing logFC distributions across Groups.

#### Expressologs

We quantified cross-species conservation of gene expression patterns, expressologs ^47^, using pseudobulk mean expression matrices for human, macaque, and marmoset for each annotation level (Neighborhood, Class, Subclass, Group) and shared set of 1:1 orthologous genes. Cell type labels were harmonized by taking the union of all observed cell types; for any missing cell type in a given species, a zero-filled row was added to ensure that all matrices were aligned and identically ordered. For each pairwise comparison, we designated one species as the reference and the other as the query. Then for every gene in the shared ortholog set, we computed the Pearson correlation between its pseudobulk expression profile across matched cell types in the reference and query species. We also recorded the standard deviation of each gene’s expression profile in each species as a measure of variability. This process produced a symmetric set of gene-wise expression correlations for all primate species pairs (reference→query and query→reference).

To derive expressolog scores for 1:1 orthologous gene pairs, we standardized these pairwise correlations using a rank-based AUROC procedure.

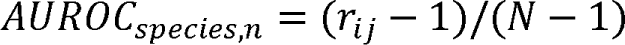

where N is the total number of genes, and r_ij_ is the rank of the Pearson correlation of gene i with its 1:1 ortholog gene j relative to all other (N□−□1) genes in the second species.

For each gene, we computed the median cross-species expressolog score across ortholog comparisons.

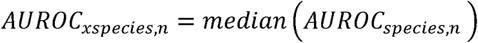

The expressolog scores were also calculated across taxonomy levels using the same formula and captures the extent to which gene expression variation is shared across species.

#### Cell type specificity metric

We quantified gene-level cell type specificity across species using a Tau-based meta-analysis of snRNA-seq expression profiles^92^. For each species (human, macaque, marmoset), we used AnnData objects containing donor-resolved cell-by-gene count matrices and consensus annotations (“Neighborhood”, “Class”, “Subclass”, “Group”). To obtain robust cell type-level estimates while accounting for donor heterogeneity, we performed repeated donor subsampling: for each species, in each iteration we randomly selected a subset of donors (number of donors = max[2, ceil(total_donors/3)]) and restricted the analysis to cells from those donors. Within each subsample, we aggregated expression per annotation level using scanpy utilities to compute mean expression for each gene in each cluster. Genes with mean expression > 1 in at least one cell type were retained, and the resulting gene × cell type matrix was used to compute Tau scores.

Tau was defined for each gene as:

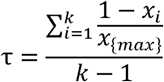

where (x_i) is the mean expression in cluster (i), (x_{\max}) is the maximum mean expression across the (k) clusters, and genes with zero maximal expression were assigned Tau = 0. This procedure was repeated for 10 random donor subsamples, yielding a matrix of Tau values (one column per subsample) for every gene. We then summarized the mean Tau across subsamples (“mean_tau”) and used this value for downstream analysis.

#### Differential expression analysis

Differential gene expression analysis was performed using scanpy.tl.rank_genes_groups (Scanpy version 1.10.1) with the Wilcoxon rank-sum test and Benjamini-Hochberg correction (method=“wilcoxon”, corr_method=“benjamini-hochberg”). As an illustrative example, to compare D2 StrioMat cells with other D2 MSNs across species, we first restricted each dataset to D2-type MSNs and annotated cells as either D2 StrioMat or non-StrioMat D2 MSN. DEG analysis was performed independently for each species using the settings above. Genes were considered enriched in D2 StrioMat cells if they met predefined thresholds for statistical significance and effect size (adjusted P < 0.01, log2fold-change > 1, and detection in >20% of cells in the target group). Species-specific DEG lists were then intersected to identify markers conserved across human, macaque, marmoset, and mouse.

#### Cactus sequence alignment

In order to prepare the 447-way Cactus alignment for use with the specific genome assemblies used with our genomics data, the replace utility of the Cactus-update-prepare was used for the macaque, marmoset and mouse genomes.

Cross-species orthologs of ATAC-seq peaks were identified using the Zoonomia 447-mammal Cactus HAL alignment ^93^. For each species, merged peak BED files were projected across the Cactus alignment using HALPER ^94^ and halLiftover (*pfenninglab/halLiftover-postprocessing*), which identify orthologous intervals in other genomes while enforcing alignment-quality constraints.

We used the following HALPER parameters to ensure high-confidence mappings:

- MIN_LEN = 50: minimum aligned length required for an orthologous interval
- PROTECT_DIST = 10: distance flanking the query interval preserved during alignment to avoid edge truncation
- MAX_FRAC = 2: maximum allowed fractional expansion of the orthologous region relative to the input peak (i.e., up to 2× peak length)

Peak BED files were mapped across all species in the 447-way HAL, and HALPER postprocessing steps were applied to merge, filter, and classify orthologous intervals. The resulting tables provided, for each input peak, the set of confidently aligned orthologs across species and ancestral nodes, enabling downstream evolutionary and conservation analyses.

#### Definitions of ortholog and species-specific alignments

Using the output of HALPER on the 447-way Cactus alignment, we defined orthologous cCREs among the three profiled primate species (human, macaque, marmoset) in a human-anchored framework. We started from merged human ATAC cCREs and HALPER narrowPeak files that align macaque and marmoset peaks into human (hg38) coordinates. Human peaks were given unique genomic IDs (chr:start–end) and converted to PyRanges; for each non-human species, we filtered HALPER alignments to retain only intervals with ≥250 bp aligned to the human peak (∼50% of a 500-bp cCRE), corrected HALPER-set peak IDs, and, for marmoset, harmonized chromosome names using a RefSeq–UCSC alias table. We then intersected species-specific HALPER tracks with the human cCRE set to assign, for each human region, the corresponding macaque and marmoset peak IDs and aligned human coordinates. This produced a human-anchored liftover table from which we classified species-specific cCREs as those lacking matches in both other species, and orthologous cCREs as those with aligned peaks in all three species.

#### Definition of cCRE evolutionary distance in Cactus alignment

We quantified evolutionary conservation of ATAC-seq peaks across mammals using a phylogenetic tree derived from the 447-way Zoonomia Cactus HAL alignment. For each peak, we computed whether ≥90% of bases were aligned to each species or ancestral node, producing a binary alignment matrix. The Zoonomia phylogeny was parsed in Newick format, and branch lengths were extracted for all internal and external nodes. Branch lengths were normalized to sum to one, providing a relative evolutionary distance where 0 indicates no conservation (species-specific) and 1 indicates alignment to all 447 species. For each peak, we calculated an evolutionary distance score by summing normalized branch lengths across all nodes to which the peak was ≥90% aligned. This yielded a continuous measure of evolutionary conservation that integrates both breadth (number of aligned species/ancestors) and phylogenetic depth (distance along the tree).

#### Annotation of transcription factor motifs underlying cCREs

We converted merged ATAC-seq peak BED files into FASTA to obtain underlying genomic sequences. Peak coordinates were standardized to Chromosome, Start, and End, converted to a PyRanges object, and used with pyfaidx-indexed reference genomes (hg38, mm10, rheMac10, calJac4) to extract sequences while preserving peak order; macaque chromosomes were harmonized via an alias table. For peaks mapping to missing contigs, we substituted 500-bp N-masked sequences and wrote all sequences to a peak-resolved FASTA file with headers of the form >chr:start-end for downstream motif scanning.

We scanned peak-derived FASTA sequences for transcription factor motifs using FIMO via the *memelite* ^95^ python interface, applying the JASPAR2024 CORE vertebrate non-redundant ^96^ motif set in MEME format. FIMO was run directly on the peak FASTA file, and all motif hits were concatenated into a single results table containing the matched TF motif, genomic position, score, p-value, and originating peak. TF identifiers were parsed from motif names, TF families were annotated from TFClass ^97^, and hits were aggregated into a peak × transcription factor count matrix by tallying all motifs detected within each peak. This matrix was stored as an AnnData object for downstream analyses, including clustering, enrichment summaries, and integration with chromatin accessibility features.

#### cCRE annotation

We annotated BG ATAC cCRE for overlap with transposable elements (TEs) using species-specific RepeatMasker tracks. For each species, merged peak BED files were converted to PyRanges objects and given a unique peak ID (chr:start-end). Species-appropriate RepeatMasker annotations were loaded in UCSC rmsk schema (hg38, rheMac10) or parsed from NCBI RepeatMasker output (marmoset), with assembly-specific chromosome aliases harmonized for macaque and marmoset. Repeat records were converted to PyRanges and intersected with peaks using a genomic join. For each peak-TE overlap, we retained the repeat name (repName), class (repClass), and family (repFamily), and required a minimum overlap of 250 bp (∼50% of a 500-bp peak); overlaps below this threshold were treated as non-TE. cCREs lacking qualifying overlaps were labeled with False and -1 for TE and repeat fields. The resulting TE-annotated peak tables were saved per species for downstream analyses.

We annotated cCREs for promoter overlap using species-specific GENCODE gene models. For each species, merged cCRE coordinates were converted to PyRanges objects with unique region identifiers. Promoter intervals were derived from GTF annotations by extracting protein-coding gene entries, computing strand-aware transcription start sites (TSS), and expanding these to asymmetric promoter windows (–1 kb / +500 bp on the positive strand; –500 bp / +1 kb on the negative strand). Species-specific chromosome aliases were applied for macaque and marmoset. cCREs were intersected with promoter intervals using genomic joins, and each region was assigned a binary promoter label based on whether it overlapped any promoter interval. For each species, promoter-annotated cCRE tables were exported for downstream analyses.

We quantified evolutionary conservation for human and cross-species orthologous cCREs using phyloP 100-way vertebrate scores. Human cCREs along with macaque and marmoset orthologous cCREs in hg38 coordinates were combined into a unified set of species-cCRE tables. For each cCRE, we retrieved base-resolution phyloP scores from the hg38 phyloP100way bigWig using pyBigWig. We queried each cCREs chromosome, start, and end coordinates; extracted the corresponding phyloP vector; removed missing values arising from unaligned or low-coverage regions; and computed the mean phyloP score cCREs with no valid phyloP positions were assigned NaN.

#### Identification of cell type-specific cCREs

The Gini coefficient provides a single-value summary of how unevenly a cCRE’s accessibility is distributed across cell types. For each cCRE, we treat its pseudobulk accessibility across k cell types as a vector *x* = (*x*_1_,…, *x_k_*) *x* = (*x*_1_,…, *x_k_*). The Gini coefficient measures inequality in this vector, taking a value of 0 when accessibility is evenly distributed across all cell types and increasing toward 1 as accessibility becomes concentrated in a single cell type. Formally, the Gini coefficient is computed by sorting accessibility values in ascending order, then evaluating for each cCRE:

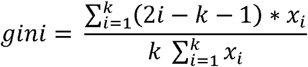

Thus, cCREs with broad accessibility have low Gini scores, whereas cell type-restricted or highly specific cCREs yield high Gini values. In our pipeline, we report for each cCRE the maximum Gini value across cell types (assigned to the cell type with the highest accessibility), which serves as the cCRE’s overall specificity score.

#### Conservation of chromatin accessibility

To integrate sequence conservation with chromatin-state conservation, we defined accessologs, the chromatin accessibility analogue of expressologs. Accessologs quantify whether a sequence-conserved cCRE retains a similar pattern of accessibility across homologous cell types in another species. Using ortholog regions derived from the Cactus multi-species alignment, we computed pseudobulk accessibility for each cCRE across the consensus BG cell types. For every pair of species, we calculated the Pearson correlation between the accessibility profile across matched cell types in the reference and query species. We then rank-standardized the similarity of the true orthologous region relative to all possible region pairs, yielding an AUROC-based accessolog score. High accessolog scores indicate that a conserved sequence element also preserves coordinated accessibility patterns across species, whereas low scores reflect divergence in regulatory activity despite sequence alignment.

#### Human-accelerated genomic element analysis

HARs, hCONDELs, and HAQERs were obtained from previously published datasets^98–100^. We quantified the intersection between human BG ATAC-seq peaks and three classes of human-evolved regulatory regions (HARs, hCONDELs, and HAQERs) using PyRanges.join to generate peak-level overlap tables for each variant class. Redundant peak entries mapping to the same genomic interval were removed, and downstream analyses were restricted to variants with at least one overlapping accessible peak. To assign putative target genes to HAQER-overlapping peaks, we intersected these overlaps with an activity-by-contact (ABC) enhancer–gene prediction dataset generated for human BG (hg38). ABC predictions were encoded as genomic intervals with associated gene IDs, and only enhancer–gene links with ABC score > 0.02 were retained. Joining HAQER–peak overlaps with this filtered ABC set yielded the set of genes whose regulatory landscapes include at least one HAQER-associated accessible peak. We then compared these genes to a curated list of human-specific differentially expressed genes (hDEGs) derived from cross-species snRNA-seq analyses of BG cell types. Genes appearing in both the ABC-linked HAQER gene list and the hDEG catalogue were annotated as human-specific. At the variant level, each HAQER was flagged as hDEG-associated if any of its ABC-linked genes overlapped the hDEG set.

#### Developmental cell type correspondences

We used the list of cell identity TFs identified in the adult mouse whole brain, plus several key missing genes (Supplementary Table 1.3) ^17^. We mapped developmental initial classes from a three-species developing whole-brain atlas to adult Groups by maximum Pearson correlation using the log gene expression values of core identity genes.^101^

#### DNA sequence model training and analysis

For each species, we trained a model using the default dilated CNN definition provided in the CREsted package (Kempynck et al., 2025). This architecture employs an initial convolutional block followed by a series of dilated convolutional layers with residual connections for peak regression. Training was performed using a custom wrapper that enabled TensorFlow distributed strategies for multi-GPU execution, with all core training logic handled by crested.tl.Crested. Cell type-specific accessibility targets were generated by aggregating snATAC-seq profiles per cell type within each species. The aggregated peak signal was normalized across groups using crested.pp.normalize_peaks() on the top 3% of peaks per group. Data were partitioned by chromosomes: chromosome 10 was assigned for validation, chromosome 18 for testing, and the remaining chromosomes for training. All models followed the default peak-regression training strategy and configuration in CREsted. Pretraining used all consensus peaks extended symmetrically on both sides of each summit to a 2,114-bp window, a learning rate of 0.001, and the CosineMSELogLoss loss. Models were then fine-tuned on variably accessible regions defined by crested.pp.filter_regions_on_specificity(), using a reduced learning rate of 1 × 10□□.

We computed a two-dimensional UMAP embedding of regions based on sequence representations extracted from the respective species CREsted model. The top 100 most specific regions per cell type were selected using *crested.pp.sort_and_filter_regions_on_specificity()* with method=”proportion”. Per-region model embeddings were extracted from the *global_average_pooling_1d* layer via *crested.tl.extract_layer_embeddings*(). The embedding dimensions were then reduced to 2-dimensions using the UMAP python package with default parameters.

We quantified group correspondence across species by correlating model predictions on matched, highly specific region sets and visualized per-cell-type correlations across species pairs. For each species we selected the top 500 most specific regions per cell type using *crested.pp.sort_and_filter_regions_on_specificity(method=“proportion”)*. For each species’ region set, we computed predictions with each species model (*crested.tl.predict*) to obtain cross-prediction tensors keyed by source species and cell type. For every pair of species (Macaque–Human), we formed celltype-by-celltype correlation matrices by comparing per-region prediction vectors between the two models, using Spearman correlation on log1p-transformed values; we also computed a “combined” matrix by concatenating both directional comparisons. We extracted the diagonal (same-name cell types) to summarize per-cell-type cross-species agreement.

#### TF motif analysis

We derived cell type-specific sequence patterns for all species with their corresponding CREsted models using the contribution scores of the top 500 most specific regions per celltype. We selected these regions with *crested.pp.sort_and_filter_regions_on_specificity()* on the averaged peak heights and model predictions. For each region, we computed nucleotide-level contribution scores using CREsted’s *contribution_scores_specific()* function, method=’integrated_grad’. We ran *tfmodisco-lite* via *crested.tl.modisco.tfmodisco()* to discover recurrent patterns per group, and matched them to known motifs using the TOMTOM option (motifs from *crested.get_motif.db()*). Discovered patterns were post-processed and merged across groups using *crested.tl.modisco.process_patterns()*.

We matched motifs to TF candidates using the matched snRNA-seq data from the respective species. We obtain pattern-to-motif matches from the motif database using *crested.tl.modisco.find_pattern_matches(q_val_thr=0.05)* and mapped motifs to TFs with *crested.tl.modisco.create_pattern_tf_dict()* using the human-oriented annotation columns. Finally, we constructed a TF by cell type importance matrix with *crested.tl.modisco.create_tf_ct_matrix(min_tf_gex=0.95, importance_threshold=3, filter_correlation=True, zscore_threshold=1.5, correlation_threshold=0.49).* The resulting matrix summarizes, per cell type, TF candidates supported jointly by model derived motif evidence and expression concordance.

#### Predicting cell type activity for BG enhancer-AAV sequences

We used the CRESted library to predict Group-specific activity of candidate enhancer sequences. Validated and candidate enhancers were compiled from Hunker, Wirthlin et al. 2025, and genomic coordinates were parsed from cloned fragment coordinates. For each enhancer, we used the cloned sequence, converted to lowercase, and padded or center-trimmed to a fixed length of 2,114 bp so that the sequence was centered and matched the CRESted model input requirements. Predictions were generated with the DeepHumanBG model. For each enhancer, we obtained a vector of predicted accessibility scores across BG Groups which we used for visualization and analysis.

To quantify how enhancer activity evolves across mammals, we aligned each candidate enhancer to the 447-species Zoonomia Cactus HAL and predicted its Group level accessibility using CRESted. For each enhancer, we used hal2maf to extract the corresponding aligned sequence block from the 447-way HAL alignment. The enhancer’s genomic interval (species, chromosome, start, end, length) was supplied as the reference coordinate. hal2maf returned per-species aligned MAF blocks containing orthologous sequence fragments. MAF blocks were stitched using an AWK postprocessing script to merge adjacent aligned fragments. Stitched sequences were then globally aligned across species with MAFFT (--globalpair --maxiterate 1000). This yielded a species-by-sequence alignment for each enhancer, preserving orthology relationships and internal indel structure.

Aligned sequences were converted to FASTA and normalized to the CRESted input length (2,114 bp) by padding or center-trimming, and by replacing MSA gap characters with “N” to preserve alignment information without introducing model-specific biases. We applied the DeepHumanBG model to each species’ aligned enhancer sequence. For each enhancer-species pair, we generated Group accessibility predictions for its annotated target cell type (e.g., D1 MSN, D2 MSN, Pan-MSN, Cholinergic) by computing the average accessibility over the corresponding cell type indices from the DeepHumanBG model.

#### Disease enrichment analysis

We examined SCZ alongside three psychiatric comparison phenotypes (BD, MDD, and PTSD) for which well-powered, polygenic GWAS data were available. Alcohol use per week was included as a continuous behavioral measure without diagnostic ambiguity. To contrast psychiatric versus neurodegenerative processes, Alzheimer’s disease (AD) was also analyzed due to its distinct etiology. Ancient DNA data, known to be enriched for immune functions, were used as an additional control and to assess gene conservation across BG clusters. GWAS summary statistics were obtained from the most recent and best-powered European-ancestry studies available between March and August 2025 (excluding 23andMe data). Table S6.1 lists sample sizes, the number of genome-wide significant loci, and source publications. Analyses were limited to European ancestry due to the lack of equivalently powered GWAS for other populations.

SnRNA-seq data were processed following Duncan et al. (2025)^36^. Expression values were log-transformed (ln[1 + x]) to reduce outliers and scaled to obtain mean expression per gene per cell type. Only protein-coding genes (NCBI) were retained, excluding unexpressed or ambiguously named genes and those within the MHC region (chr6: 25-34 Mb). Ensembl IDs were mapped to Entrez IDs using org.Hs.eg.db (Bioconductor v3.12.0), with non-unique mappings removed. Gene expression specificity was calculated as the transformed expression of a gene in one cell type divided by its total expression across all cell types, yielding values between 0 and 1.

MAGMA (v1.10) was used for gene and gene-set analysis. SNPs were mapped to genes within ±35 kb, and the SNP-wise mean model was applied using European ancestry LD structure from 1000 Genomes Phase 3. Gene-level P-values were derived assuming the test statistic follows a mixture of independent distributions under the null hypothesis. The MHC, X, and Y chromosomes were excluded to avoid confounding LD patterns. Using MAGMA’s gene-property framework, we tested whether cell type-specific gene expression correlated with GWAS signal strength. Gene-level association z-scores were calculated as zg = probit(1 – Pg), where Pg is the P value of the gene and truncated to ±3/6 SD from the mean (MAGMA defaults). Expression specificity for each cell type was used as an explanatory variable in the regression model:

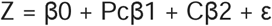

where covariates (C) included gene size, density, sample size, inverse mean minor allele count, and their logarithmic values. We conducted a one-sided test of β₁ to identify cell types enriched for each phenotype (453 clusters tested).

Genome-wide significant SNPs (P < 5 × 10⁻□) were expanded using PLINK v1.9 on the 1000 Genomes Phase 3 (European) reference panel (hg38 build), retaining variants in high LD (r² ≥ 0.8) within 1 Mb. Potential TF motif disruptions were assessed using TOMTOM from the MEME Suite v5.5.8.

#### Dynamic UMAP exploration and differential analysis

To enable interactive exploration and cross-species comparison of the RNA-seq data, we developed a dynamic UMAP explorer project file within the Cytosplore Viewer data visualizer (https://viewer.cytosplore.org). The dynamic UMAP explorer module offers interactive analysis facilities for on-the-fly regeneration and refinement of UMAP and tSNE maps of the underlying data. This allows users to probe the data for potential heterogeneity and detailed inspection of cell subpopulations. Cells can be selected from a taxonomy view, from metadata, or free-form selection in the UMAP. Once two sets of cells are selected, differential expression can be computed on-the-fly based on expression averages per cluster. By approximating gene expression by cluster averages, real-time interaction, searches and visualization for data exploration can be achieved on a wide range of computers. This enables gene expression comparisons within and between species, as well as between user-steered cell populations.

Re-computed embeddings can be colored by cell metadata, or gene painting by clicking on gene symbols in the differential expression tables. Altogether, this enables an interactive, integrated exploration highlighting the specific features of each species and cell cluster in an integrated manner. Cytosplore Viewer can be downloaded from https://viewer.cytosplore.org.

