## Supplemental Table Legends and Figures for "Cross-species consensus atlas of the primate basal ganglia"

Supplemental Tables

**Table S1.1. snRNA-seq cell cluster and Group quality metrics for each species.**

**Table S1.2. Consensus taxonomy hierarchy and Group short names.**

**Table S1.3. Developmental TF relationships and inferred origins.**

**Table S1.4. MetaNeighbor cross-species cell Group classification accuracy.**

**Table S1.5. NS-Forest cell Group markers per species.**

**Table S2.1. Expressolog metric for Group annotations.**

**Table S2.2. Divergent gene lists.**

**Table S3.1. Mouse whole brain mapping to the BG consensus taxonomy.**

**Table S3.2. Human whole brain mapping to the BG consensus taxonomy.**

**Table S3.3. STRd D2 StrioMat Hybrid MSN DEG lists.**

**Table S4.1. Cactus species annotations.**

**Table S4.2. TF-motif annotations.**

**Table S4.3. Genomic variant annotation and ABC results.**

**Table S6.1.** **MAGMA results. Cluster-level associations across brain cell types.**

Gene-set analysis results from MAGMA, testing for associations between genetic risk for each phenotype and gene expression profiles across cell clusters. Each row represents a cell cluster, and columns include the test statistic, p-value. Results are shown for multiple phenotypes and ancestries (SCZ only).

**Table S6.2.** **Effects of genome-wide significant variants on predicted chromatin accessibility.**

Summary of sequencing model analysis for genome-wide significant variants associated with SCZ and Alzheimer’s disease. For each variant, a proximal gene is listed, and the predicted impact on chromatin accessibility is quantified using contribution scores across MAGMA-significant cell types. The Delta of Chromatin Accessibility is reported (predicted accessibility of the reference sequence minus accessibility of the mutated sequence).

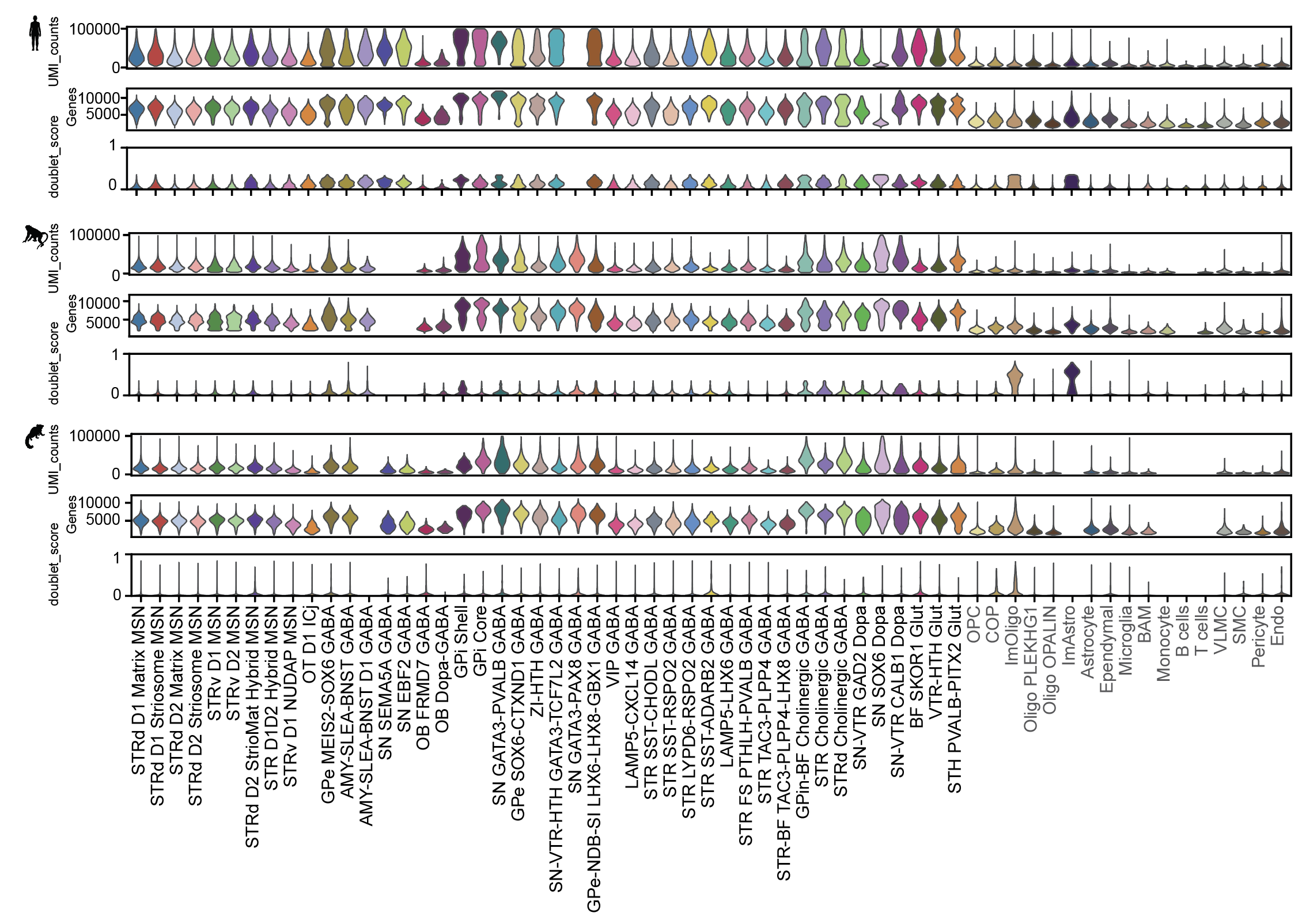

Figure S1.1. BG nuclei quality metrics across cell Groups and species, related to Figure 1.

Violin plots showing median UMI counts, detected genes, and single nuclei doublet scores across Groups and human, macaque, and marmoset.

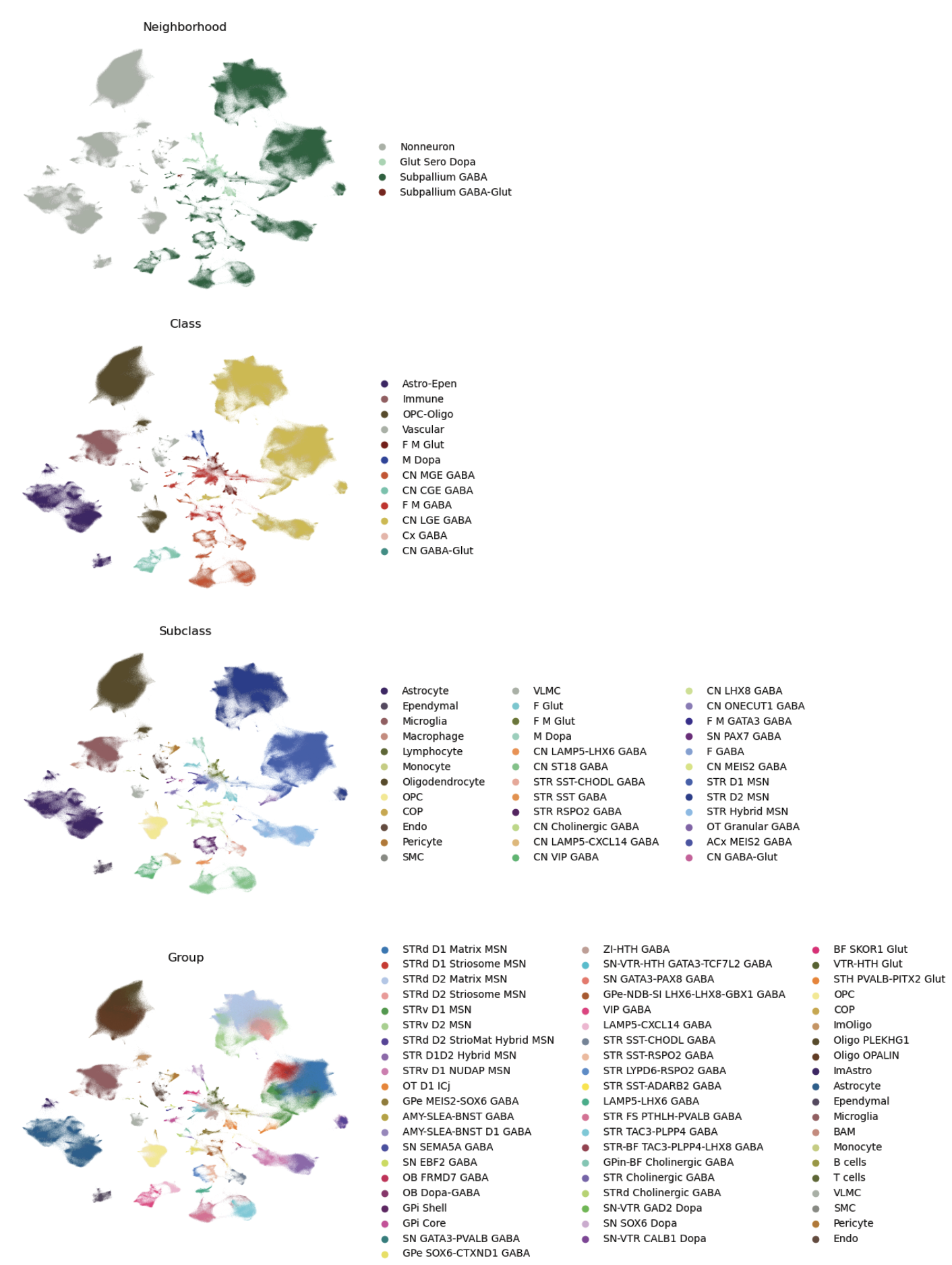

Figure S1.2. UMAP representation of the hierarchical taxonomy, related to Figure 1.

UMAP visualization of the taxonomy at different hierarchical levels (Neighborhood, Class, Subclass, and Group) in the integrated embedding of human, macaque, and marmoset.

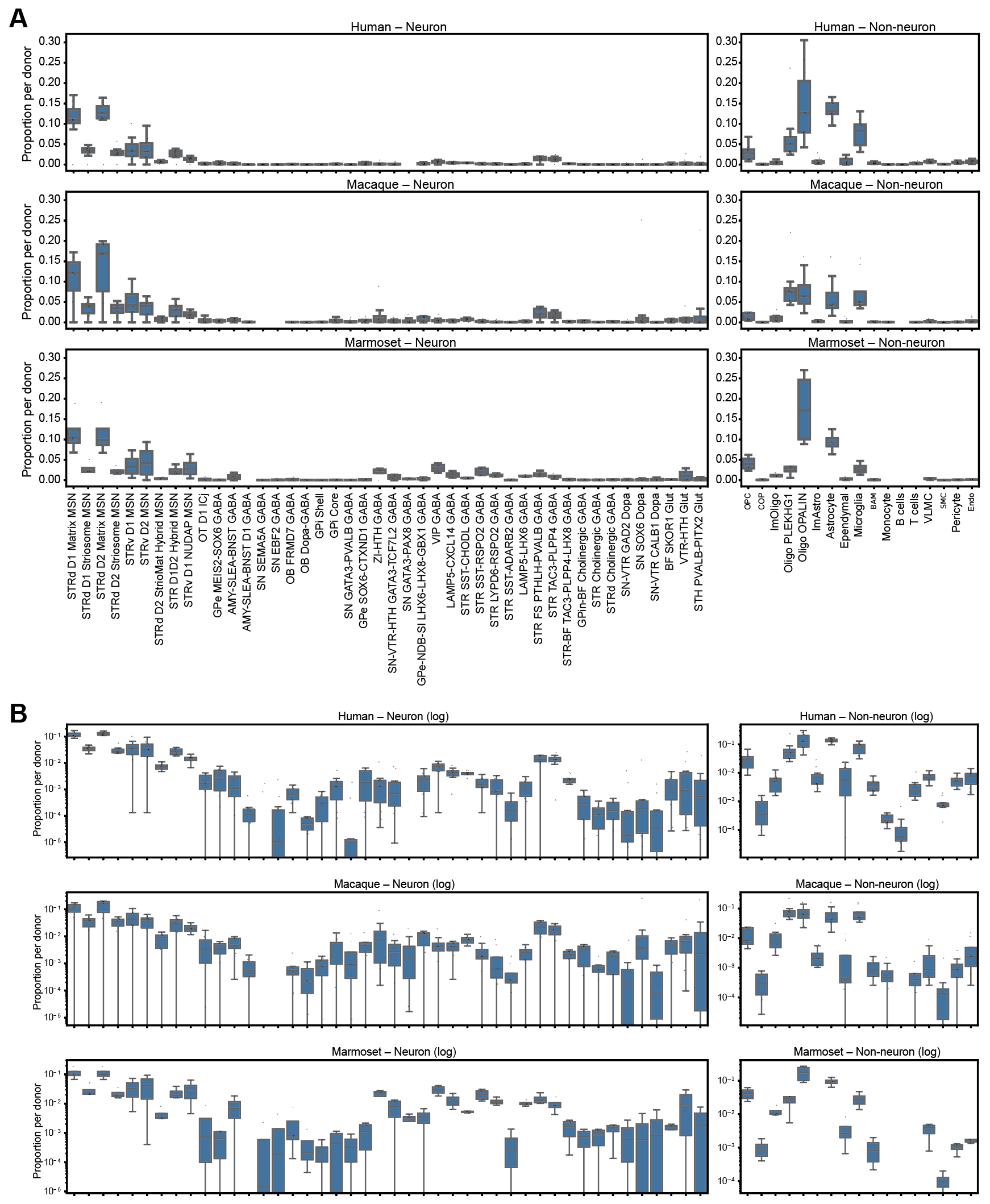

Figure S1.3. Cross-species proportions of consensus Groups across primates, related to Figure 1.

(**A**) Donor-normalized proportions of all consensus Groups for human, macaque, and marmoset. Neuron and non-neuron Groups are calculated and shown separately. Empty x-axis positions indicate Groups absent in each species.

**(B**) Same data are plotted on a log-scale to compare low-abundance neuronal and non-neuronal Groups.

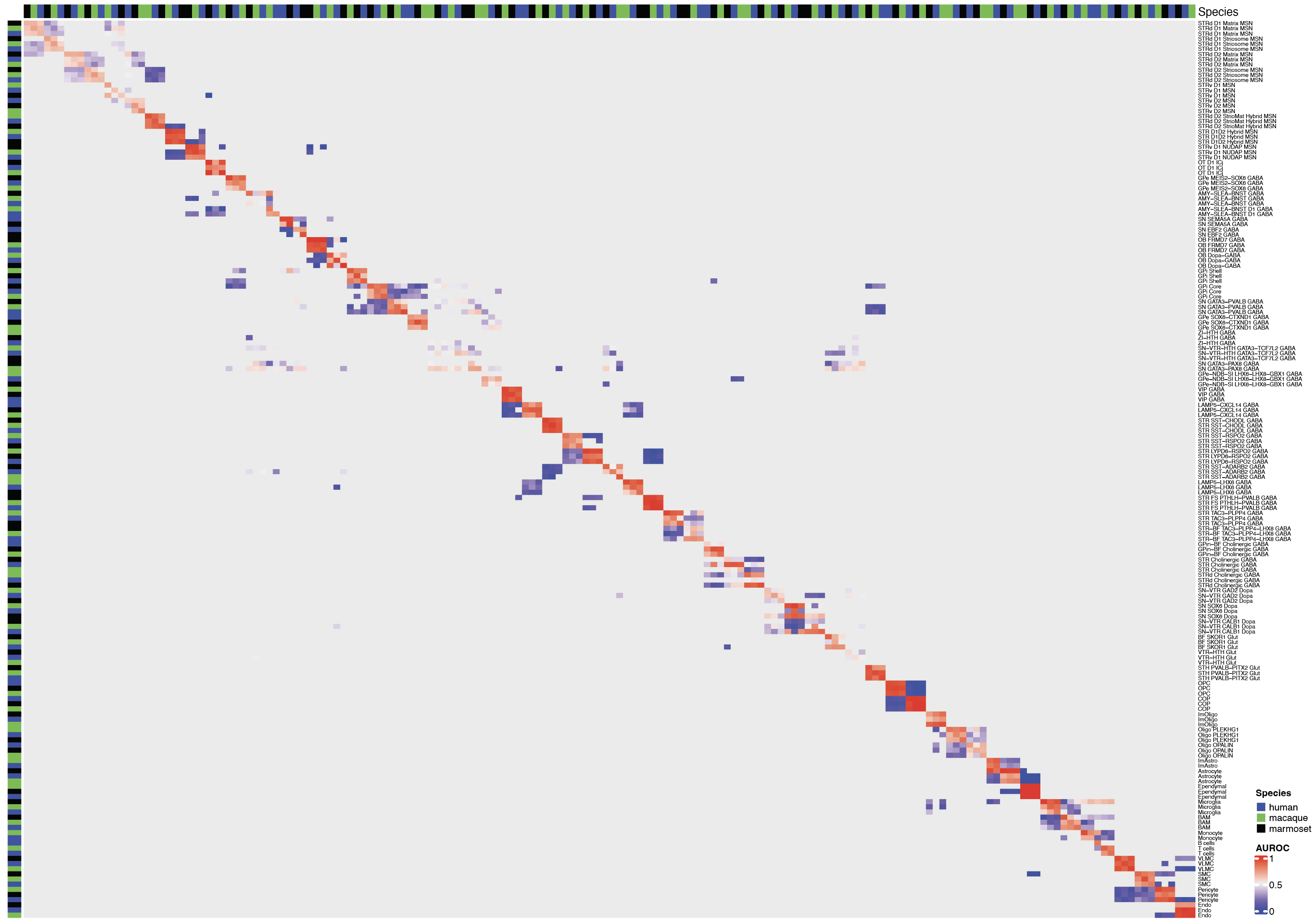

Figure S1.4. Robustness of cell group homologies across primates, related to Figure 1.

MetaNeighbor heatmap of ‘one_vs_best’ classification scores (AUROC) between Group terms across species using the conserved marker gene panel.

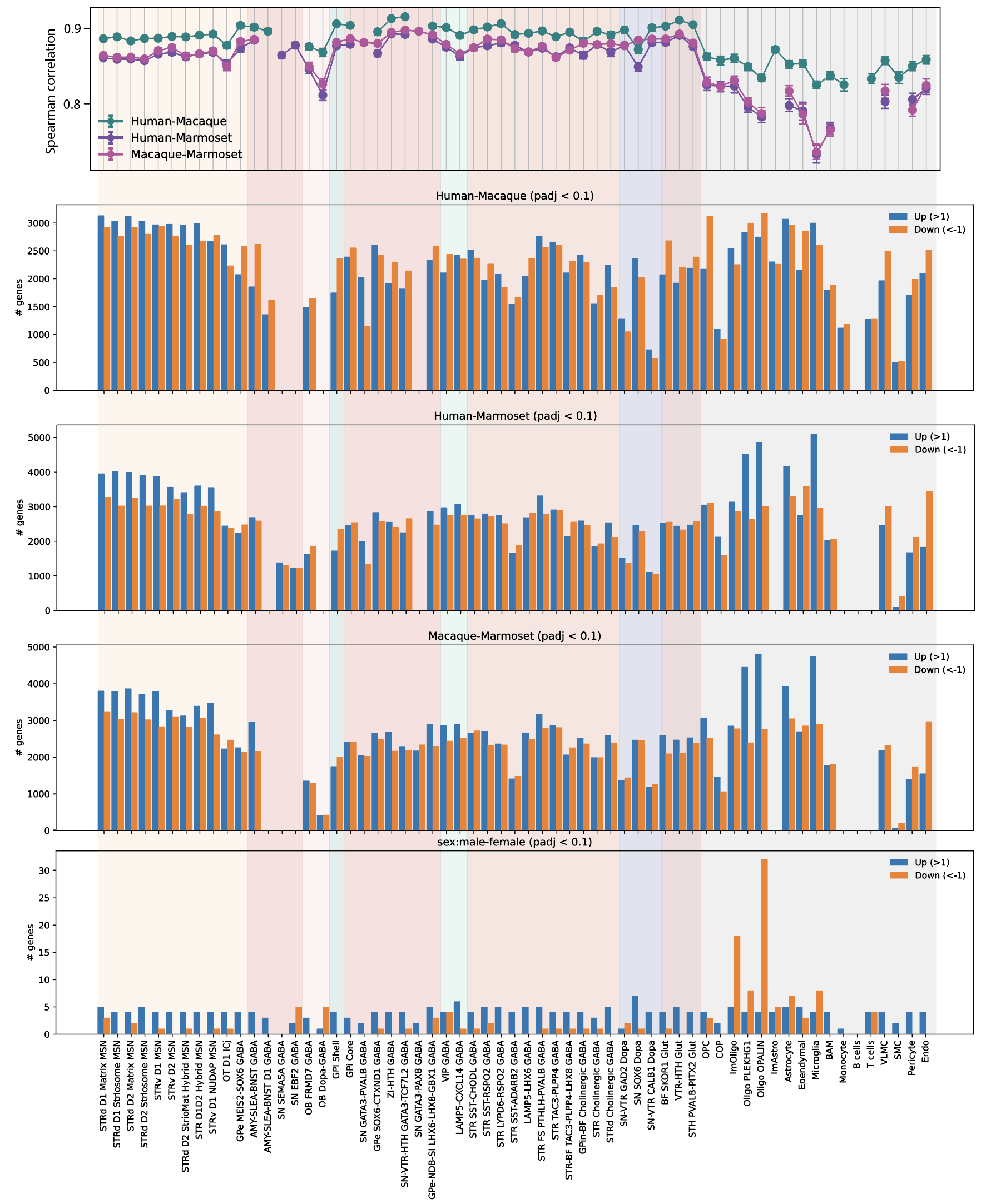

Figure S2.1. Evolutionary divergence of expression varies across cell Groups, related to Figure 2.

Top: Bootstrapped Spearman correlations of expression (mean +/- SD) between pairs of primate species for each Group. Middle: Bar plots of the number of DEGs that are up- (blue) or down- (orange) regulated between each pair of species. Bottom: Bar plots of the number of DEGs that are up- (blue) or down- (orange) regulated between male and female donors across all species.

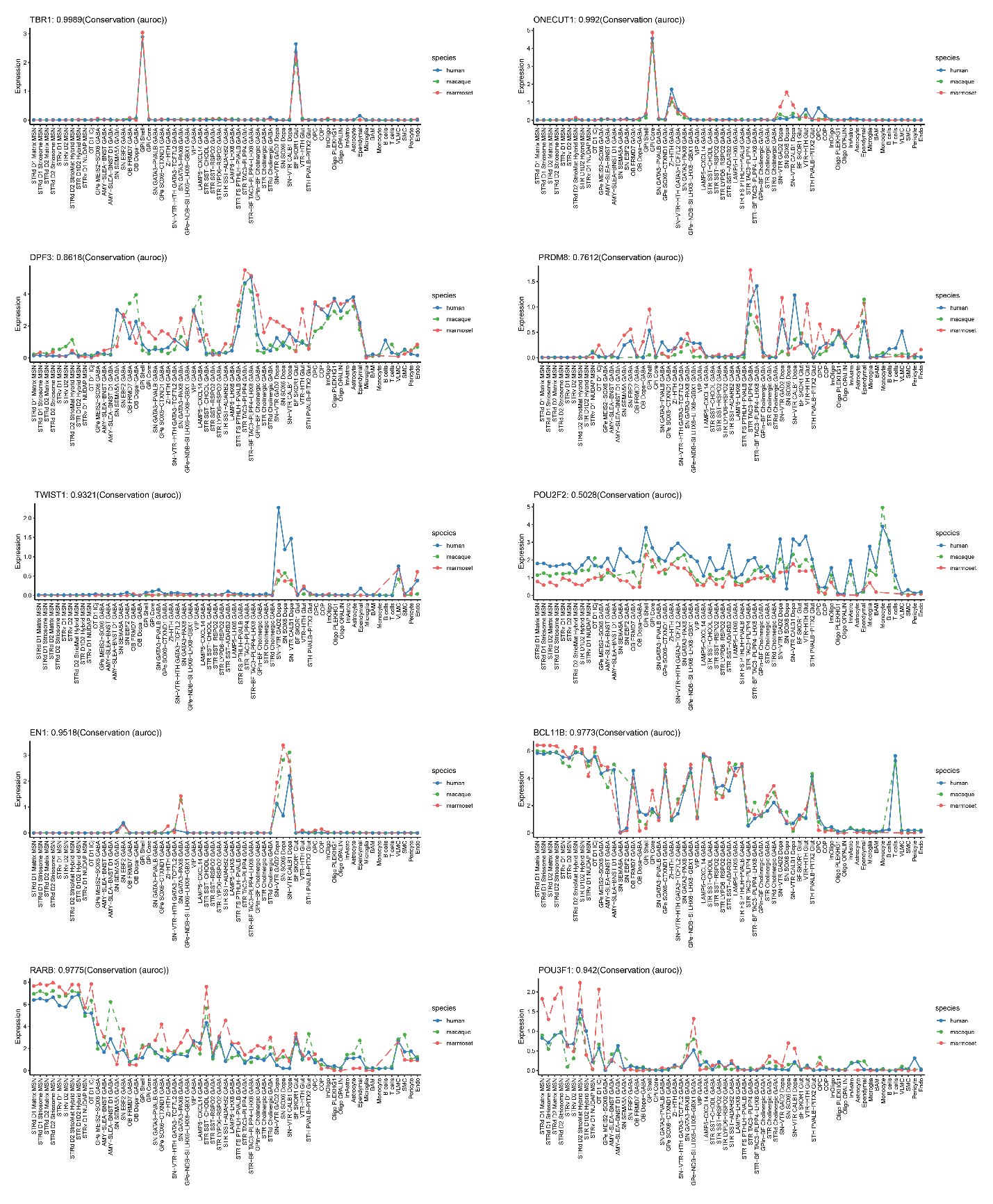

Figure S2.2. TF expression across cell Groups and species, related to Figure 2.

Line plots of mean expression per Group and species. Conservation (AUROC) values were calculated as the mean of pairwise species expressolog scores.

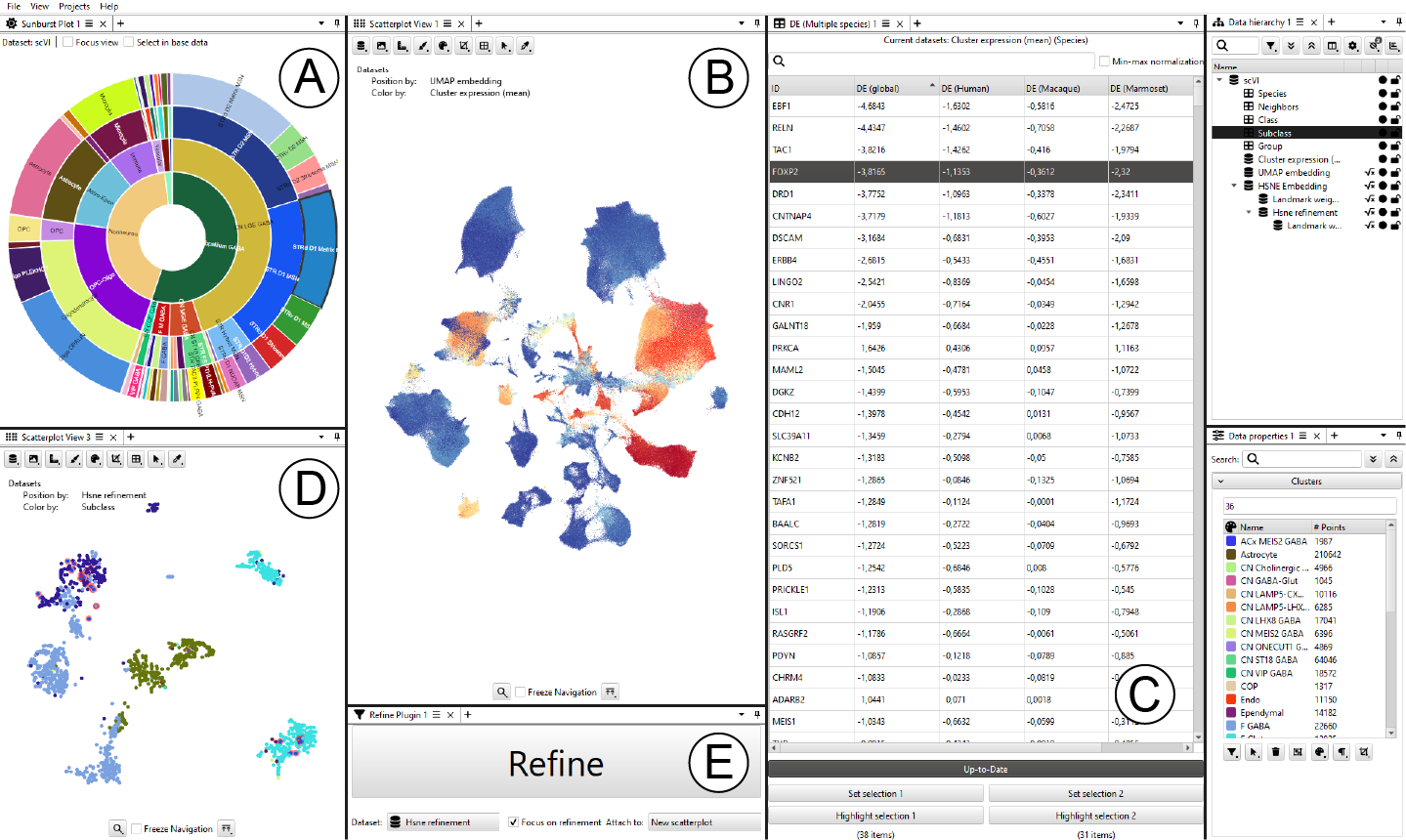
Figure S2.3. Cytosplore software with linked views of the consensus BG taxonomy snRNA-seq data, related to Figure 2.

(**A**) The hierarchical cell type taxonomy for categories that are consistent across species.

(**B**) A UMAP embedding of the integrated scVI snRNA-seq data from all three species, recolored by the mean expression across clusters.

(**C**) Differential expression between two user-defined selections.

(**D**) A refined HSNE embedding for a user-defined region of interest in the center of the UMAP, revealing clearer cluster separation.

(**E**) Main panel for interactively refining new cell selections.

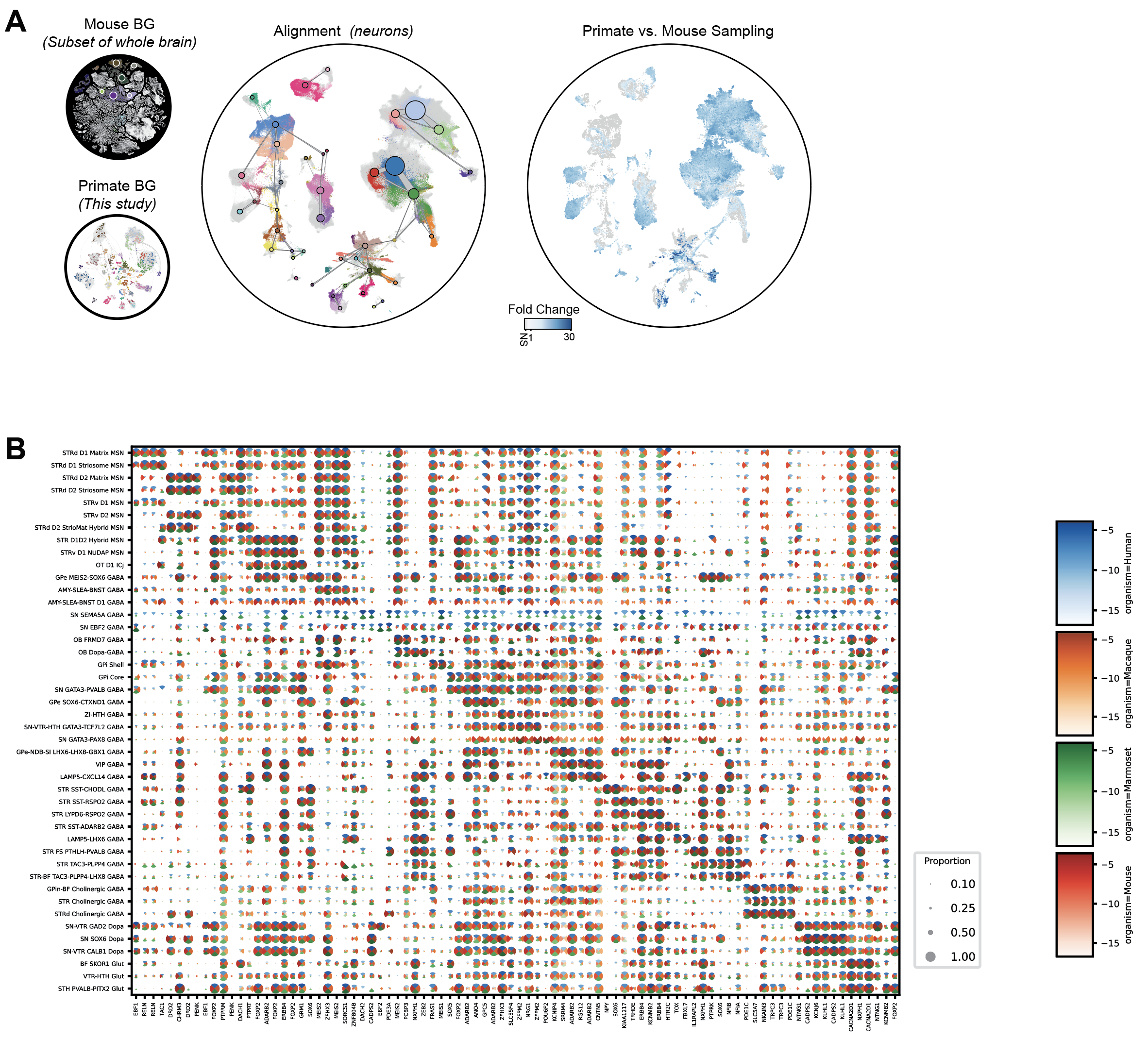

Figure S3.1. Primate and mouse integration defines cell type homologies and conserved markers, related to Figure 3.

(**A**) UMAP visualization of the integrated scVI embedding of primate and mouse BG regions colored by cell group and fold change in cell sampling in primates versus mouse.

(**B**) Expression of markers shown in Figure 3A across the four species.

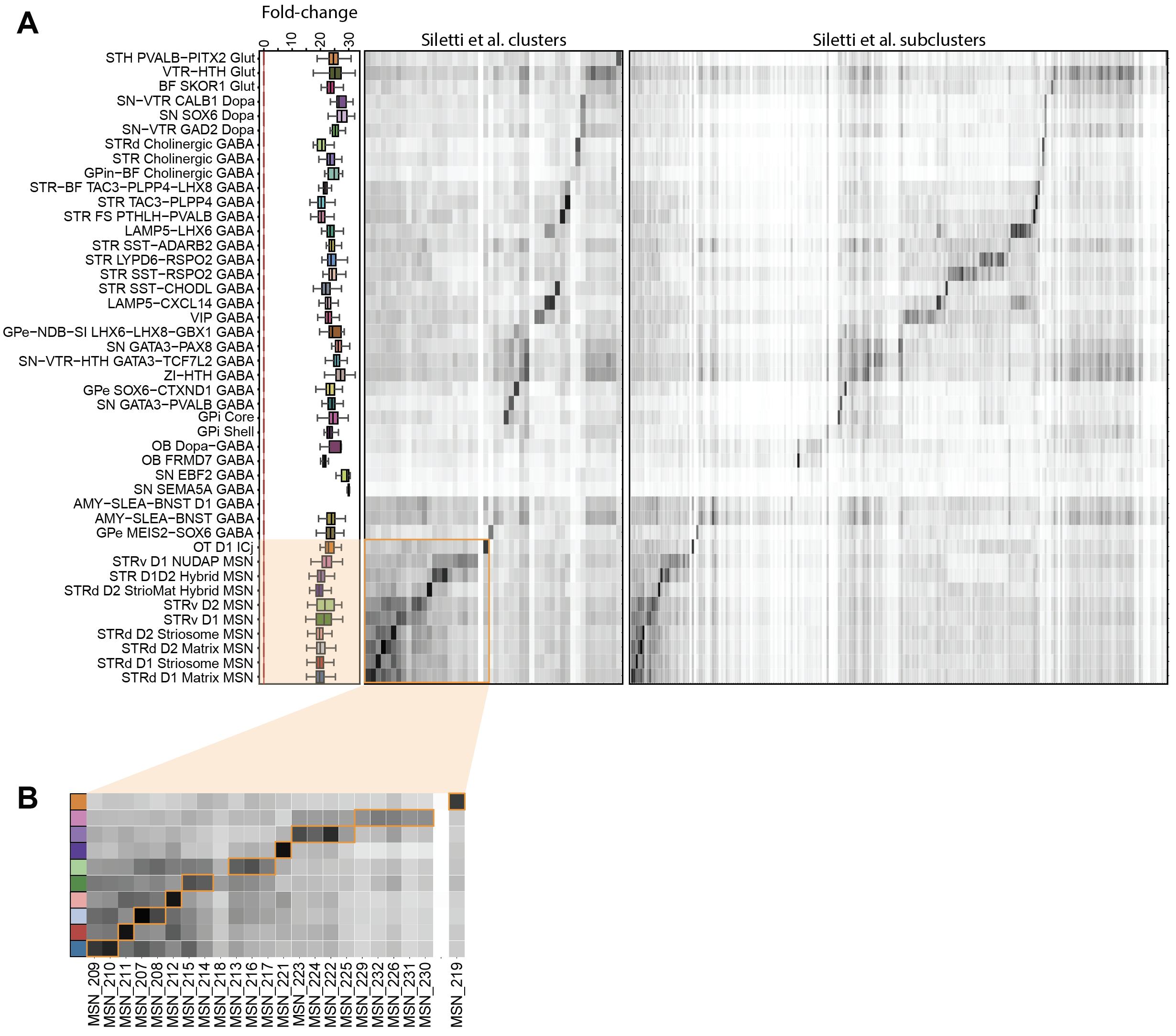

Figure S3.2. BG taxonomy alignment to a human whole brain atlas, related to Figure 3.

(**A**) Comparison of single nucleus sampling and cell type homologies in this study versus Siletti et al. 2023. Fold change indicates the increase in sampling depth based on a Milo analysis of the integrated data sets.

(**B**) Human whole brain clusters that correspond to MSN groups in this study.

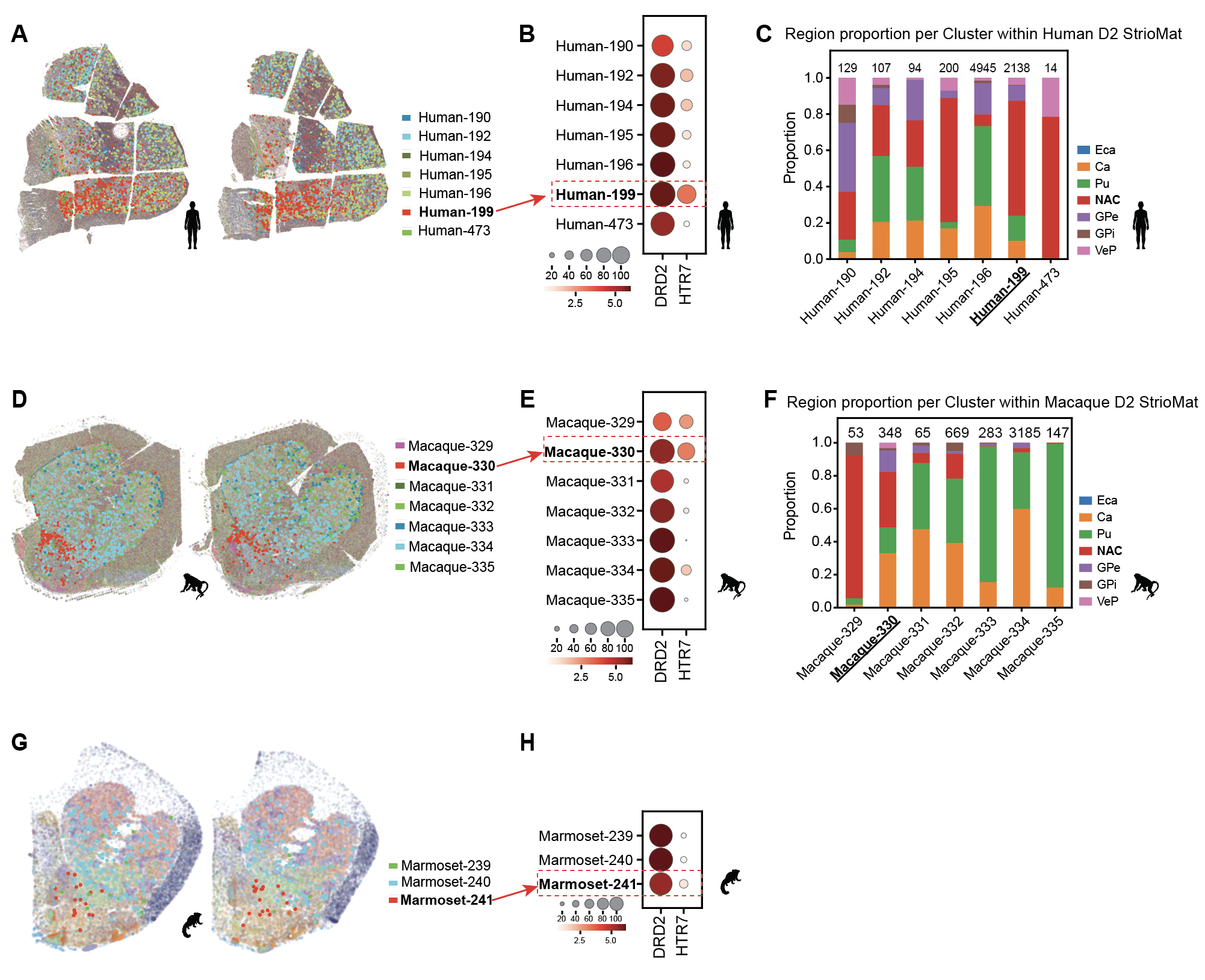
**Figure S3.3. Dorsal-ventral differences in the D2 StrioMat Hybrid MSN Group, related to Figure 3.**

(**A**) Clusters within the D2 StrioMat Hybrid MSN population display distinct spatial transcriptomic patterns in human tissue.

(**B)** The Human-199 cluster has higher expression of *HTR7* based on snRNA-seq data.

(**C**) Regional composition of clusters within the human D2 StrioMat Hybrid MSN Group based on dissections used to generate the snRNA-seq data.

(**D**) Clusters within the D2 StrioMat Hybrid MSN population display distinct spatial transcriptomic patterns in macaque tissue.

(**E)** The Macaque-330 cluster has higher expression of *HTR7* based on snRNA-seq data.

(**F**) Regional composition of clusters within the macaque D2 StrioMat Hybrid MSN Group based on dissections used to generate the snRNA-seq data.

(**G**) Clusters within the D2 StrioMat Hybrid MSN population display distinct spatial transcriptomic patterns in marmoset tissue.

(**H)** The Marmoset-241 cluster has higher expression of *HTR7* based on snRNA-seq data.

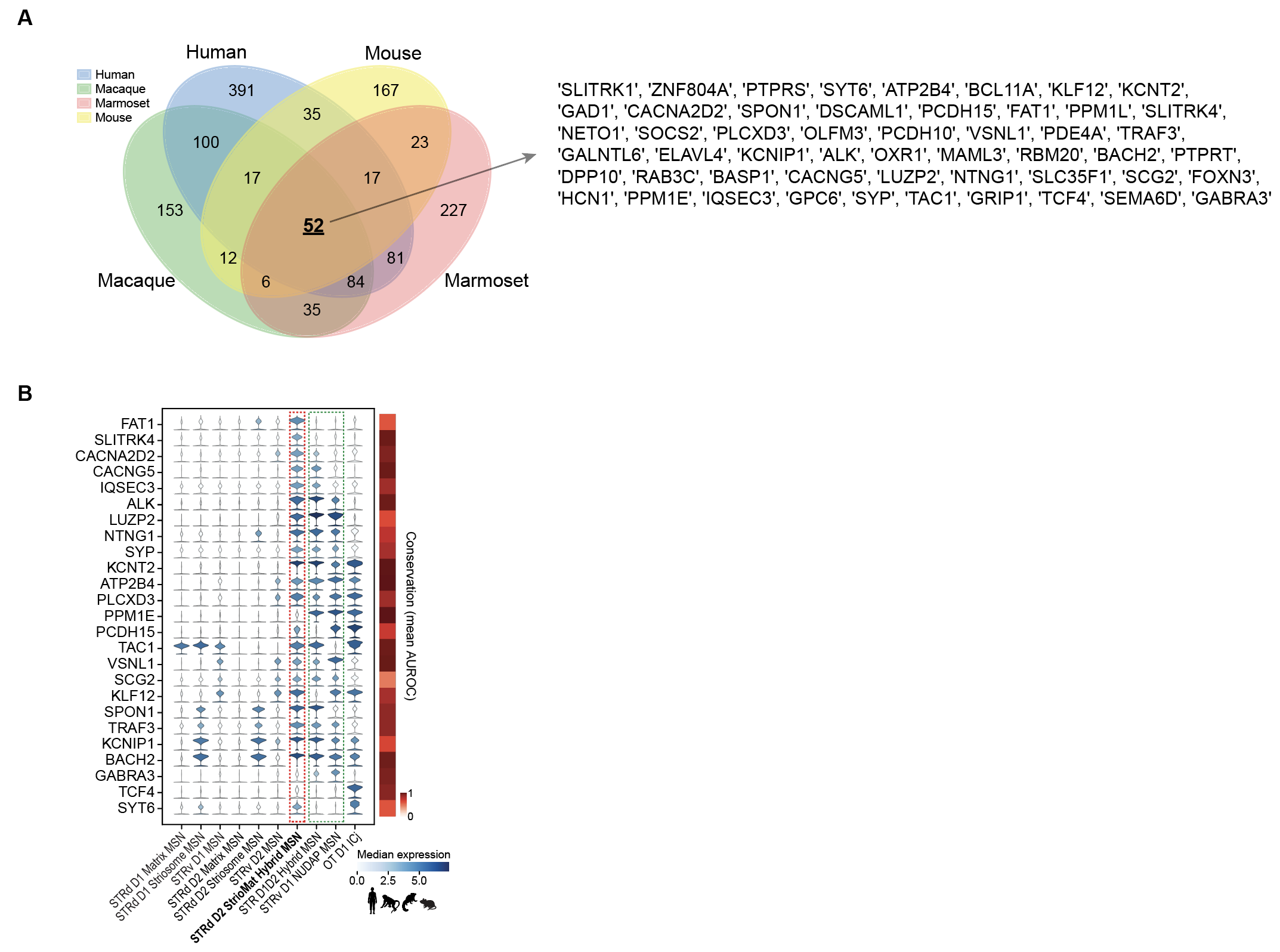

**Figure S3.4. Conserved markers of STRd D2 StrioMat Hybrid in primates and mouse, related to Figure 3.**

(**A**) Venn diagram showing the intersection of species DEGs that are significantly higher in the STRd D2 StrioMat Hybrid MSN Group as compared to other D2 MSNs.

(**B**) Violin plot showing the subset of genes in (A) that are not ubiquitously expressed across D2 MSN Groups. Note that these genes are broadly expressed in eccentric MSNs.

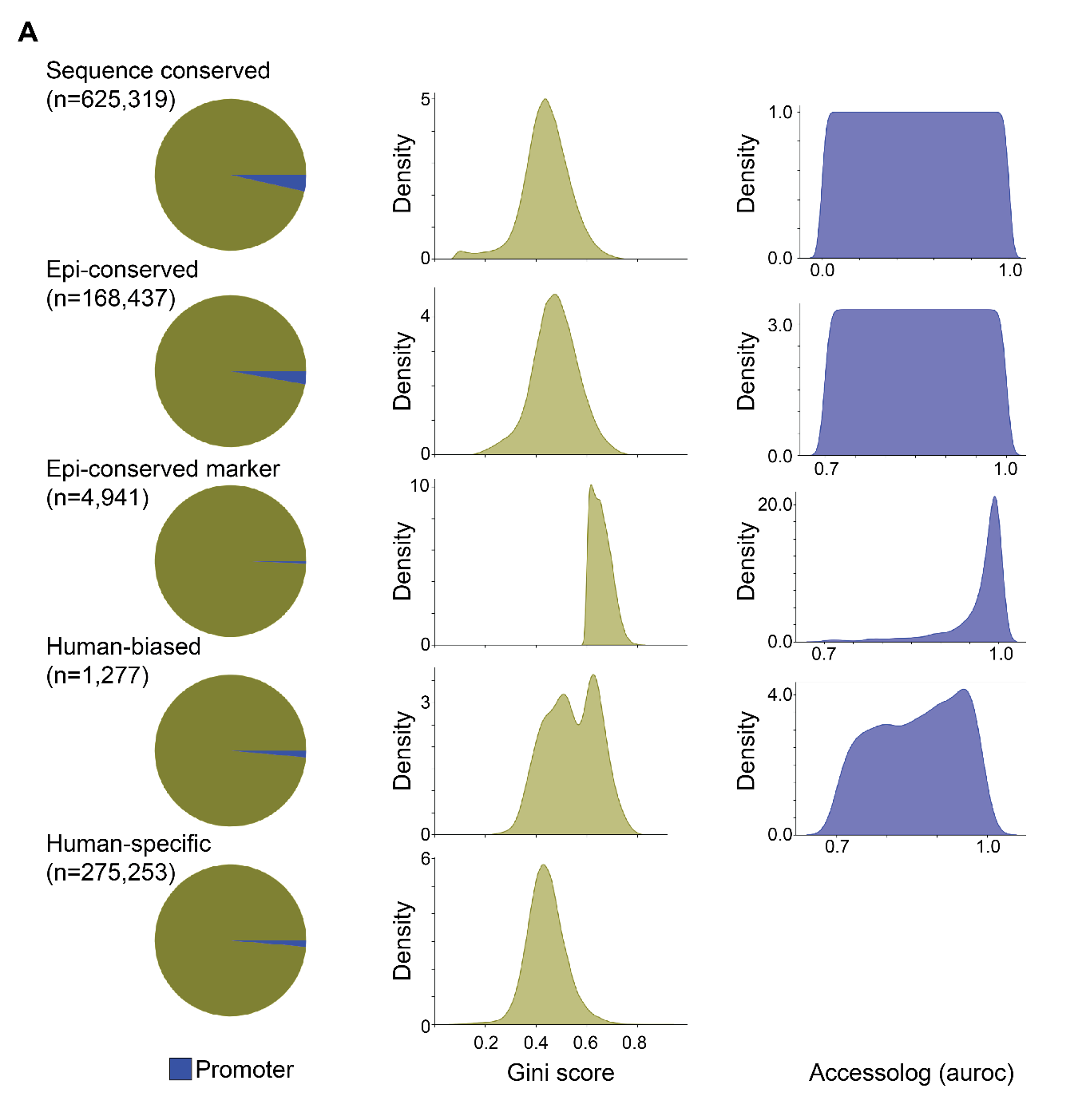

**Figure S4.1. cCRE conservation category annotations, related to Figure 4.**

Left: Pie chart of promoter fractions within each conservation category for cCREs. Middle: Distribution of cell Goup specificity (Gini scores) within each conservation category for cCREs. Right: Distribution of chromatin accessibility conservation across cell Groups (accessolog scores) within each relevant conservation category for CREs.

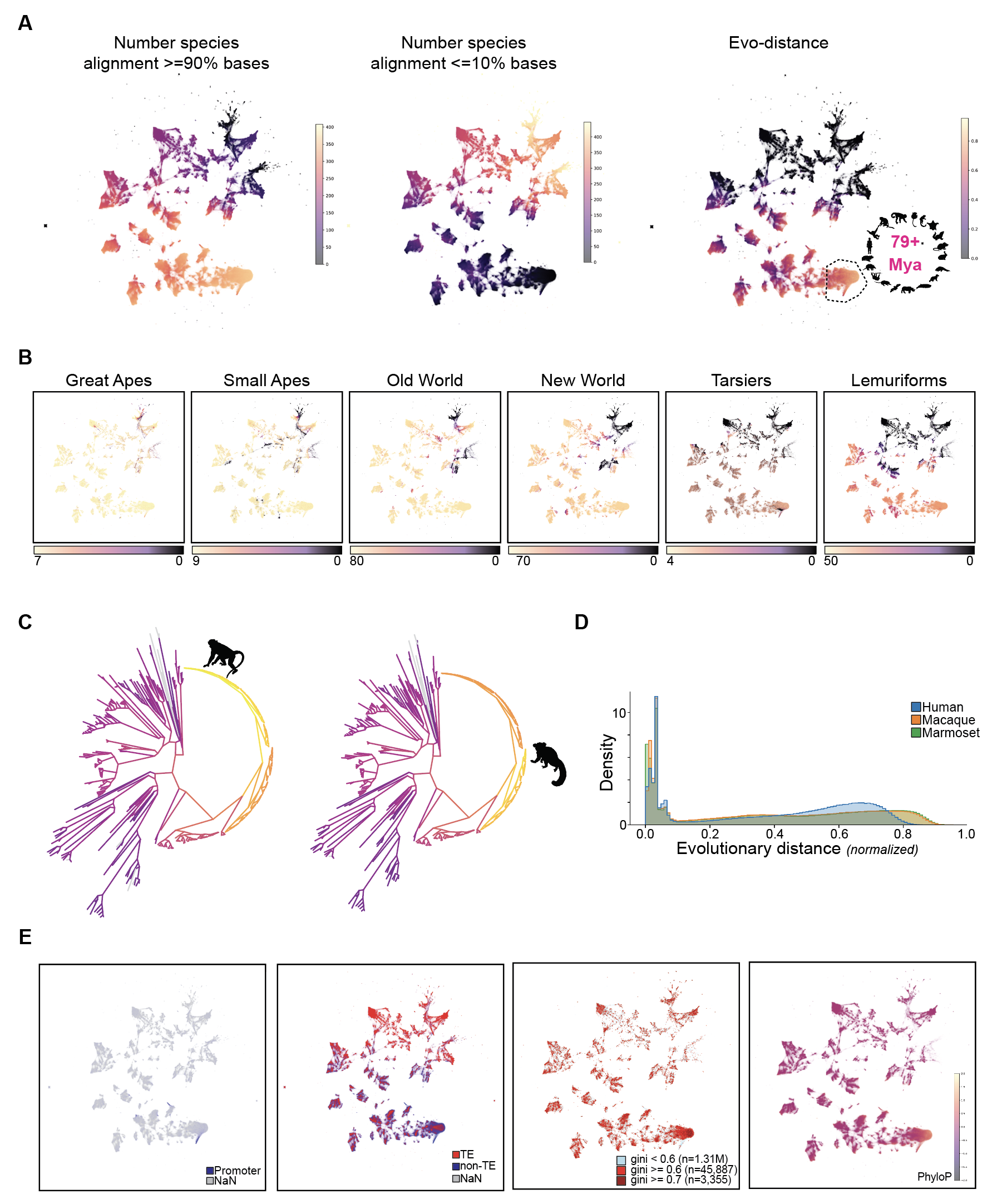

**Figure S4.2. cCRE conservation metrics, related to Figure 4.**

(**A**) UMAP projection of human cCREs (1.3M) based on the count of positions aligning to the 447 mammalian genomes. Point color represents the number of species which each cCRE that aligns within the given threshold (left and middle plots) or evolutionary distance (right plot), a measure of conservation (Methods).

(**B**) UMAP projection of human cCREs colored by the number of species within each primate clade to which each cCRE aligns >= 90% bases.

(**C**) Macaque and marmoset cCRE conservation across mammals and primates. Percent of species-cCREs aligning to each node in the 447 species phylogeny.

(**D**) Distribution of cCRE evolutionary distance metric, normalized per-species.

(**E**) UMAP projection of human cCREs colored by annotations: promoter, TE, cell Group specificity (Gini score), and sequence conservation (PhyloP).

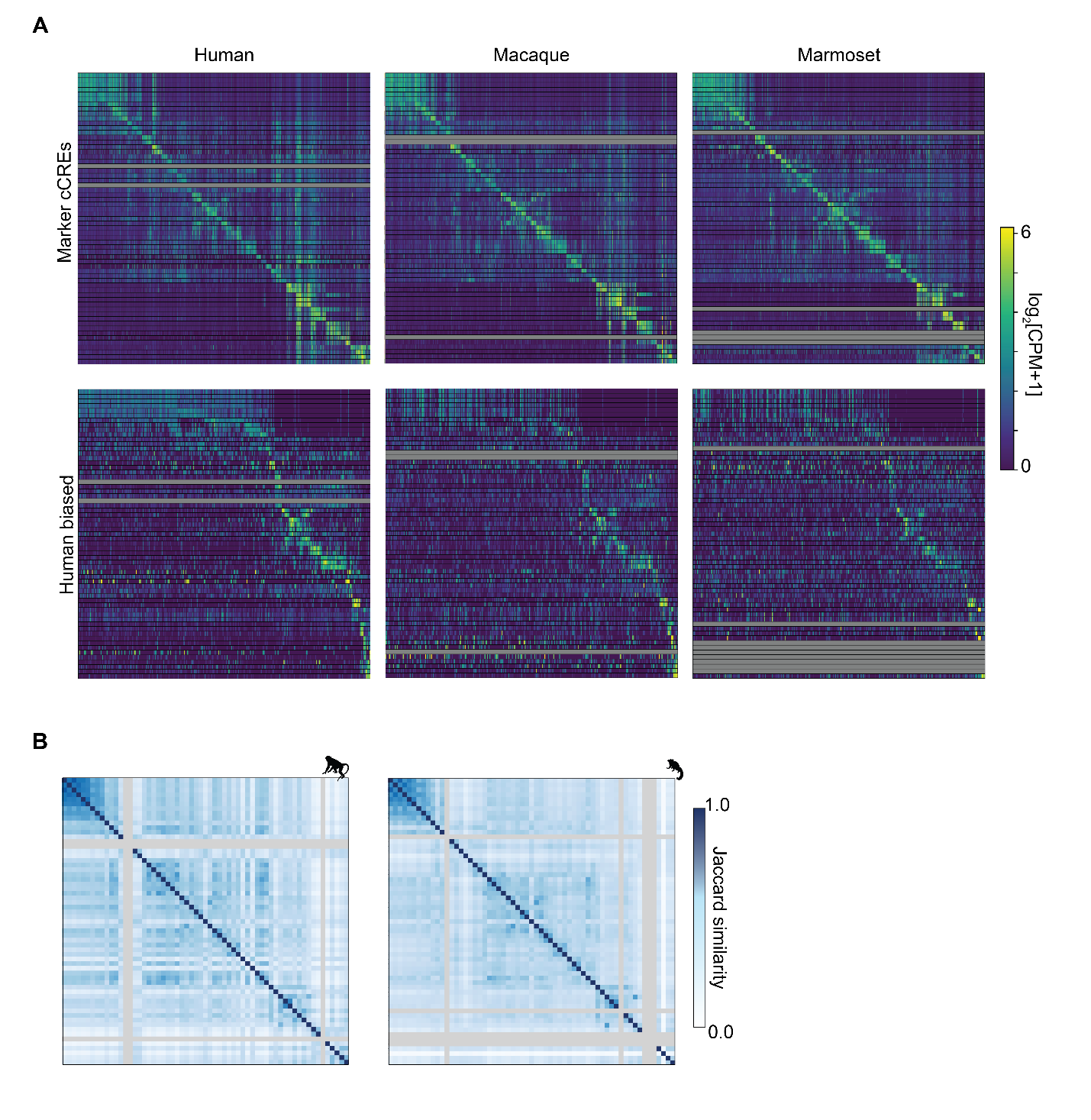

**Figure S4.3. Chromatin accessibility markers and patterns across Groups and species, related to Figure 4.**

(**A**) Heatmap of Group-specific cCREs identified within each species. Heatmap of human-biased cCREs accessibility across species.

(**B**) Jaccard similarity of cCRE accessibility across Groups in macaque and marmoset.

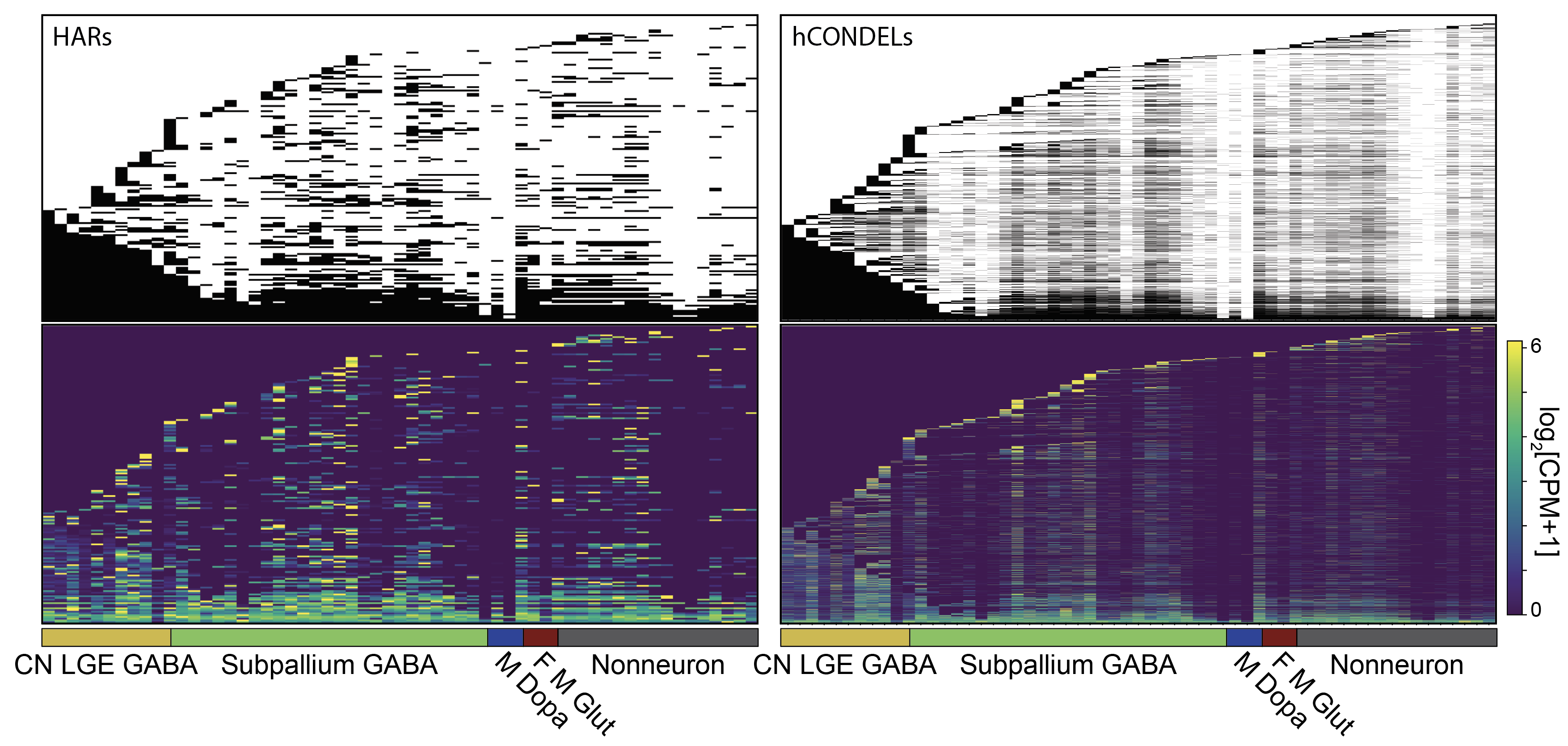

Figure S4.4. HAR and hCONDEL intersection with human cCREs, related to Figure 4.

Human cCREs (rows) that intersect with rapidly evolving genomic regions, including HARs and hCONDELs. Groups (columns) are ordered as in Figure 1 and annotated with cell class.

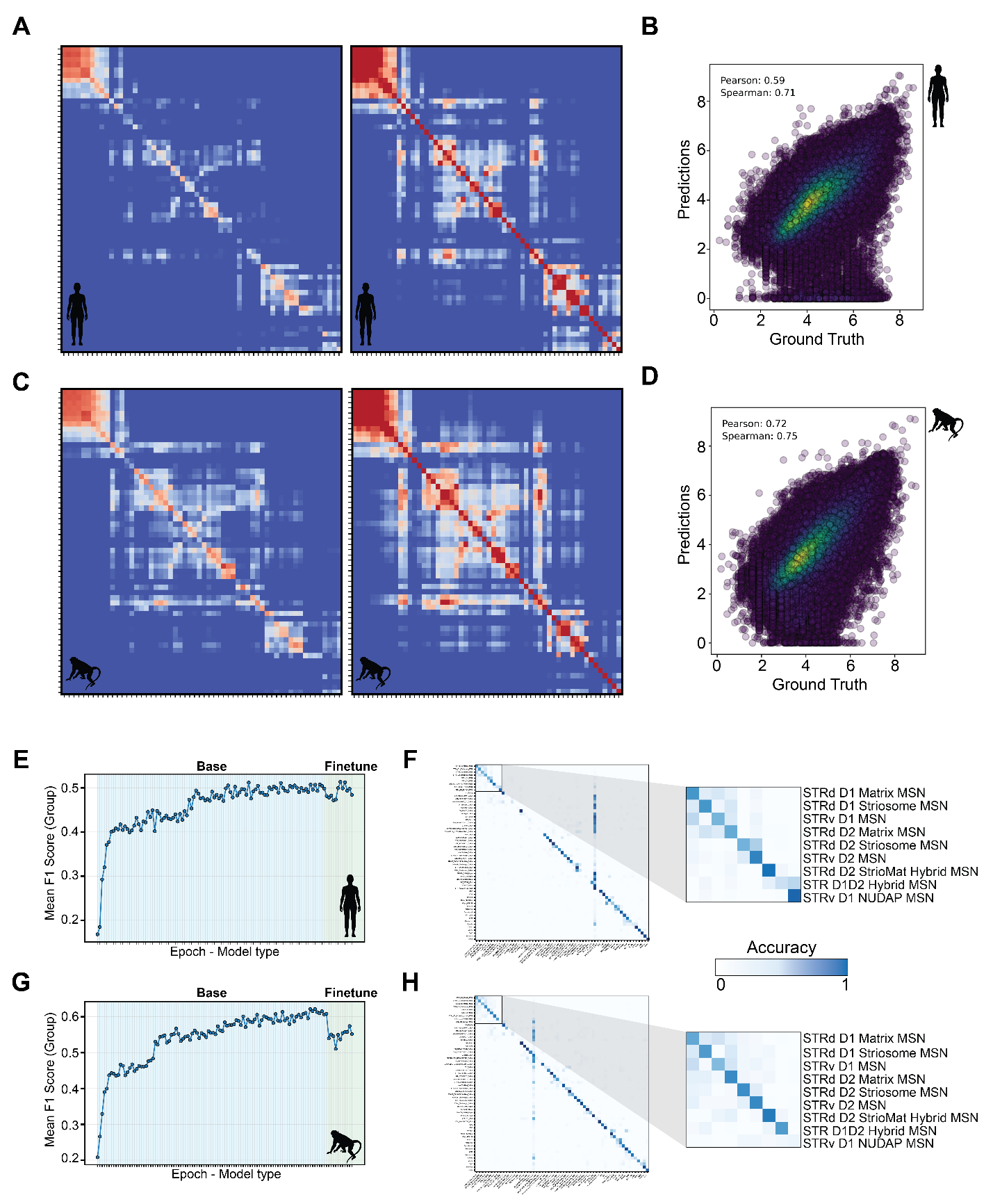

Figure S5.1. Evaluation of deep learning models for DNA sequence across species, related to Figure 5.

(**A**) Heatmap of Pearson correlations for predictions and ground truth (left) and self-correlations of ground truth (right) accessibility patterns for DeepHumanBG.

(**B**) Scatterplot of cCRE accessibility magnitudes as predicted by DeepHumanBG (y-axis) and measured from snATAC-seq (x-axis). Annotated with Pearson and Spearman correlations.

(**C**) Heatmap of Pearson correlations for predictions and ground truth (left) and self-correlations of ground truth (right) accessibility patterns for DeepMacaqueBG.

(**D**) Scatterplot of cCRE accessibility magnitudes as predicted by DeepMacaqueBG (y-axis) and measured from snATAC-seq (x-axis). Annotated with Pearson and Spearman correlations.

(**E**) Classification accuracy (F1 score) for top 100 cCREs per Group based on model embedding at each epoch for DeepHumanBG.

(**F**) Confusion matrix of DeepHumanBG embedding classifier predictions (y-axis) as compared to ground truth labels (x-axis).

(**G**) Classification accuracy (F1 score) for top 100 cCREs per Group based on model embedding at each epoch for DeepMacaqueBG.

(**H**) Confusion matrix of DeepMacaqueBG embedding classifier predictions (y-axis) as compared to ground truth labels (x-axis).

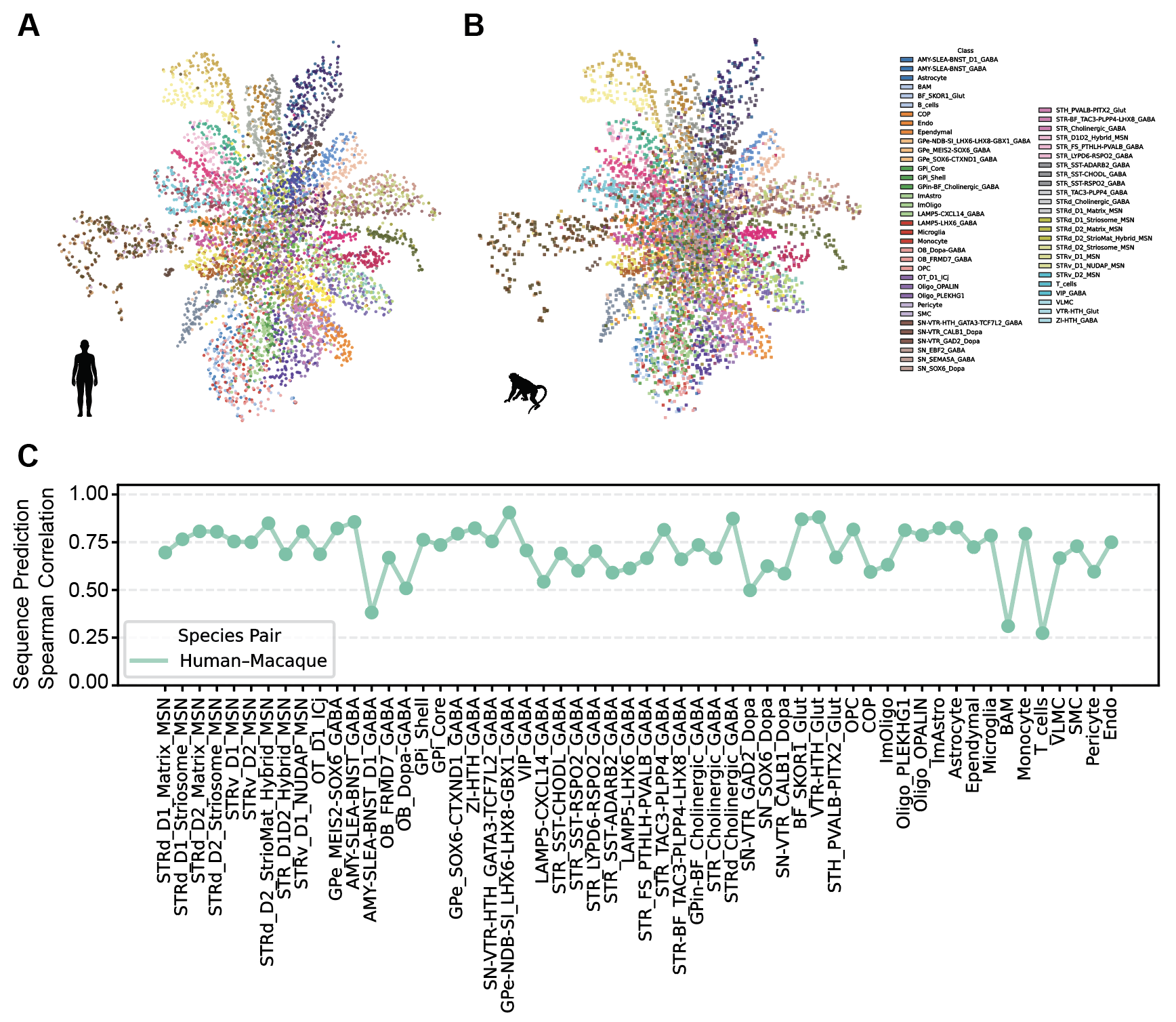

Figure S5.2. Comparative analysis of DNA-sequence underlying cCREs, related to Figure 5.

(**A**) Joint embedding of top 100 cCREs from human and macaque snATAC-seq with DeepHumanBG colored by Group assignment.

(**B**) Spearman correlation of DeepHumanBG predictions between human and macaque cCREs for each Group.

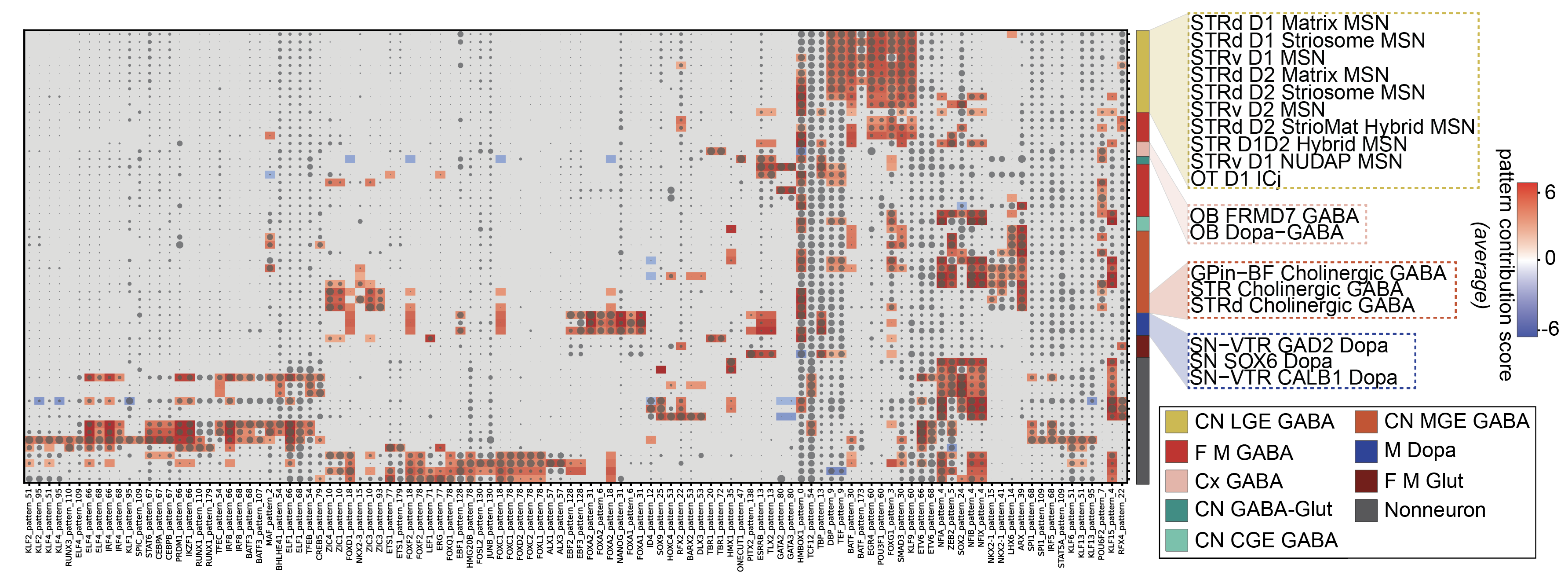

Figure S5.3. TF expression and DNA-sequence importance from Groups from DeepMacaqueBG, related to Figure 5.

(**A**) Cluster map of scaled mean log-normalized TF expression over Groups, indicated by dot size. Sequence patterns from DeepMacaqueBG matched to known TF motifs (columns) with mean importance per Group indicated by color.

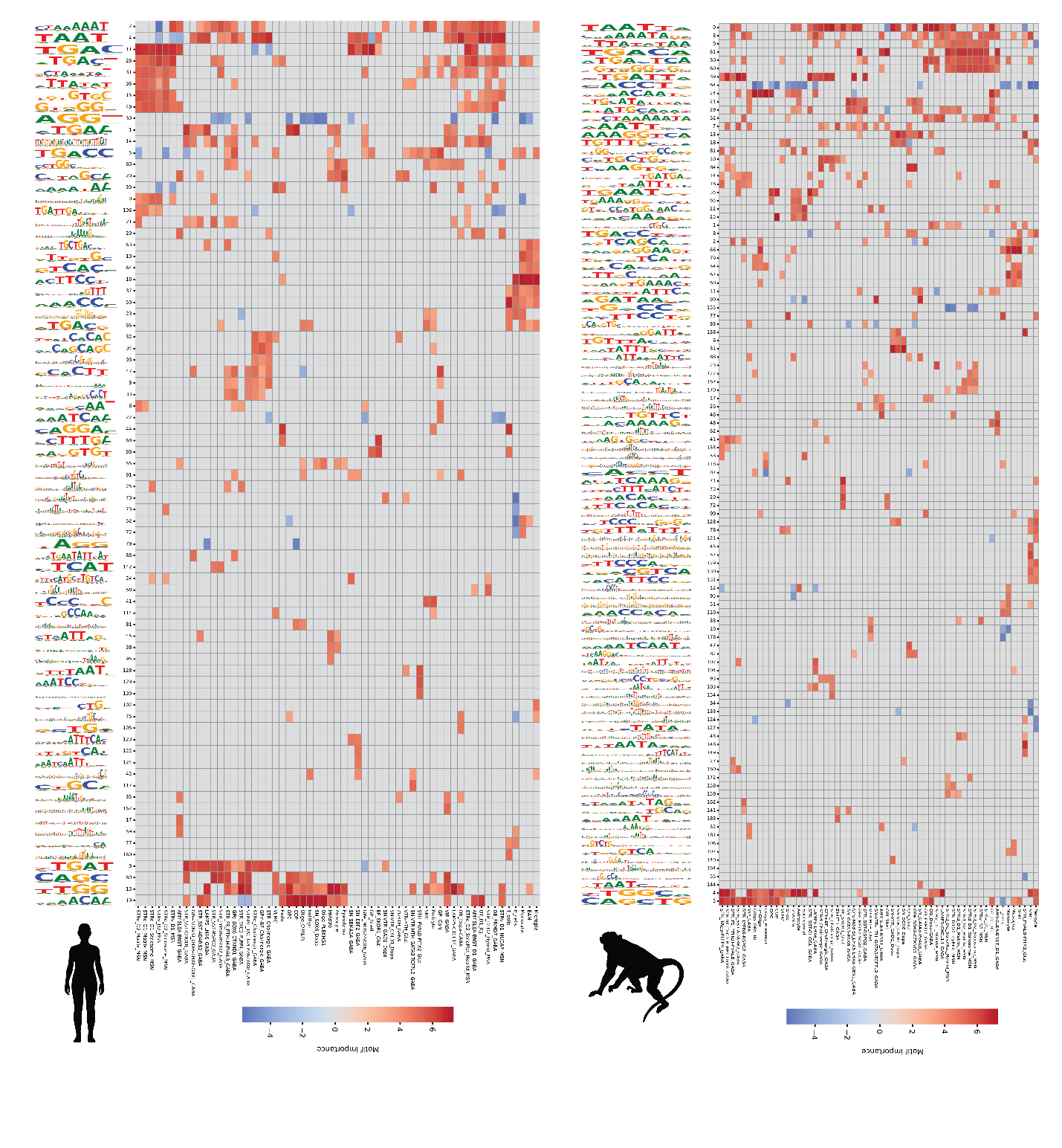

Figure S5.4. DNA motif associations with Groups in human and macaque, related to Figure 5.

(**A**) Heatmap of pattern (motif) importances for each Group from DeepHumanBG colored by pattern importance.

(**B**) Heatmap of pattern (motif) importances for each Group from DeepMacaqueBG colored by pattern importance.

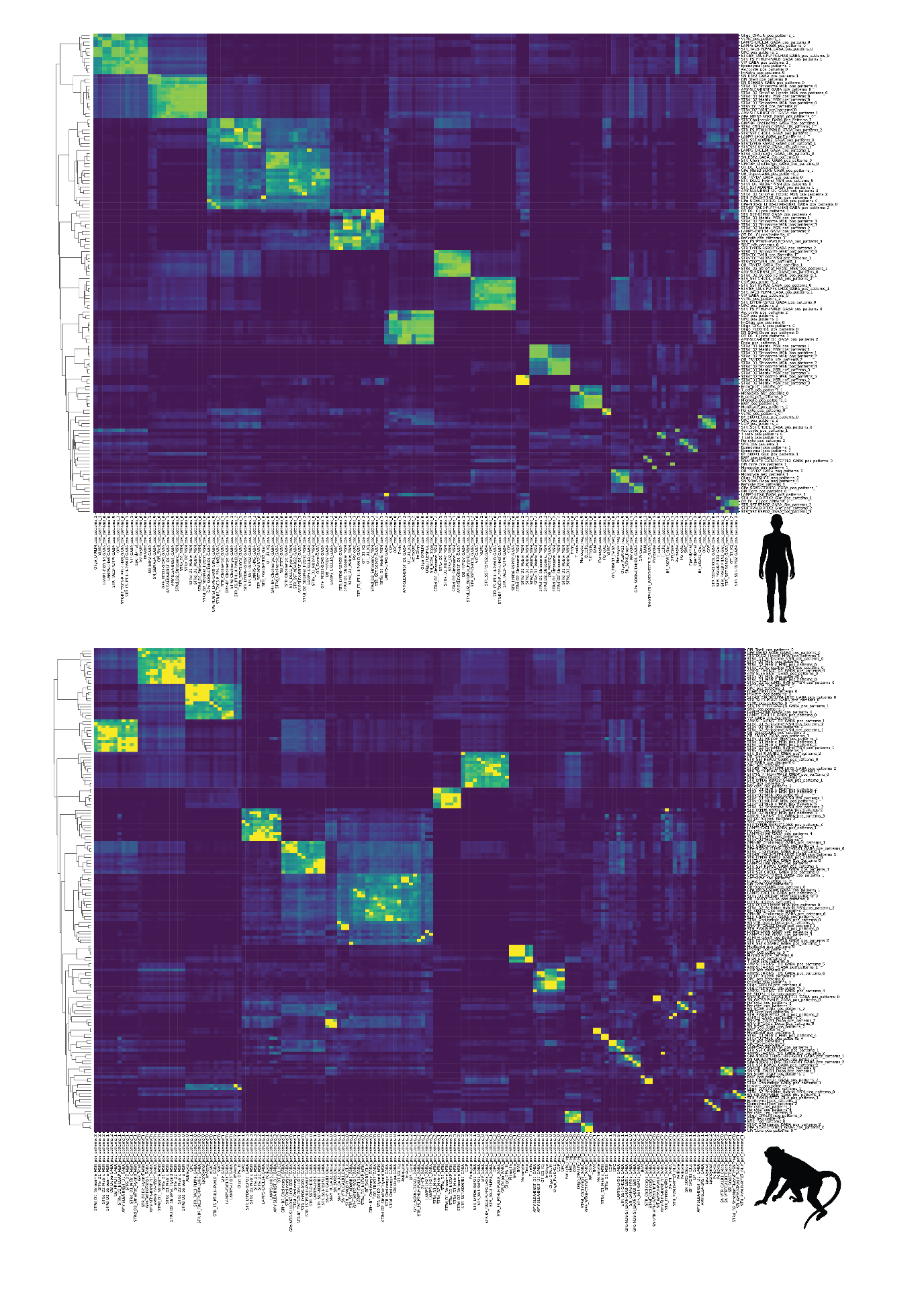

Figure S5.5. Relatedness of DNA motifs between Groups, related to Figure 5.

(**A**) Heatmap of similarities for DNA motifs between Groups from DeepHumanBG.

(**B**) Heatmap of similarities for DNA motifs between Groups from DeepMacaqueBG.

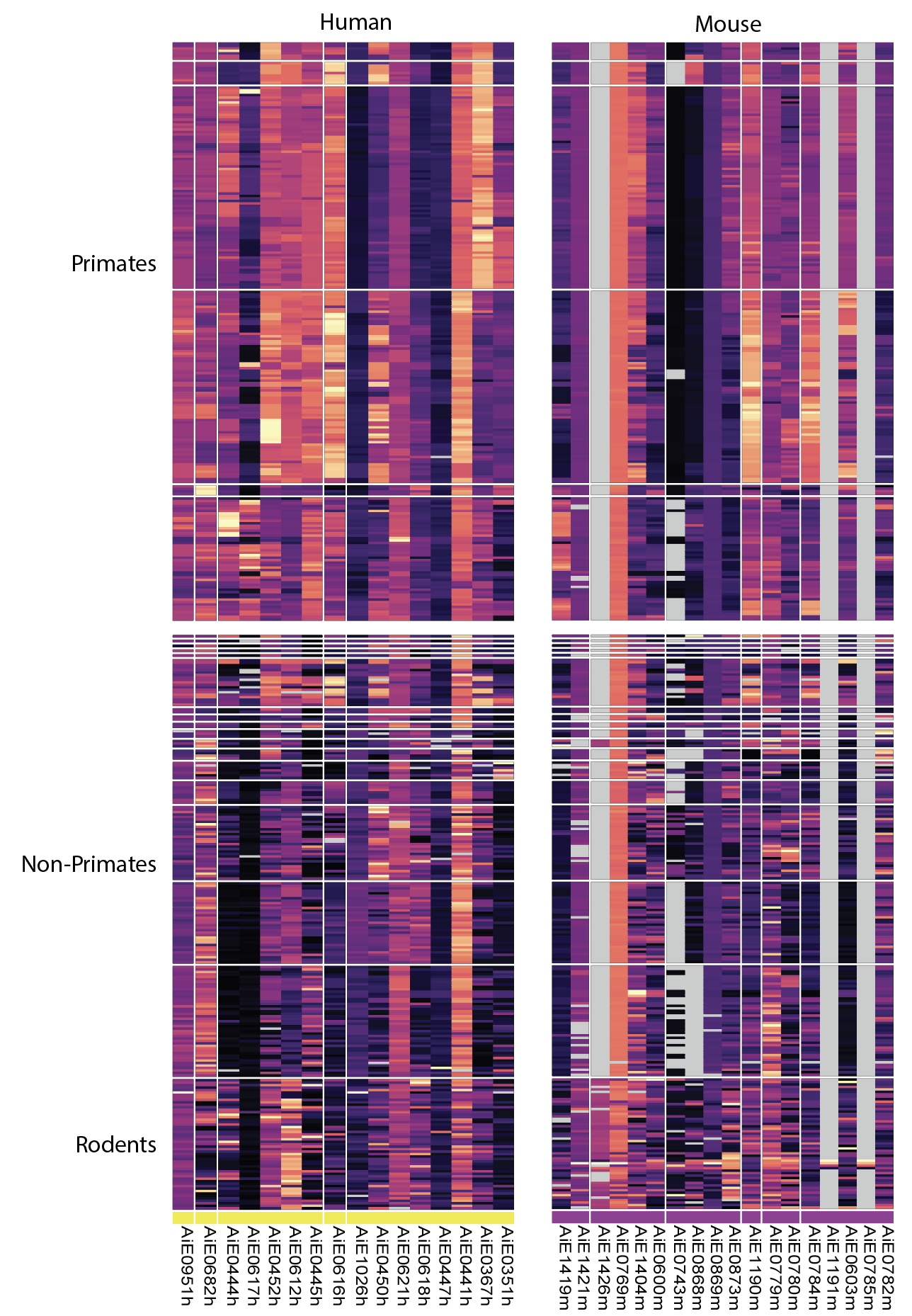

Figure S5.6. Comparative analysis of striatal tool collection across 447 species, related to Figure 5.

Predicted human or mouse enhancer activity in the targeted striatal cell type using the sequence model to predict importances of the orthologous sequence for each species. Lighter colors correspond to higher predicted enhancer activity, and grey indicates that no orthologous sequence was identified in a species based on the CACTUS alignment.

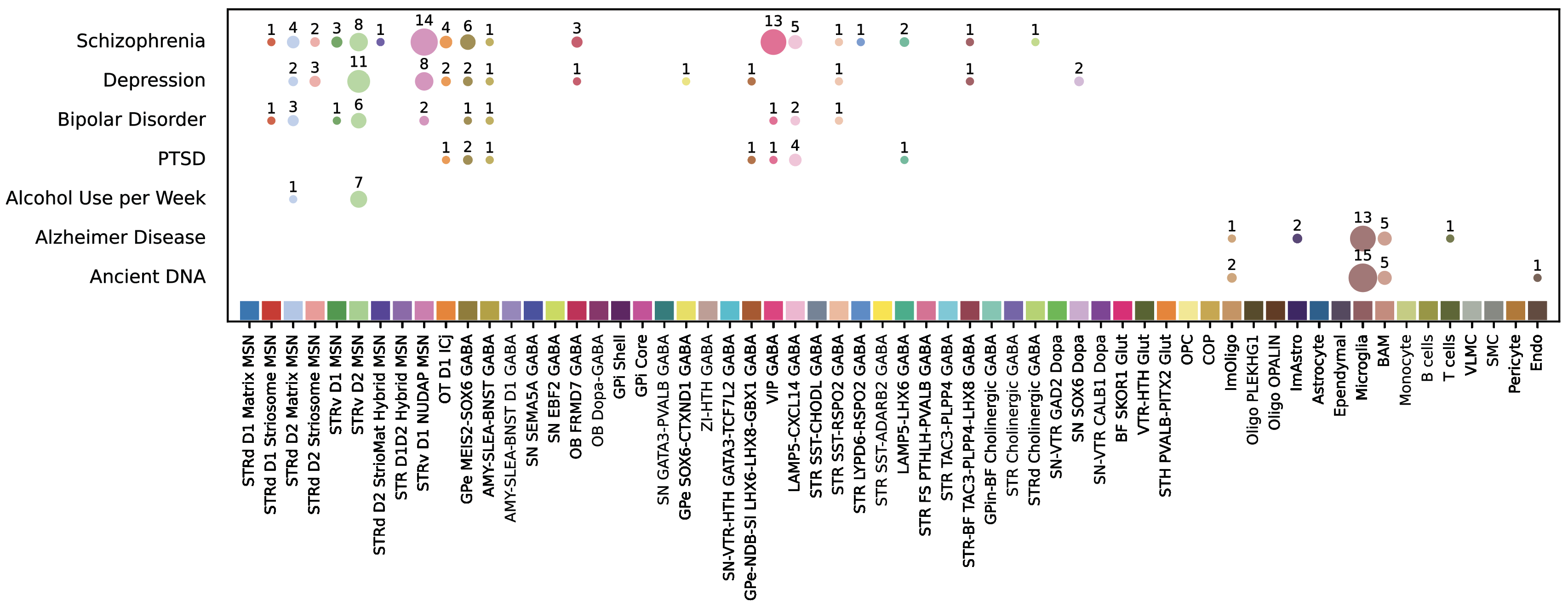

Figure S6.1. Disease associations with the complete BG taxonomy, related to Figure 6.

Phenotype associations with BG groups obtained using MAGMA. Dot size and labels indicate the number of significant (P < 0.0001) clusters per group.

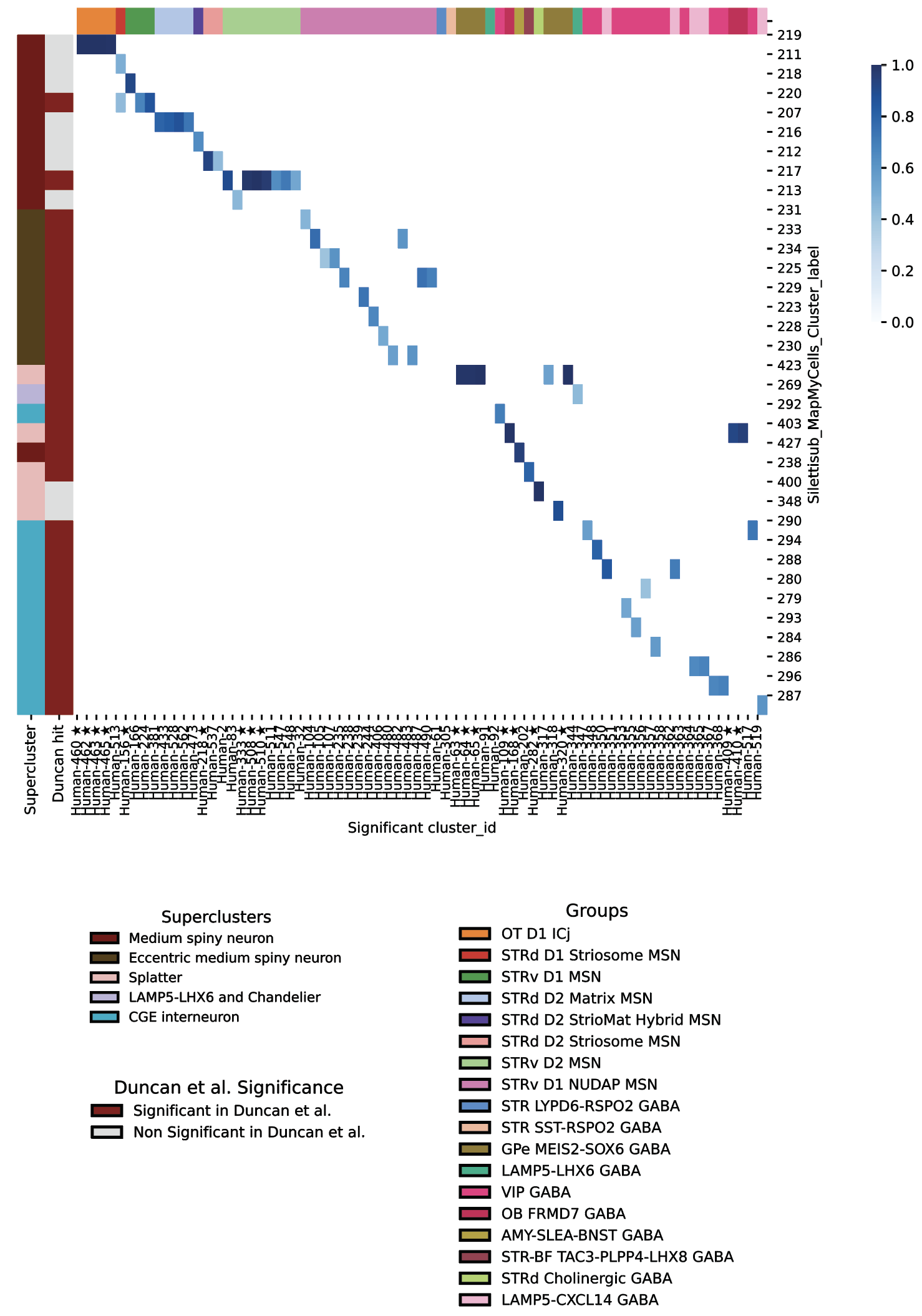

Figure S6.2. Correspondence of BG cell type associations with SCZ to previous work, related to Figure 6.

Confusion matrix comparing BG clusters (X-axis) that were significantly associated (MAGMA; P < 0.0001) with SCZ with clusters reported in the whole human brain by Siletti et al. 2023 (Y-axis). Clusters with purity > 0.9 are marked with a star, and mappings with values < 0.4 are excluded. Column colors represent BG taxonomy groups, while row colors represent superclusters from Siletti et al. The second vertical color bar indicates whether the corresponding cluster in Siletti’s taxonomy was associated with SCZ (dark red) by Duncan et al. 2025.

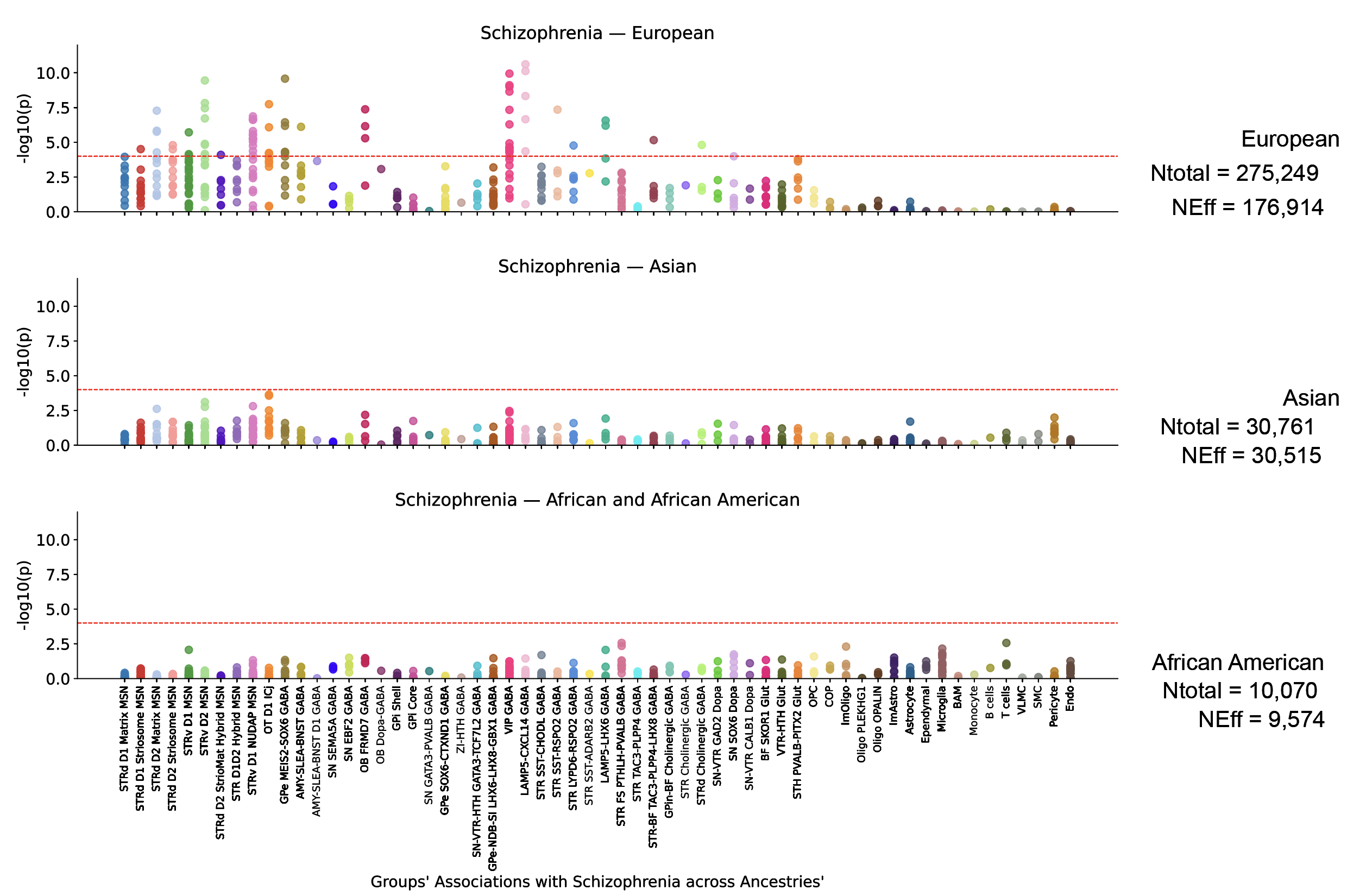

Figure S6.3. More cell types associated with SCZ using larger GWAS, related to Figure 6.

Detailed associations (-log10 p-values) of clusters with SCZ across ancestries (European, Asian, African-American) compared to the number of individuals per cohort and the corresponding effective population size (Neff). Points are color-coded by group.

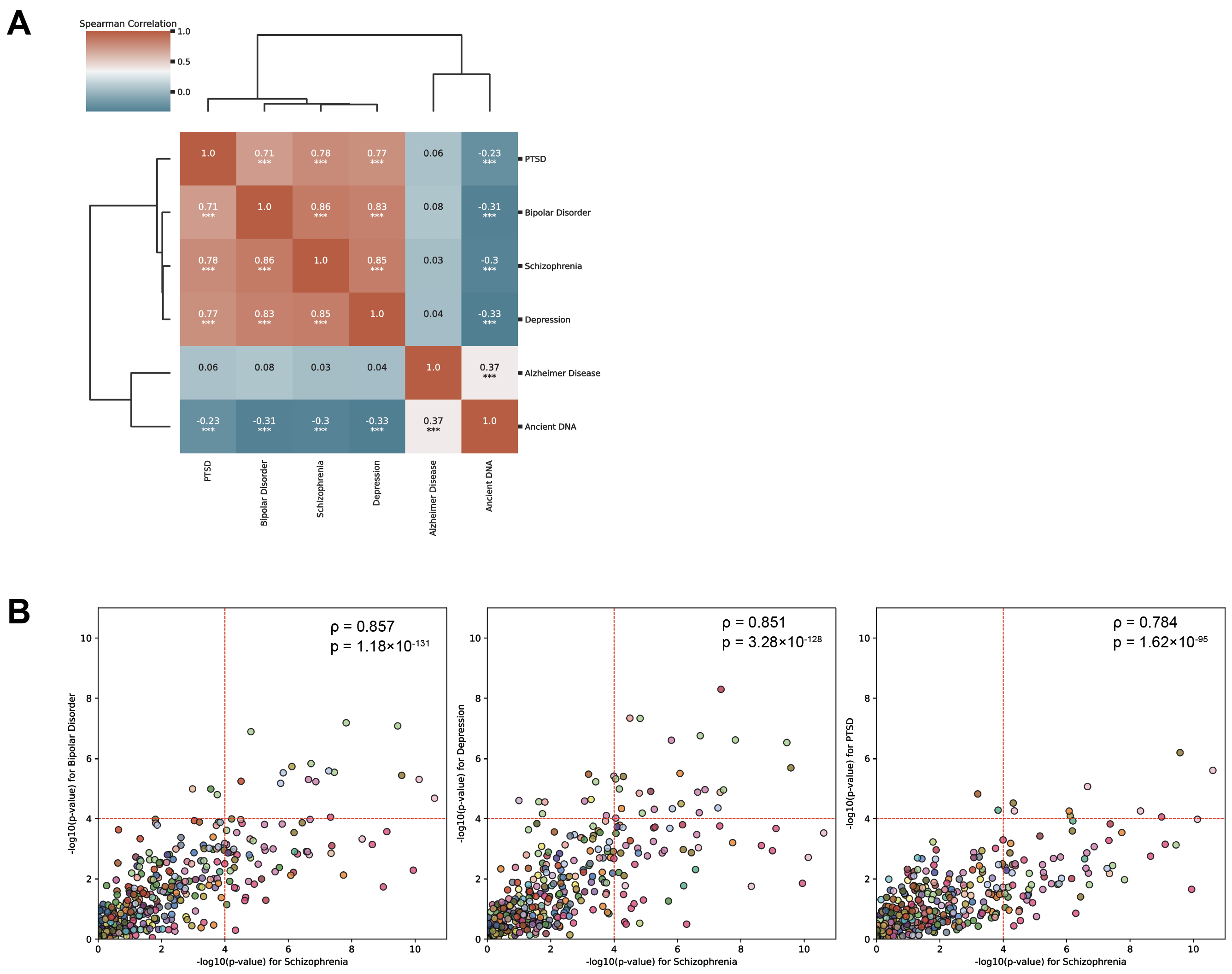

Figure S6.4. Disease-specific associations with BG cell clusters, related to Figure 6.

(**A**) Spearman correlations of phenotypes based on the significance of cluster associations.

(**B**) The scatter plots show -log₁₀ p-values for SCZ versus those for other psychiatric conditions. Significant positive Spearman correlations were observed: bipolar disorder (ρ = 0.857, *p* = 1.18×10⁻¹³¹), depression (ρ = 0.851, *p* = 3.28×10⁻¹²⁸), and PTSD (ρ = 0.784, *p* = 1.62×10⁻⁹⁵). Points are color-coded by group.
